# Alternative splicing is coupled to gene expression in a subset of variably expressed genes

**DOI:** 10.1101/2023.06.13.544742

**Authors:** Guy Karlebach, Robin Steinhaus, Daniel Danis, Maeva Devoucoux, Olga Anczuków, Gloria Sheynkman, Dominik Seelow, Peter N Robinson

## Abstract

Numerous factors regulate alternative splicing of human genes at a co-transcriptional level. However, how alternative splicing depends on the regulation of gene expression is poorly understood. We leveraged data from the Genotype-Tissue Expression (GTEx) project to show a significant association of gene expression and splicing for 6874 (4.9%) of 141,043 exons in 1106 (13.3%) of 8314 genes with substantially variable expression in ten GTEx tissues. About half of these exons demonstrate higher inclusion with higher gene expression, and half demonstrate higher exclusion, with the observed direction of coupling being highly consistent across different tissues and in external datasets. The exons differ with respect to sequence characteristics, enriched sequence motifs, RNA polymerase II binding, and inferred transcription rate of downstream introns. The exons were enriched for hundreds of isoform-specific Gene Ontology annotations, suggesting that the coupling of expression and alternative splicing described here may provide an important gene regulatory mechanism that might be used in a variety of biological contexts. In particular, higher inclusion exons could play an important role during cell division.

## INTRODUCTION

Over 95% of human multi-exon genes undergo alternative splicing (AS) in a developmental, tissue-specific, or signal transduction-dependent manner (1). Splicing is a highly regulated process by which an intron is excised from a pre-mRNA transcript and the flanking exons are ligated together by a series of steps, whereby all or part of the splicing process occurs co-transcriptionally (2–4). Transcript elongation follows the initiation of transcription, adding ribonucleoside triphosphates to the growing mRNA chain. Splicing, as well as other processes involved in mRNA maturation is influenced by interactions with the RNA polymerase II (RNAP2) transcript elongation complex (2). Changes in promoter sequence and occupation can modify the splicing pattern of several genes, evidencing a coupling between transcription and alternative splicing (5–8). It has been proposed that the promoter effect involves modulation of RNA polymerase II elongation rates (9, 10). Two major and potentially complementary models have been proposed to explain how transcription and splicing are coupled, referred to as the kinetic coupling and the spatial coupling models.

Kinetic coupling refers to the notion that the rate of transcription elongation determines the temporal “window of opportunity” for selection or rejection of an upstream sequence. If upstream and downstream events on the nascent transcript compete, the upstream sequence will have a “head start” because it emerges from RNAP2 before the downstream sequence does. The advantage conferred by the head start is greater when elongation is slow (10, 11). It has been shown that elongation rate can influence AS by modulating several classes of co-transcriptional events including alternative splice site recognition, binding of regulatory proteins, and formation of RNA secondary structures (12, 13). These observations led to the notion that slow elongation expands the “window of opportunity” for recognition of an upstream 3’ splice site before it must compete with a downstream site, therefore promoting inclusion of the upstream cassette exon. In contrast, slow elongation was shown to favor promoter skipping of CFTR exon 9 by increasing the recruitment of the negative factor ETR-3 onto the UG-repeat at the 3’ splice site of the exon (14, 15).

Spatial coupling refers to the ability of the transcription machinery to recruit various classes of RNA processing factors to the site of transcript. The RNAP2 C-terminal domain (CTD) plays a central role in recruiting factors involved in transcriptional elongation, splicing, and other functions related to mRNA maturation. The RNAP2 CTD is extensively phosphorylated and dephosphorylated upon different stages of transcription and acts as a dynamic docking site for factors required for the mRNA processing events that occur together with transcript elongation (16). Transcribed exons are tethered to the elongating RNAP2 transcription complex (17, 18). The serine and arginine-rich splicing factor 3 (SRSF3) was shown to possess a CTD-dependent inhibitory action on the inclusion of fibronectin cassette exon 33 (19).

Numerous other factors influence AS, including nucleosome occupancy, chromatin remodelers, RNA secondary structure, as well as histone marks and DNA methylation and the protein factors that interact with them (20–24). In principle these factors could influence AS by modulating elongation through differential nucleosome density, histone modification profiles, DNA methylation density, or by recruiting splicing factors to the chromatin template as the transcriptional machinery passes (11, 25). Two studies have demonstrated a pervasive impact of elongation rate on splicing. The first showed that reduction of RNAP2 elongation speed by drugs or RNAP2 mutations tended to increase exon inclusion levels (26). Interestingly, many of the corresponding splicing events often introduce premature truncation codons (PTCs), which are predicted to lead to nonsense-mediated decay (NMD). This has been shown experimentally to be a common mechanism for gene regulation, including the autoregulation of proteins that affect the splicing process (27–29). A second study investigated RNAP2 mutants that increased or decreased elongation rates, characterizing exons for which a faster elongation rate results in more inclusion of the exon in transcripts, and exons, for which a faster gene expression rate results in more skipping of the exon in transcripts (30).

Although gene expression is controlled by numerous transcriptional and posttranscriptional factors, substantial evidence argues that expression of most genes is controlled in part at the level of transcription elongation (31–36). In this work, we leverage comprehensive bulk RNA-seq data from the Genotype-Tissue Expression (GTEx) project (37, 38) to investigate associations between gene expression and AS. We identify thousands of exons whose inclusion or exclusion is correlated to the overall level of gene expression and characterize significantly different properties of the exons and the transcripts and genes they are contained in.

## MATERIAL AND METHODS

### Data

*RNA-seq data*: The Genotype-Tissue Expression (GTEx) project offers a genome-wide quantification of the expected number of transcripts in thousands of samples across tens of different human tissues (37). Quantification is performed using bulk RNA-Sequencing and the RSEM tool (39). We used the file GTEx_Analysis_2017-06-05_v8_RSEMv1.3.0_transcript_tpm.gct.gz, which provides transcripts per million counts across tissues such that expression levels are normalized across experiments.

The tissues we tested include Lung, Spleen, Thyroid, Brain - Cortex, Adrenal Gland, Breast - Mammary Tissue, Heart - Left Ventricle, Liver, Pituitary, and Pancreas.

We repeated our analysis of type 0, UHP, and DHP exons in three breast (SRP301453), left ventricle (SRP237337), and liver (SRP326468) bulk RNA-seq datasets that were obtained from the Sequence Read Archive (SRA) (40).

Gene models, which were used for the definition of exon bounds and transcript affiliation, were derived from the GTF file Homo_sapiens.GRCh38.91.gtf from GENCODE (41). The GTF file contained 683,196 unique exons.

*ChIP-seq data*: For ChIP-Seq peaks, we downloaded BED files from ENCODE using the provided filters to select ChIP-Seq files for POLR2A in human cells (42). This resulted in 105 BED files containing peaks. File names are provided in Supplemental Table S6.

### Gene expression variability threshold

We reasoned that genes that do not display a certain minimum level of expression variability would not be highly powered to discover associations of expression with alternative splicing. Therefore, we applied the following inclusion criteria. The GRCh38 GENCODE annotations of the human genome comprise 683,196 exons. Exons were removed from further analysis unless they were expressed in at least half of the samples from a given tissue (i.e., had a read count of at least one) and which displayed a mean expression level across all samples from the tissue of 20 counts or more. Additionally, we calculated the ratio of the 95^th^ percentile and 5^th^ percentile of the expression values, and removed exons whose ratio was less than 2.0. Finally, we limited analysis to genes that contained at least one exon that showed alternative splicing, defined as a gene with at least two transcripts that differed with respect to inclusion or exclusion of an exon or exon segment.

### Percent Spliced In (ψ)

For each gene that passed that threshold defined in the previous section, we investigated whether the transcripts differ with respect to inclusion or exclusion of a cassette exon. If so, we treat each affected cassette exon in the gene separately, and define the count of transcripts that contain the exon *e*_*i*_ as *IT*^*inc*^ *(e*_*i*_) and the count of transcripts that exclude the exon as *IT*^*excl*^ *(e*_*i*_) to calculate the Percent Spliced In, *ψ(e*_*i*_) as

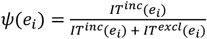

If multiple sets of exons are perfectly correlated with respect to transcript structure, they are collapsed such that the statistics for the event are calculated only once. For instance, if a gene has two transcripts with exon structure A-B-C-D-E and A-C-E, then we calculate the selection criteria for only one of the alternatively spliced exons B and D and apply them to both.

### Correlation between gene expression and alternative splicing

We investigated potential associations between gene expression and alternative splicing of cassette exons as defined above. We applied the following linear regression model for cassette exon *e*_*i*_ of gene *g*, whereby *Y(g)* is the total expression of the gene (sum of counts of all transcripts assigned to the gene), and *ψ(e*_*i*_) is the percent spliced in as defined above.

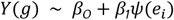

In words, the model predicts the gene expression level based on exon inclusion fraction. The *p*-value for the coefficient *β*_*1*_ tests the null hypothesis that *ψ(e*_*i*_) has no correlation *Y(g)*. This *p*-value is corrected for multiple testing using the Benjamini Hochberg method (43) in each tissue separately.

We conclude that there is a significant relationship between alternative splicing and expression if the corrected *p*-value is 0.05 or less, the coefficient of determination (*R*^*2*^) is at least 0.5, and additionally the ratio of the 95 percentile and 5 percentile of the expression values is at least 2. The results of this analysis are used to define the exon type. For each analyzed cassette exon, if there is a significant correlation and *β*_-_>0, that is, higher inclusion predicts higher expression, the exon is classified as upregulated-high ψ (UHP). If *β*_-_ < 0, that is, higher inclusion predicts lower expression, the exon is classified as downregulated-high ψ (DHP). If the relationship is not significant, the exon is classified as type 0. We note that exons that are not cassette exons are not classified by our definition.

### RNAP2 ChIP-seq

CHIP-seq peaks from the 105 RNAP2 ChIP-Seq experiments (Supplemental Table 2) were obtained from the ENCODE project website. Each file was treated as a separate experiment. For each gene that had type 0 exons and at least one more exon type, the number of bindings per base pair on type 0 and the other exon types were summed over the ChIP-Seq experiments, and the ratio between UHP/DHP and type 0 bindings per base pair were computed.

Exon coordinates of UHP, DHP, and type 0 exons were intersected with the peak coordinates using the bedtools intersect program with default parameters (44). The ratios of positive counts were computed for every gene that contained at least one UHP/DHP exon.

### Analysis of PRO-Seq datasets

We obtained the aligned reads for the dataset of (45) in .bam file format from the ENCODE website using the PRO-Seq filter, which retrieves 8 files corresponding to two biological samples. For the dataset of (46) we obtained the fastq files from SRA and processed them using the pipeline described in (47), using the ‘output-genome-bam’ option of RSEM. In order to compute overlaps with intronic regions we used bedtools intersect with default parameters (44). The ratios of positive counts were computed for every gene that contained at least one UHP/DHP exon.

### Enriched motif testing

Here, we characterized predicting sequence motifs for transcript factor binding sites (TFBS), RNA-binding protein (RBP) binding sites, and core promoter elements (CPE).

We characterized TFBS predicted by detailed transcript factor flexible models (TFFM) (48) in the promoters of genes containing at least one type 0 exon but no UHD or DHP exon (referred to as type 0 gene), genes containing no type 0 or DHP exon but at least one UHP exon (referred to as UHP gene), and genes containing no type 0 or UHP exon but at least one DHP exon (referred to as DHP gene). TFFMs binding motifs were taken from JASPAR (49), RBP matrices were taken from the RNA-binding protein database (50), and CPEs were characterized as previously (51). The calculations were conducted within the backend infrastructure of the FABIAN-variant application (52).

We derived empirical p-values by random sampling (without replacement) with one million permutations of our variable of interest. The p-value is the proportion of samples that have a test statistic larger than that of our observed In our case, the statistic of interest is the difference of the proportion of hits for some protein-binding factor in UHP (or DHP) vs. type 0 exons. For instance, let’s say that the proportion of UHP promoters with a TATA box is 32.6% and the proportion of type 0 promoters with a TATA box is 17.2%. Then our statistic of interest is Δ = 32.6 −17.2 = 15.4. We then run the same analysis 1,000,000 times with permutations of the promoters (start with the same collection of promoters and randomize the assignments to UHP, DHP, and type 0 while retaining the same overall numbers). Call the result of each randomizing analysis Δ′. Then our p-value is the proportion of times that Δ′ > Δ.

Since we are performing the above procedure for hundreds of covariates (i.e., several tests for each TFBS), we needed to adjust for multiple testing. This was accomplished by Bonferroni correction and by excluding tests where either |Δ′ −Δ| < 0.5 or |Δ′ −Δ| / Δ < 0.05. Equation (1)

### Functional enrichment analysis

Using isoform-level function assignment from (53), the hypergeometric test was used to determine the probability of observing at least the observed number of UHP-containing isoforms out of the total isoforms annotated to a given GO term that were UHP,DHP or type 0. Only GO terms with at least 5 UHP isoforms annotated to them were considered. Bonferroni correction was applied to the resulting p-values.

## RESULTS

### Association between gene expression and alternative splicing

We focused on alternative splicing events that differentiate between a subset of a gene’s transcripts and the rest of its transcripts. We examined rates of exon inclusion/exclusion in comparison to the overall rate of gene expression in ten tissues with 226 to 653 samples each (Figure 1, Supplemental Table S1).

**Figure 1.**
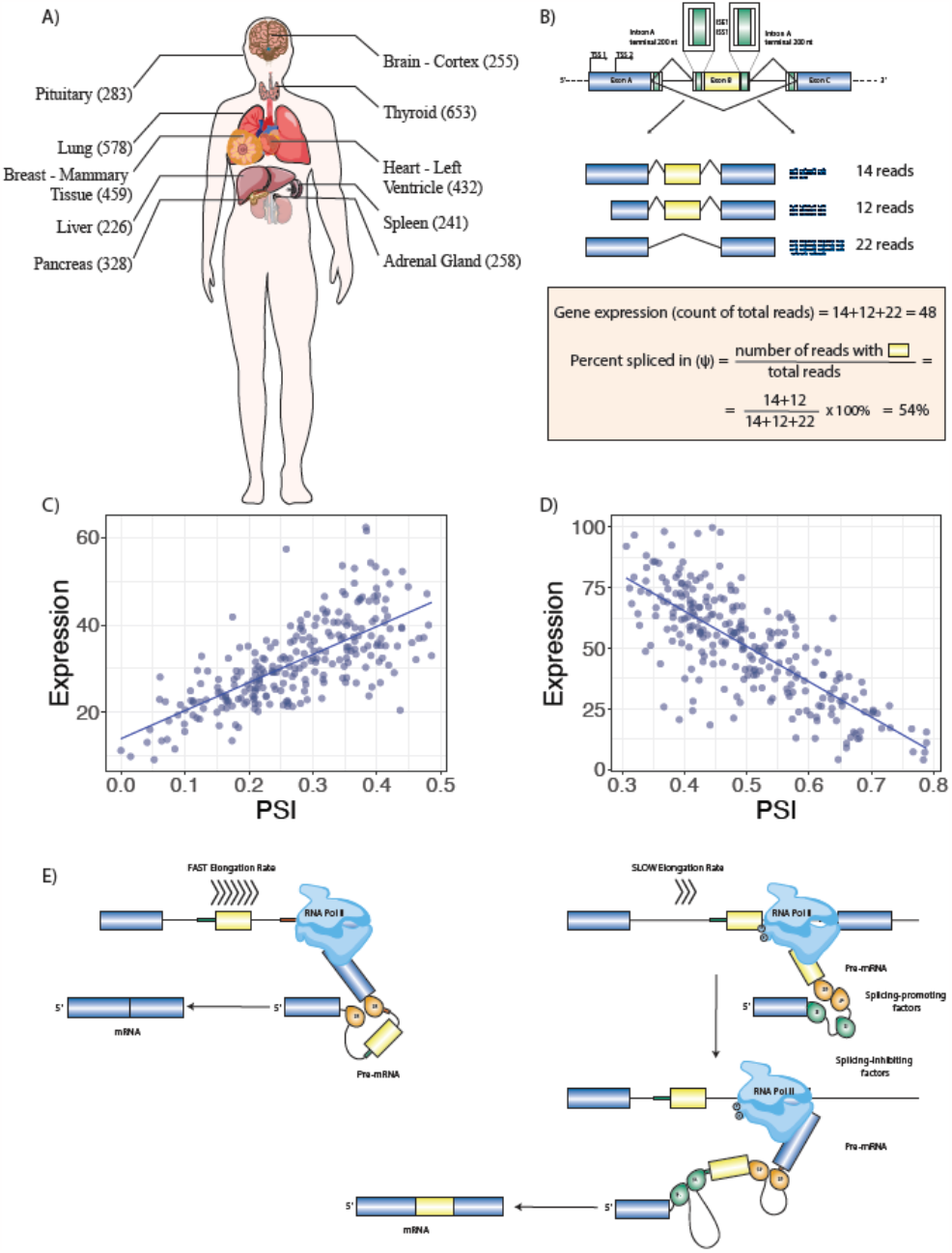
Multitissue RNA-seq Analysis Identifies Association between Gene Expression and Alternative Splicing. **A)** RNA-seq samples from 10 tissues with the largest number of samples were analyzed. **B)** For each of 141,043 alternative splicing events with above-threshold variability in the ten tissues, total gene expression and percent-spliced in (ψ) were calculated and logistic regression was performed to test the association of gene expression and ψ. The cartoon at the top shows the regions of the introns surrounding the cassette exon that were investigated bioinformatically. **C)** 3,667 UHP (for “upregulated-high ψ”) exons with a statistically significant positive association were identified (ψ increases as total gene expression increases). One example is shown, exon 2 of ABI2. **D)** 3,207 DHP (for “downregulated-high ψ”) exons with a statistically significant negative association were identified (ψ decreases as total gene expression increases). In the example, exon 4 of ABLIM2 is shown. **E)** We hypothesized that our observations are related to mechanisms including coupling of RNAP2 extension speed with splicing decisions. In this example, a relatively fast RNAP2 elongation rate exposes a regulatory element (red box) at the 3’ end of intron B, which promotes skipping of exon B (left); in contract, slower RNAP2 elongation fails to expose this element for a period of time sufficient for the splicing machinery to include exon B. This is one of many mechanisms that link transcription and alternative splicing.

### Type 0, upregulated-high ψ (UHP), and downregulated-high ψ (DHP) exons

We filtered 683,196 annotated human exons for those that show a threshold amount of variability in RNA-seq experiments from ten GTEx organ cohorts with between 226 and 653 samples each, identifying 141,043 exons that showed a degree of variable expression equal to or above a threshold of a mean count of at least 20 reads per sample and at least a two-fold ratio of the 95^th^ percentile to the 5^th^ percentile of expression values.

We classified the relationship between overall gene expression and the percent-spliced in (ψ) values of these exons, defining exons where increasing values of ψ (higher exon inclusion) are associated with higher gene expression as UHP exons (“upregulation of gene expression associated with high percent splice in”), exons where increasing values of ψ are associated with lower gene expression as DHP (“downregulation of gene expression associated with high percent splice in”), and exons that show alternative splicing with a significant association between the ψ value and gene expression as type 0 exons. For each of the ten investigated tissues from the GTEx resource, we performed linear regression to predict gene expression based on ψ, and determined the significance of the coefficient for ψ. Raw p-values were corrected for multiple testing by the Benjamini-Hochberg method, and associations are reported as significant at a corrected p-value threshold of 0.05 (Methods).

Using these definitions, we identified 3,667 UHP and 3,207 DHP exons; a total of 6,874 unique exons were identified as UHP or DHP in at least one tissue, corresponding to 4.9% of the 141,043 exons that showed a at least a threshold level of gene expression variability (Methods). 989 exons were identified as UHP or DHP in multiple tissues (Figure 1; additional examples are shown in Supplemental Figure S1). In all, exons were identified as UHP or DHP 8,282 times across the 10 tissues that were tested.

In all 989 cases in which exons were identified as UHP or DHP in multiple tissues, the assignment to UHP or DHP was consistent. We further used the same criteria to find the same UHP/DHP exons in sets of samples that originated from the same donor, for donors with at least 20 tissue samples. A total of 63,961 of the same UHP/DHP exons (1,916 unique exons, ∼89%) were detected in 528 donors. For 63,255 (∼99%) the assignment to UHP or DHP was consistent with the assignment from tissue samples. The small number of inconsistencies is possibly a result of wrong classification due to the relatively small number of samples per donor (a median of 27 samples per donor vs. 342.5 per tissue).

We repeated the same analysis in unrelated breast, left ventricle and liver bulk RNA-seq datasets obtained from the SRA (Methods). In all three datasets, most of the overlapping exons were type 0 in both the GTEx and the SRA dataset, and most of the other exons were type 0 in one of the datasets. For the breast and left ventricle datasets, we observed a highly significant overlap of UHP or DHP classifications between the GTEx and SRA datasets. For liver, there were 52,521 exons that were classified as type 0, 17 exons that were classified as UHP and 47 exons that were classified as DHP. 14 exons were classified as DHP in both datasets, one exon was classified as UHP in both datasets, and all other exons were type 0 in at least one of the datasets (Supplemental Table S2). These results suggest that there is a significant consistency of exon types across different donor cohorts and experimental procedures.

### Minimum prevalence of expression/splicing regulation coupling

In order to estimate how prevalent the coupling between expression and splicing is, we counted the number of exons that were neither detected as UHP nor as DHP, had a 95^th^/5^th^ expression percentile ratio of at least 2, and were assigned a Benjamini-Hochberg-corrected p-value of at least 0.5, in addition to being expressed in at least half the samples in a tissue and at a mean level of 20 transcripts. This definition of type 0 exons intends to identify exons with substantial gene expression variability but with no evidence for being UHP or DHP exons. This resulted in 67,814 cassette exons identified as type 0. Since observing an effect of expression on splicing requires the presence of regulatory factors, such as RNA binding proteins, not observing a correlation does not immediately imply that an exon is type 0 in all tissues. However, since we examined ten different tissues, it is likely that there is roughly an order of magnitude difference between the counts of UHP/DHP exons and type 0 exons (6,874 UHP/DHP vs. 67,814 type 0). In the ten tissue dataset from GTEx, there were a total of 8,314 genes that contained at least one exon classified as UHP, DHP, or type 0. Of these, 1106 genes (13.3%) had at least one UHP or DHP exon.

### Characteristics of type 0, UHP, and DHP exons and the transcript and genes that contain them

UHP/DHP exons differ from type 0 exons in a number of characteristics including exon count, intron length, and distribution of biotypes (Figure 2). Genes containing UHP/DHP exons have on average more exons than genes containing only type 0 exons. The genes containing them had on average slightly fewer transcripts (13 and 12 for UHP and DHP, respectively, and 14 for type 0. Furthermore, type UHP/DHP exons are included in a larger proportion of transcripts than type 0 exons.

**Figure 2.**
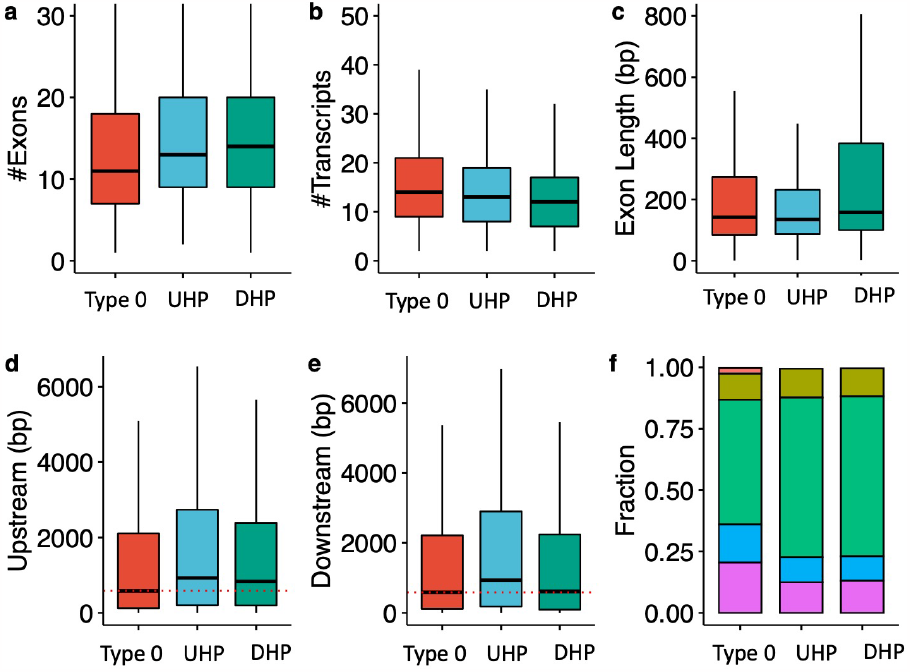
Characteristics of type 0, UHP, and DHP exons. **a)** The number of exons of genes that includes a type 0, UHP, and DHP exon. **b)** The number of transcripts per gene containing a UHP, DHP or type 0 exon. **c)** Exon length in base pairs for type 0, UHP, and DHP exons. **d)** The length of introns upstream of type 0, UHP, and DHP exons. **e)** The length of introns downstream of type 0, UHP, and DHP exons. **f)** Fraction of transcripts of each type associated with different biotypes. Green: protein coding; purple: retained intron; blue: protein coding CDS not defined; khaki: nonsense mediated decay; red: lncRNA. **a-e**: Outliers were removed to limit the y-axis range; **d-e:** The dashed red line shows the median for the up/downstream intron length type 0 exons.

We define the “upstream” intron as the last contiguous non-coding region that is transcribed 5’ to the exon, and the “downstream” intron as the first such region that is transcribed 3’ to the exon. The median upstream intron lengths were 572 bp for types 0, 857 for type UHP, and 732 for DHP; the differences between UHP or DHP and type 0 were statistically significant. In contrast, the median downstream intron lengths were 576 bp for type 0, 834.5 bp for UHP, and 485 bp for DHP. The differences are statistically significant between all types. DHP exons had a median length of 158 bp, which is significantly longer than UHP (median 135 bp) and type 0 (median 142 bp) exons. Finally, transcripts containing UHP/DHP exons have a higher fraction of protein coding transcripts (65 % for UHP/DHP and 50.7 % for type 0 exons), and a smaller fraction of retained introns (12.5 % and 13 % for UHP/DHP, respectively, and 20.5 % for type 0) and long non-coding RNA (0.47 %, 0.36 % and 2.3 % for UHP/DHP, and type 0, respectively) (Figure 2 and Table 1). Additionally, the mean MaxEnt (54) acceptor and donor splice site scores were higher for both UHP and DHP exons than for type 0 exons (Supplemental Figures S3 and S4).

**Table 1.** Characteristics of type 0, UHP, and DHP exons. The values for genes that had both exon types were counted for both types of exons. a) Mann-Whitney test.

| Feature | type 0 | UHP | DHP | 0 vs. UHP | 0 vs. DHP | UHP vs. DHP |
| --- | --- | --- | --- | --- | --- | --- |
| exons per gene<sup>a</sup> | 11 | 13 | 14 | $2.1 \times 10^{-12}$ | $1.7 \times 10^{-14}$ | $6.7 \times 10^{-1}$ |
| transcripts per gene<sup>a</sup> | 14 | 13 | 12 | $7.7 \times 10^{-24}$ | $1.5 \times 10^{-76}$ | $1.06 \times 10^{-10}$ |
| inclusion in proportion of transcripts<sup>a</sup> | 9% | 24.1% | 20% | $2.2 \times 10^{-308}$ | $1.2 \times 10^{-280}$ | $2.4 \times 10^{-4}$ |
| upstream intron length<sup>a</sup> | 584 bp | 930 bp | 832 bp | $1.6 \times 10^{-30}$ | $8.4 \times 10^{-17}$ | 0.03 |
| downstream intron length<sup>a</sup> | 587 bp | 932 bp | 613 bp | $9.8 \times 10^{-23}$ | $1.3 \times 10^{-1}$ | $3.9 \times 10^{-14}$ |
| exon length<sup>a</sup> | 142 bp | 135 bp | 158 bp | 0.03 | $1.9 \times 10^{-32}$ | $7.9 \times 10^{-25}$ |

### High consistency of UHP vs. DHP classification across multiple tissues and datasets

We hypothesized that if the classification of exons as UHP or DHP is related to one or more core regulatory processes, then the classification should be largely conserved across different tissues. Among the detected UHP/DHP exons, there are 194 exons that appear in more than one tissue as DHP always, 160 that appear in more than one tissue always as UHP, and none that appear in more than one tissue as conflicting types. The slopes of the regression lines fitted in different tissues may have different slopes, but the change in slope is correlated across UHP/DHP exons (Supplemental Figure S5). In addition, the slope is a linear function of the mean expression level, with coefficient close to 1, possibly indicating that differences in expression rates affect the impact of UHP/DHP exons on the gene’s transcript profile (Supplemental Figure S2).

### Distribution of RNA polymerase II binding in type 0, UHP, and DHP exons

RNA Pol II accumulates on exons in yeast and human and pauses over the 5’ and 3’ splice sites of human exons (55). Additionally, Pol II density is lower at skipped exons than at alternative retained exons (56, 57). We therefore hypothesized that RNAP2 density might differ between the type 0, UHP, and DHP exons investigated in the current study.

We investigated 105 POLR2A (RNA polymerase II subunit A) ChIP-seq experiments available from the ENCODE project (42). We computed the ratios of the number of POLR2A peaks overlapping UHP/DHP vs. type 0 exons of the same gene (Methods). Both ratios were significantly different than 1 (p=1 × 10^2^ for UHP vs. type 0 and p=1 × 10^−12^ for DHP vs. type 0), and from one another (p=2.3 × 10^−13^), where the median ratio for UHP vs. type 0 positive exon bindings was 0.86, and the median for DHP vs. type 0 was 0.31. These results could suggest that RNAP spends more time on UHP exons compared to DHP exons, and less on DHP exons, possibly due to a different type regulatory interaction that occurs before the decision to include or exclude is made. For example, recruitment time of different types of RBPs may be different. (Figure 3).

**Figure 3.**
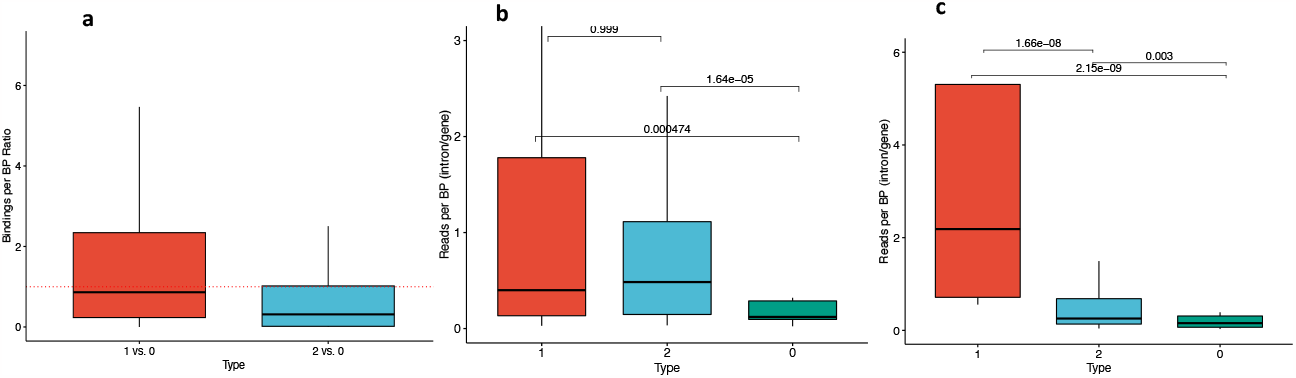
UHP/DHP exons and RNAP2 profiles. **a)** Relative RNAP binding to UHP/DHP exons vs. type 0 on the same gene. The data was obtained from POLR2A peaks over 106 ChIP-Seq experiments. Outliers were removed to limit the y-axis range. **b**,**c)** The ratio between PRO-Seq reads per base pair in downstream introns UHP, DHP, or type 0 exons and the sum of these values for the corresponding gene, for every UHP, DHP, or type 0 exon, from the Wissink et al. dataset **(b)** (45) and the Gupta et al. dataset **(c)** (46). Outliers were removed to limit the y-axis range. The mean values for UHP/DHP exons are significantly larger than type 0, suggesting a longer processing time of that section of the nascent mRNA.

In order to estimate the difference in transcription speed of UHP and DHP exons compared to type 0 exons, we used two PRO-Seq datasets (45, 46) (Methods). These datasets sequenced nascent mRNA in addition to mature mRNA, and therefore allowed reads to be counted in the intronic parts of the nascent mRNA of each gene. The introns downstream of UHP/DHP exons are more likely to be sequenced, suggesting that RNA polymerase spends more time transcribing them (Figure 3c; Mann-Whithney test p =*4*.*7* × *10*^.*4*^ and p =*1*.*6* × *10*^.*5*^ for UHP and DHP vs. type 0 in the Wissink et al dataset, respectively, and p =*2*.*1* × *10*^.*9*^ and *2*.*9* × *10*^.*3*^ for UHP and DHP vs. type 0 in the Gupta et al. dataset, respectively). The longer transcription time may be necessary for the regulatory interactions that promote or suppress the splicing of the exon, and thus may be sensitive to changes in expression rate.

### Enriched motifs

Binding of transcription factors to promoters may influence splicing by altering the rate of RNAP2 elongation or recruiting splicing factors to pre-mRNAs (58). We reasoned that if this were a common factor related to the mechanisms that underlie UHP/DHP exons, then we would expect to see enrichment of predicted transcription factor flexible models (TFFM) sites in the promoter regions of UHP/DHP exons compared to type 0 exons, and would also see enrichments of predicted RBP binding sites in the sequences surrounding the UHP and DHP exons. We therefore calculated the numbers of predicted binding sites and compared the observed counts to those observed in 1,000,000 permutations in which the labels of UHP, DHP, and type 0 exons had been randomly shuffled (Methods).

321 of 610 tested TFFMs showed significant enrichment in genes with UHP or DHP exons but no type 0 exons as compared to genes with at least one type 0 exon but no UHP/DHP exon. However, the maximum difference between the two classes was 3%, suggesting that no individual transcription factor is associated with a majority of the observed effects (Table 2, Supplemental Table S3). We tested enrichment for core promoter elements and CpG islands and found that a significantly higher proportion of DHP genes co-localized with a CpG island and a lower proportion contain a TATA box (Supplemental Table S4). We examined 71 RBP models, 31 of which showed significant differences between UHP or DHP and type 0 exons (Supplemental Table S5).

**Table 2.** Transcription factor flexible models (TFFMs) in promoters of genes harboring UHP and DHP exons. TBFSs were assessed for overrepresentation in genes harboring UHP or DHP exons compared to genes only harboring one or more type 0 exon. 291 models showed a significant difference in permutation testing in which labels of exons (UHP, DHP, type 0) were randomly permuted and the p-value was calculated empirically as the proportion of permutations in which the observed difference between UHP (DHP) and type 0 exons was at least as extreme as the observed difference. The top ten are shown in this table and all results are presented in Supplemental Table 3. Of the significant models, the mean difference was 1.7% (UHP vs. type 0) and 1.1% (DHP vs. type 0). No significant differences were observed between UHP and DHP (not shown). *: significant at a Bonferroni-corrected threshold of 9.11 × 10^−5^.

| motif | model | Type 0 | UHP | DHP | Type 0 vs. UHP | Type 0 vs. DHP | UHP vs. DHP |
| --- | --- | --- | --- | --- | --- | --- | --- |
| PRDM14 | TFFM0987.1 | 41.5% | 44.5% | 44.2% | $p < 1.0 \times 10^{-6} *$ | $p < 1.0 \times 10^{-6} *$ | n.s. |
| SP1 | TFFM0097.2 | 39.2% | 37.9% | 36.8% | n.s. | $p < 1.0 \times 10^{-6} *$ | n.s. |
| KLF4 | TFFM0056.3 | 36.7% | 38.0% | 35.8% | n.s. | n.s. | $6.80 \times 10^{-5} *$ |
| KLF4 | TFFM0056.2 | 35.1% | 35.3% | 33.3% | n.s. | $2.20 \times 10^{-5} *$ | 0.000394 |
| ZNF75D | TFFM0647.1 | 34.4% | 34.2% | 32.3% | n.s. | $p < 1.0 \times 10^{-6} *$ | 0.000751 |
| KLF15 | TFFM0515.1 | 34.1% | 33.8% | 32.0% | n.s. | $p < 1.0 \times 10^{-6} *$ | 0.001070 |
| ZBTB6 | TFFM0624.1 | 32.7% | 30.6% | 30.2% | $p < 1.0 \times 10^{-6} *$ | $p < 1.0 \times 10^{-6} *$ | n.s. |
| FLI1 | TFFM0031.1 | 31.6% | 29.7% | 31.9% | $p < 1.0 \times 10^{-6} *$ | n.s. | $5.90 \times 10^{-5} *$ |
| CTCF | TFFM0014.1 | 29.7% | 31.6% | 30.6% | $p < 1.0 \times 10^{-6} *$ | n.s. | n.s. |
| NEUROD1 | TFFM0143.1 | 30.6% | 28.4% | 29.2% | $p < 1.0 \times 10^{-6} *$ | n.s. | n.s. |

### Biological rationale for coupling expression and splicing

The ubiquity of UHP/DHP led us to further investigate associations of transcript-specific functions. Alternative splicing of many genes can produce isoforms that differ with respect to enzymatic activities and subcellular localizations, as well as protein–protein, protein–DNA, and protein–ligand physical interactions (59). Gene Ontology (GO) overrepresentation analysis is a standard approach to assessing the functional profile of differentially expressed genes (60), but analogous methods for examining the functional profile of differential isoforms have not been available, possibly because of the paucity of experimentally confirmed functional annotations of isoforms (61). We recently developed an expectation-maximization framework for predicting isoform-specific Gene Ontology (GO) annotations (62). We used these annotations to assess overpresentation of GO terms in the set of isoforms that were found to be UHP in our study, with the universe of comparison being the set of all isoforms. A total of 410 GO terms displayed significant overpresentation (See Table 3 for the top ten and Supplemental File 2 for a complete list). possible biological role. The first indication for a functional role comes from the GO terms that are over-represented in UHP isoforms.

**Table 3.** The ten GO terms in which UHP isoforms are most over-represented. Enrichment was tested using the hypergeometric test, where we draw UHP isoforms or other isoforms a number of times that equals the number of isoforms that are annotated to the GO term (Methods). Bonferroni multiple testing correction was applied (adjusted p-value column). Coverage refers to the proportion of isoforms annotated to the term that contain a UHP exon.

| GO Term | Coverage | p.value | adj.p |
| --- | --- | --- | --- |
| JUN kinase activity (GO:0004705) | 0.62 | $2.04 \times 10^{-54}$ | $2.89 \times 10^{-51}$ |
| MAP kinase activity (GO:0004707) | 0.49 | $2.22 \times 10^{-54}$ | $3.13 \times 10^{-51}$ |
| MAP kinase kinase activity (GO:0004708) | 0.59 | $7.68 \times 10^{-50}$ | $1.08 \times 10^{-46}$ |
| response to light stimulus (GO:0009416) | 0.60 | $7.90 \times 10^{-44}$ | $1.11 \times 10^{-40}$ |
| actin binding (GO:0003779) | 0.19 | $2.78 \times 10^{-35}$ | $3.91 \times 10^{-32}$ |
| Fc-epsilon receptor signaling pathway (GO:0038095) | 0.51 | $2.66 \times 10^{-33}$ | $3.74 \times 10^{-30}$ |
| GPI-anchor transamidase complex (GO:0042765) | 0.65 | $6.37 \times 10^{-31}$ | $8.95 \times 10^{-28}$ |
| cytoskeleton organization (GO:0007010) | 0.32 | $7.16 \times 10^{-31}$ | $1.01 \times 10^{-27}$ |
| cytoskeletal protein binding (GO:0008092) | 0.31 | $1.30 \times 10^{-30}$ | $1.82 \times 10^{-27}$ |
| cellular senescence (GO:0090398) | 0.44 | $1.02 \times 10^{-29}$ | $1.43 \times 10^{-26}$ |

Interestingly, the top three overrepresented terms, JUN kinase activity(supplementary figure S6), MAP kinase activity, and MAP kinase kinase activity annotate genes and pathways that coordinately regulate gene expression, mitosis, metabolism, motility, survival, apoptosis, differentiation and protection against DNA damage and deleterious mutations (63–65). Our results suggest that cells can upregulate these pathways both by increasing overall expression of genes in the pathway and also by alternative splicing to favor transcription of isoforms that specifically possess pathway activity.

We investigated the association between the degree of differential expression and the degree of alternative splicing by regression ψ against gene expression level for all exons identified as UHP/DHP. We found that the slope of the expression-ψ line is approximately the mean expression level of the gene (Supplementary Figure S2). Therefore, in dividing cells, which need to double their protein content, increasing expression of genes annotated to JUN kinase activity, MAP kinase activity, and MAP kinase kinase activity will additionally strongly favor transcription of UHPs that also are specifically annotated to these GO terms. Thus, increased gene expression will both increase the overall amount of genes and shift their transcript distributions to transcripts with Jun kinase activity. In order to test this hypothesis, we computed the Pearson correlation between UHP exons *ψ* and Cyclin D1 gene expression, as the latter is a marker for the level of mitosis (66). This correlation is mostly positive, while the same correlation between DHP/Type 0 exon *ψ* and Cyclin D1 expression is mostly negative (supplementary figure S7). We further calculated correlation between UHP exons *ψ* and Cyclin D1 expression in The Cancer Genome Atlas (TCGA) transcript expression dataset, for primary tumor and normal tissue, and found a significant reduction in correlation in tumor samples (Supplementary Figure S8).

## DISCUSSION

We developed an approach to characterize associations between overall gene expression, defined as the sum of read counts for all transcripts assigned to a gene, and the regulation of alternative splicing, defined as the inclusion or exclusion of an exon belonging to some, but not all, transcripts of the gene. We identified exons whose exclusion or inclusion was correlated with total gene expression. UHP (upregulated-high ψ) exons show a significant association of higher overall gene expression with higher degrees of exon inclusion, and DHP (downregulated-high ψ) exons show a significant association of lower overall gene expression with higher degrees of exon inclusion. It is likely that the total number of such exons identified by our study, 3,667 UHP exons and 3,207 DHP exons, corresponding to a total of 6,874 exons in 1106 genes, represents a lower bound, because the experiments investigated in our study do not comprise a sufficient range of conditions to assay a sufficiently variable range of expression and splicing to detect all UHP and DHP exons.

A previous work assayed RNAP2 mutants that change average elongation rates genome-wide and showed two classes of cassette exons that displayed higher degrees of inclusion with slower RNAP2 mutants (type I) and lower degrees of inclusion with faster RNAP2 mutants (type II). The type I exons tended to have weaker splice sites, to be surrounded by shorter introns compared to type II exons, and to harbor distinct sequence motifs (30). The exons identified by this work were mapped to the hg19 genome, and splicing was quantified using the MATS tool, which does not reconstruct full transcripts, limiting comparability with our results.

Speculatively, however, the association of type I/II as well as of UHP/DHP exons with intron length, splice site strength, and sequence motifs could indicate partially shared mechanism, with differences being due to the fact that the previous study was investigating global changes of RNAP2 extension speed.

Our study identified significant differences in the strength of splice sites, intron and exon length, and different proportions of predicted TFBS in promoter regions of gene harboring UHP/DHP exons compared to genes with type 0 exons. Additionally, we identified a significantly higher relative RNAP binding to UHP/DHP exons vs. type 0 on the same gene in data from 106 POLR2A ChIP-Seq experiments, and a higher count of nascent RNA reads per base pair in introns downstream of UHP and DHP exons as compared to type 0 exons, suggesting a role of RNAP2 in mediating the observed effects. The consistency of UHP/DHP classification across tissues of the direction of correlation between expression and exon proportion suggests an intrinsic mechanism that is not the sole result of epigenetic modifications. Our interpretation is that local modulation of transcription speed (55) could play a role in modulation of alternative splicing. In our study, we identified 141,043 exons with a mean count of at least 20 reads per sample and at least a two-fold ratio of the 95^th^ percentile to the 5^th^ percentile of expression values. Of these, 4.8% were classified as either UHP or DHP. We expect that the figure of 4.8% of exons displaying a significant relation between splicing and expression is a lower bound, and that comprehensive profiling of large-scale datasets representing a wider range of tissues, developmental stages, and disease states may reveal additional instances of coupled splicing and expression regulation. 1106 genes, corresponding to 13.3% of genes with non-trivial expression in the ten investigated GTEx tissues, contained at least one UHP or DHP exon. This proposed mechanism is decentralized and stems from intrinsic properties of transcription.

Finally, UHP/DHP exons are enriched for hundreds of distinct GO terms, suggesting that the coupling between expression and alternative splicing may provide an important gene regulatory mechanism that might be used in a variety of biological contexts.

## DATA AVAILABILITY

The data used in this study are available at the NCBI Sequence Read Archive (SRA).(40) The individual datasets can be downloaded by using the Snakemake(67) script that is provided under an MIT License at https://github.com/TheJacksonLaboratory/gene_exp_psi. The The Snakemake file additionally runs a collection of scripts that were used to generate the main results presented in the manuscript. Source code for the C++ application used to analyze motifs associated with UHP and DHP exons is also provided at the GitHub repository. Any additional information required to reanalyze the data reported in this paper is available from the corresponding author upon request.

## SUPPLEMENTARY DATA

Supplementary Data are available at NAR online.

## ACKNOWLEDGEMENTS

The authors would like to thank Dr. Gloria Fuentes, The Visual Thinker, for designing Figure 1.

