## Supplementary material for "Alternative splicing is coupled to gene expression in a subset of variably expressed genes": Online supplement

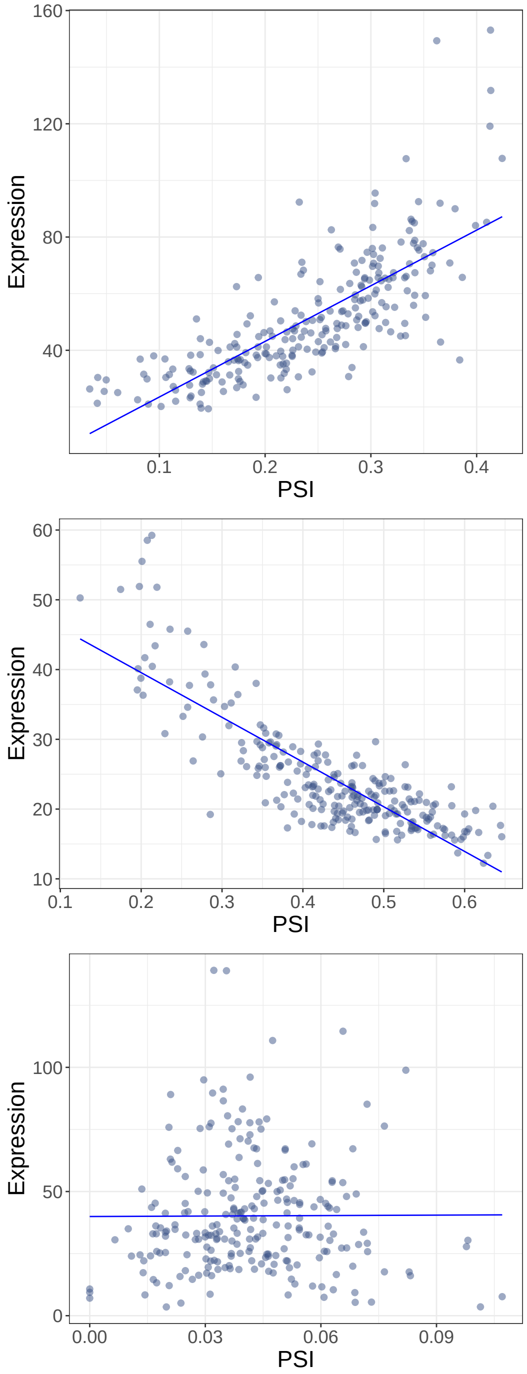


**Figure S1: The expression of genes containing UHP (upregulated-high $\psi$), DHP (downregulated-high $\psi$), and type 0 exons as a function of the proportion of transcripts including these exons**. The expression value and proportions were computed in Spleen. The x-axis shows the proportion of transcript counts for transcripts that include the exon (percent-spliced in, $\psi$), and the y-axis value is the gene expression or total number of transcript counts for the gene. The genes for the UHP, DHP and type 0 exons displayed in this figure, *CASP8*, *CSTF3* and *ANPEP*, are shown from top to bottom.

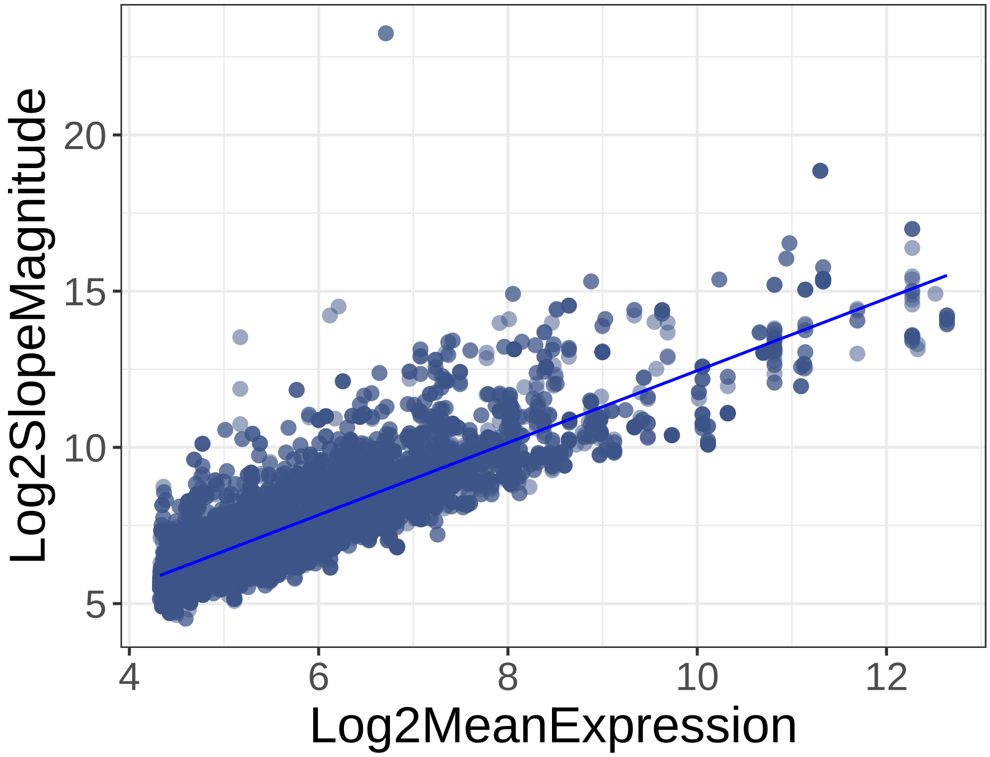


**Figure S2:** $Log_2$ **Expression-**$\psi$ **regression slopes vs** $log_2$ **mean expression.** This Figure the mean expression of genes with at least one UHP or DHP exon (X-axis) with the absolute value of the slope of the the corresponding expression-percent-spliced-in ($\psi$) regression curve (Y-axis).

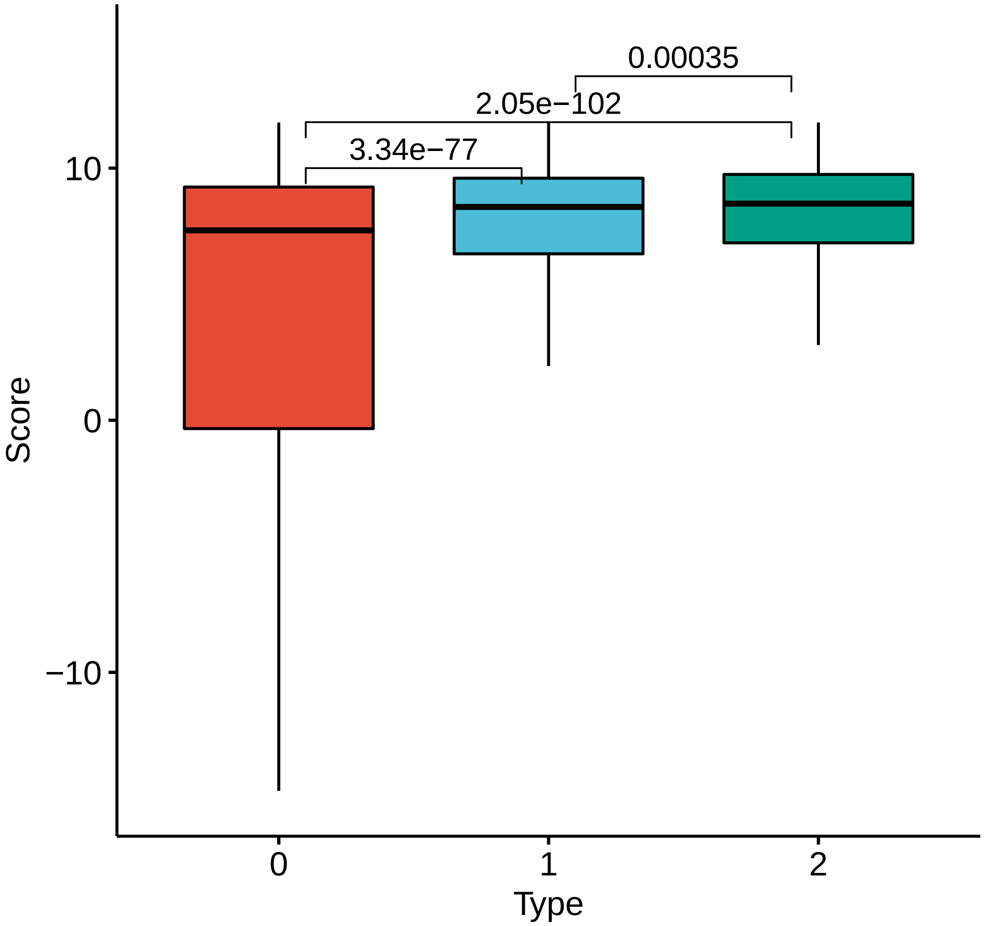


**Figure S3: 5’ (donor) splice score distributions**. Boxplots illustrate the distribution of the 5’ donor splice score calculated using MaxEntScan (39). 0: type 0 exon; 1: UHP exon; 2: DHP exon.

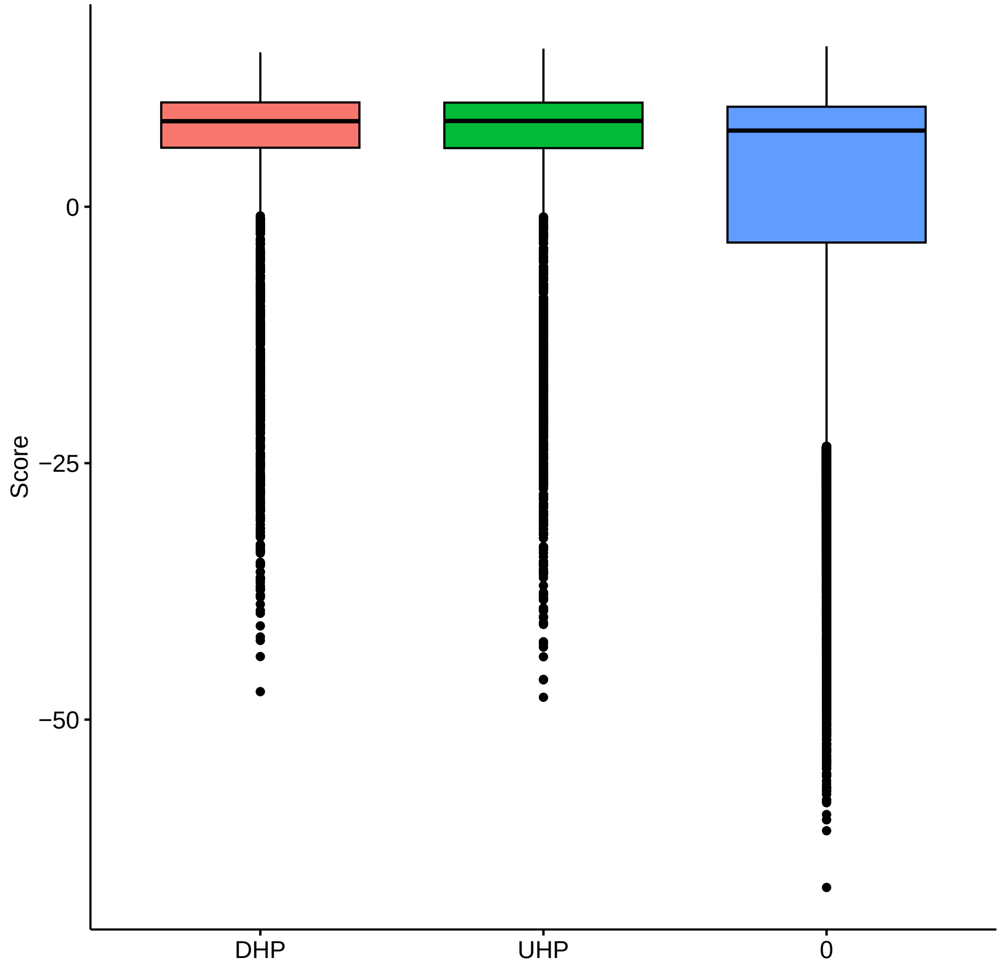


**Figure S4: 3' (acceptor) splice score distributions**. Boxplots illustrate the distribution of the 3' donor splice score calculated using MaxEntScan (39). The Mann-Whitney p-value of DHP vs type 0 was $p = 2.09^{-43}$, of UHP vs type 0 is $p = 1.19^{-48}$, and UHP vs DHP was $p = 0.95$. 0: type 0 exon; 1: UHP exon; 2: DHP exon.

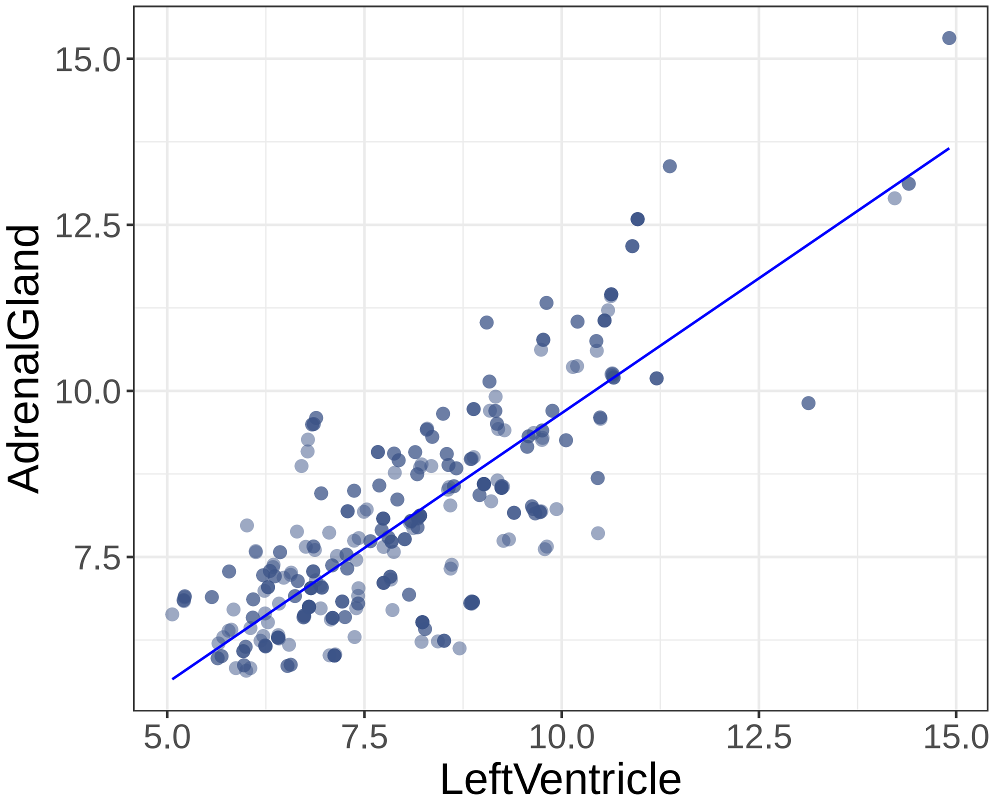


**Figure S5: Expression-$\psi$ log2-regression slopes in Heart left ventricle vs adrenal gland.** Each point represents the absolute value of the slope of the expression-percent-spliced-in ($\psi$) regression curves for one UHP or DHP exon.

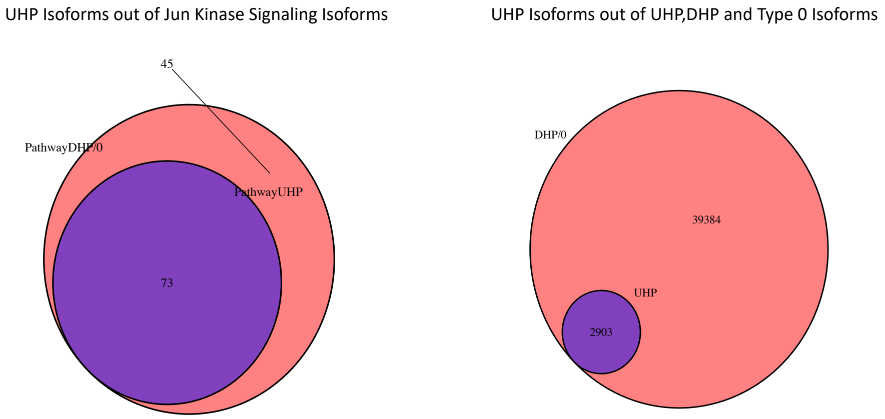


**Figure S6: Over-representation UHP-containing isoforms**. Proportion of UHP-containing isoforms out of all isoforms belonging to the Jun kinase signaling GO term (left) and the same proportion out of all isoforms containing UHP,DHP or a type 0 exon. The isoforoms are over-represented in the Jun kinase signaling GO term.(Benjamini-Hochberg corrected hyper geometric $p = 1.57^{-51}$)

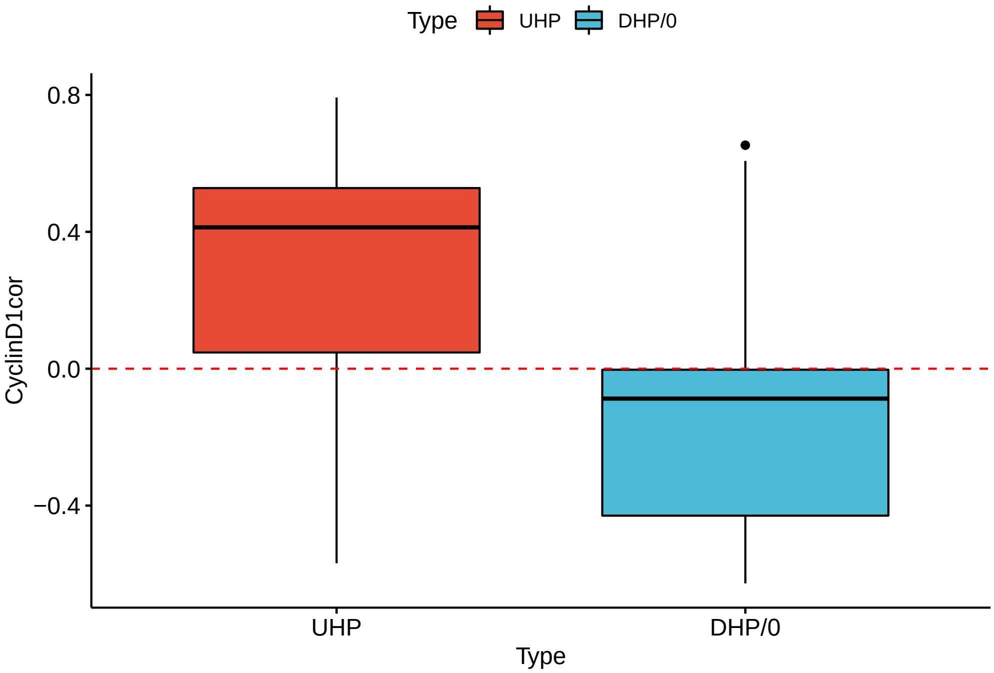


**Figure S7: Pearson correlation between PSI of UHD/DHP/Type 0 exons and Cyclin D1 gene expression in the GTEx dataset**. Pearson correlation between the expression levels of Cyclin D1 and the PSI of every exon was computed across all GTEx tissues that were examined in this study. Cyclin D1 expression is a proxy for the level of mitosis. As thew figure shows, UHP exons are mostly positively correlated with Cyclin D1 expression, and other exons are mostly negatively correlated.

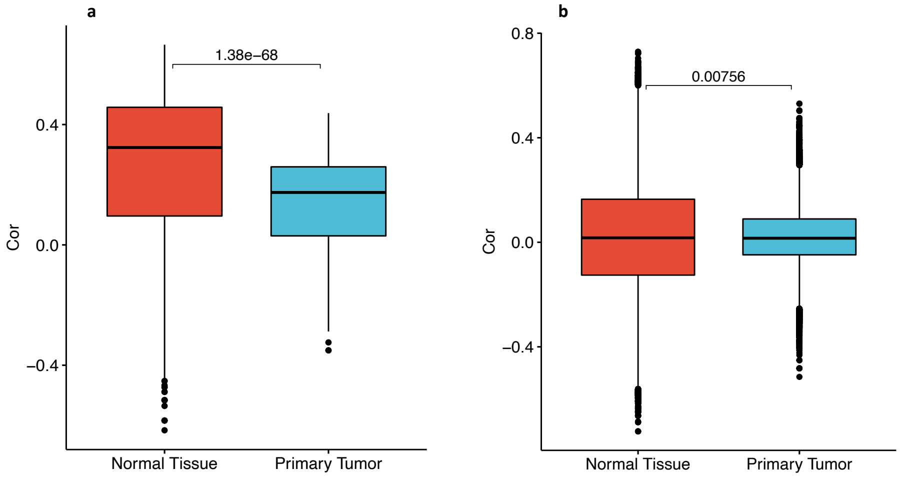


**Figure S8: Pearson correlation between PSI of UHD/Type 0 exons and Cyclin D1 gene expression in The Cancer Genome Atlas transcript expression dataset**. Pearson correlation between the expression levels of Cyclin D1 and the PSI of every exon was computed in TCGA samples classified as 'Thyroid Carcinoma', separately in the 'Solid Tissue Normal' and 'Primary Tumor' sub-categories, for UHP exons (a) and Type 0 exons (b). UHP exons are mostly positively correlated with Cyclin D1 expression, and Type 0 exons have a median correlation of approximately 0. In tumor the correlation is significantly reduced compared to healthy tissue (Mann-Whitney p-values displayed in the figure), with the gap being significantly larger for UHP exons (for UHP medians of 0.32 and 0.17 for normal and tumor tissues, respectively, and for Type 0 exons medians of 0.017 and 0.016, for normal and tumor tissues, respectively).

| Tissue | #Samples |
|---|---|
| Adrenal Gland | 258 |
| Brain - Cortex | 255 |
| Breast - Mammary Tissue | 459 |
| Heart - Left Ventricle | 432 |
| Liver | 226 |
| Lung | 578 |
| Pancreas | 328 |
| Pituitary | 283 |
| Spleen | 241 |
| Thyroid | 653 |

**Table S1:** Organs included in the analysis. The column "#Samples" shows the number of samples used from the GTEx RNA-seq resource (37) for the current analysis.

| Dataset | Breast tissue | Left Ventricle | Liver |
|---|---|---|---|
| SRA | SRP301453 | SRP237337 | SRP326468 |
| DHP/DHP | 115 | 128 | 14 |
| UHP/UHP | 34 | 179 | 1 |
| DHP/UHP | 7 | 41 | 0 |
| UHP/DHP | 5 | 53 | 0 |
| FET | $1.15 \times 10^{-22}$ | $1.54 \times 10^{-26}$ | $6.67 \times 10^{-2}$ |

**Table S2: UHP/DHP/type 0 exon analysis on three external datasets**. We repeated our analysis of UHP/DHP/type 0 exons on three external datasets from NCBI's sequence read archive (SRA) (48). In all three datasets, most of the overlapping exons were type 0 in both the GTEx and the SRA dataset, and most of the other exons were type 0 in one of the datasets (not shown in the table). Rows such as "DHP/UHP" show the counts of exons that were classified as indicated in the SRA/GTEx datasets. FET: Fischer's Exact Text p-value. These results suggest that there is a significant consistency of exon types across different donor cohorts and experimental procedures.

**Table S3:** Frequency of occurrence of Transcription Factor Binding Sites (TFBS). See main text for definitions of `type 0`, `UHP`, and `DHP`. The Bonferroni corrected threshold of $\alpha = 0.05$ is $9.11 \times 10^{-5}$. *) significant at this threshold.

| TFBS | type 0 | UHP | DHP | 0 vs. UHP | 0 vs DHP | UHP vs. DHP |
|---|---|---|---|---|---|---|
| ARNT2 (TFFM0853.1) | 9.0% | 9.7% | 10.5% | 0.006970 | $p < 10^{-6}$ * | 0.008200 |
| ARNTL (TFFM0162.1) | 1.7% | 2.4% | 2.4% | $p < 10^{-6}$ * | $p < 10^{-6}$ * | n.s |
| ASCL1 (TFFM0131.1) | 24.6% | 21.6% | 22.0% | $p < 10^{-6}$ * | $p < 10^{-6}$ * | n.s |
| ASCL1 (TFFM0131.2) | 27.6% | 25.3% | 25.5% | $p < 10^{-6}$ * | $p < 10^{-6}$ * | n.s |
| ASCL1 (TFFM0890.1) | 28.2% | 25.2% | 25.6% | $p < 10^{-6}$ * | $p < 10^{-6}$ * | n.s |
| ASCL2 (TFFM0440.1) | 14.7% | 12.5% | 13.3% | $p < 10^{-6}$ * | $p < 10^{-6}$ * | 0.041738 |
| ATF2 (TFFM0653.1) | 6.1% | 7.4% | 7.2% | $p < 10^{-6}$ * | $p < 10^{-6}$ * | n.s |
| ATF3 (TFFM0003.2) | 5.4% | 6.6% | 6.3% | $p < 10^{-6}$ * | $3.90 \times 10^{-5}$ * | n.s |
| ATF4 (TFFM0163.2) | 2.2% | 2.4% | 3.1% | n.s | $p < 10^{-6}$ * | $7.90 \times 10^{-5}$ * |
| BCL6 (TFFM0006.1) | 18.0% | 16.1% | 16.4% | $p < 10^{-6}$ * | $p < 10^{-6}$ * | n.s |
| BHLHE22 (TFFM0165.1) | 26.2% | 23.3% | 23.2% | $p < 10^{-6}$ * | $p < 10^{-6}$ * | n.s |
| BHLHE22 (TFFM0794.1) | 26.0% | 23.1% | 23.2% | $p < 10^{-6}$ * | $p < 10^{-6}$ * | n.s |
| BHLHE22 (TFFM0892.1) | 25.6% | 22.6% | 22.7% | $p < 10^{-6}$ * | $p < 10^{-6}$ * | n.s |
| CREB1 (TFFM0012.2) | 4.4% | 5.5% | 5.3% | $p < 10^{-6}$ * | $p < 10^{-6}$ * | n.s |
| CTCFL (TFFM0133.2) | 21.5% | 23.8% | 23.4% | $p < 10^{-6}$ * | $p < 10^{-6}$ * | n.s |
| ELF3 (TFFM0170.2) | 10.3% | 8.5% | 8.5% | $p < 10^{-6}$ * | $p < 10^{-6}$ * | n.s |
| ELF4 (TFFM0472.1) | 20.6% | 18.6% | 20.0% | $p < 10^{-6}$ * | n.s | 0.001419 |
| ETS2 (TFFM0858.1) | 8.5% | 6.9% | 6.5% | $p < 10^{-6}$ * | $p < 10^{-6}$ * | n.s |
| ETV4 (TFFM0173.1) | 14.2% | 12.3% | 12.5% | $p < 10^{-6}$ * | $p < 10^{-6}$ * | n.s |
| ETV5 (TFFM0480.1) | 7.8% | 6.9% | 6.1% | 0.000110 | $p < 10^{-6}$ * | 0.003721 |
| ETV5 (TFFM0785.1) | 4.9% | 4.2% | 3.7% | $7.00 \times 10^{-5}$ * | $p < 10^{-6}$ * | n.s |
| ETV6 (TFFM0174.1) | 20.0% | 18.2% | 19.4% | $p < 10^{-6}$ * | n.s | 0.006593 |
| FOXA2 (TFFM0036.1) | 13.8% | 12.7% | 11.9% | 0.000145 | $p < 10^{-6}$ * | 0.032277 |
| GABPA (TFFM0039.2) | 25.7% | 26.7% | 28.2% | n.s | $p < 10^{-6}$ * | 0.002089 |
| GLIS1 (TFFM0492.1) | 12.5% | 14.9% | 14.8% | $p < 10^{-6}$ * | $p < 10^{-6}$ * | n.s |
| GMEB2 (TFFM0494.1) | 6.1% | 7.6% | 7.7% | $p < 10^{-6}$ * | $p < 10^{-6}$ * | n.s |
| HOXA13 (TFFM0504.1) | 5.0% | 4.4% | 3.8% | 0.001868 | $p < 10^{-6}$ * | 0.008876 |
| HOXB13 (TFFM0180.1) | 3.2% | 2.8% | 2.3% | n.s | $p < 10^{-6}$ * | 0.006023 |
| HOXC9 (TFFM0047.1) | 6.6% | 5.6% | 5.5% | $p < 10^{-6}$ * | $p < 10^{-6}$ * | n.s |
| IKZF1 (TFFM0509.1) | 17.9% | 15.2% | 16.2% | $p < 10^{-6}$ * | $p < 10^{-6}$ * | 0.016385 |
| IRF8 (TFFM0511.1) | 14.4% | 12.4% | 13.3% | $p < 10^{-6}$ * | 0.000216 | 0.019470 |
| KLF9 (TFFM0827.1) | 15.7% | 17.6% | 16.1% | $p < 10^{-6}$ * | n.s | 0.000287 |
| MITF (TFFM0141.2) | 7.1% | 8.1% | 8.6% | $5.30 \times 10^{-5}$ * | $p < 10^{-6}$ * | 0.044470 |
| MNT (TFFM0191.1) | 7.4% | 8.6% | 8.7% | $p < 10^{-6}$ * | $p < 10^{-6}$ * | n.s |
| MZF1 (TFFM0531.1) | 8.7% | 11.2% | 10.8% | $p < 10^{-6}$ * | $p < 10^{-6}$ * | n.s |
| NEUROD1 (TFFM0143.1) | 30.6% | 28.4% | 29.2% | $p < 10^{-6}$ * | n.s | n.s |
| NFIC (TFFM0072.1) | 6.1% | 7.7% | 7.6% | $p < 10^{-6}$ * | $p < 10^{-6}$ * | n.s |
| NFIC (TFFM0072.2) | 1.6% | 1.0% | 1.2% | $p < 10^{-6}$ * | n.s | n.s |
| NFIX (TFFM0761.1) | 18.4% | 16.1% | 17.1% | $p < 10^{-6}$ * | 0.000178 | 0.013094 |
| NFYB (TFFM0075.1) | 5.9% | 4.9% | 5.7% | $p < 10^{-6}$ * | n.s | 0.002848 |
| NKX2-5 (TFFM0076.1) | 1.2% | 1.7% | 1.8% | $p < 10^{-6}$ * | $p < 10^{-6}$ * | n.s |

Continued on next page

**Table S3 – continued from previous page**

| TFBS | type 0 | UHP | DHP | 0 vs. UHP | 0 vs DHP | UHP vs. DHP |
|---|---|---|---|---|---|---|
| NR2F1 (TFFM0869.1) | 7.7% | 6.5% | 6.4% | $p < 10^{-6}$ * | $p < 10^{-6}$ * | n.s |
| NR5A1 (TFFM0543.1) | 5.1% | 4.1% | 4.0% | $p < 10^{-6}$ * | $p < 10^{-6}$ * | n.s |
| NRF1 (TFFM0082.1) | 12.2% | 14.3% | 15.2% | $p < 10^{-6}$ * | $p < 10^{-6}$ * | 0.012840 |
| NRF1 (TFFM0732.1) | 12.4% | 15.0% | 15.0% | $p < 10^{-6}$ * | $p < 10^{-6}$ * | n.s |
| PLAG1 (TFFM0561.1) | 15.2% | 17.7% | 16.9% | $p < 10^{-6}$ * | $p < 10^{-6}$ * | n.s |
| PRDM14 (TFFM0987.1) | 41.5% | 44.5% | 44.2% | $p < 10^{-6}$ * | $p < 10^{-6}$ * | n.s |
| RFX3 (TFFM0791.1) | 3.8% | 5.0% | 4.3% | $p < 10^{-6}$ * | n.s | 0.000808 |
| SNAI1 (TFFM0877.1) | 5.7% | 4.1% | 4.1% | $p < 10^{-6}$ * | $p < 10^{-6}$ * | n.s |
| SNAI2 (TFFM0203.1) | 13.5% | 11.5% | 11.4% | $p < 10^{-6}$ * | $p < 10^{-6}$ * | n.s |
| SOX3 (TFFM0734.1) | 8.3% | 7.6% | 7.1% | 0.002438 | $p < 10^{-6}$ * | n.s |
| SP1 (TFFM0097.2) | 39.2% | 37.9% | 36.8% | n.s | $p < 10^{-6}$ * | n.s |
| SP5 (TFFM0985.1) | 4.6% | 3.4% | 4.1% | $p < 10^{-6}$ * | 0.002486 | 0.004002 |
| SPI1 (TFFM0099.1) | 20.0% | 18.1% | 20.1% | $p < 10^{-6}$ * | n.s | $4.70 \times 10^{-5}$ * |
| SPIB (TFFM0204.1) | 15.0% | 13.7% | 16.0% | $5.60 \times 10^{-5}$ * | 0.001336 | $p < 10^{-6}$ * |
| SPIB (TFFM0204.2) | 11.8% | 9.8% | 10.9% | $p < 10^{-6}$ * | 0.002175 | 0.001131 |
| STAT2 (TFFM0207.1) | 5.2% | 5.8% | 6.2% | 0.002084 | $p < 10^{-6}$ * | n.s |
| STAT2 (TFFM0593.1) | 8.0% | 7.1% | 6.8% | 0.000223 | $p < 10^{-6}$ * | n.s |
| TCF12 (TFFM0736.1) | 4.1% | 3.1% | 3.3% | $p < 10^{-6}$ * | $p < 10^{-6}$ * | n.s |
| TCF3 (TFFM0108.1) | 25.1% | 22.3% | 22.3% | $p < 10^{-6}$ * | $p < 10^{-6}$ * | n.s |
| TCF4 (TFFM0601.1) | 16.1% | 14.3% | 14.1% | $p < 10^{-6}$ * | $p < 10^{-6}$ * | n.s |
| TEAD2 (TFFM0153.1) | 8.9% | 10.2% | 9.5% | $p < 10^{-6}$ * | 0.015357 | 0.019089 |
| TFAP2A (TFFM0112.1) | 13.3% | 11.3% | 12.9% | $p < 10^{-6}$ * | n.s | $8.70 \times 10^{-5}$ * |
| TFAP2B (TFFM0114.1) | 11.3% | 9.5% | 11.7% | $p < 10^{-6}$ * | n.s | $p < 10^{-6}$ * |
| TFE3 (TFFM0798.1) | 7.8% | 9.0% | 8.9% | $p < 10^{-6}$ * | $p < 10^{-6}$ * | n.s |
| TFEB (TFFM0768.1) | 13.5% | 15.2% | 15.2% | $p < 10^{-6}$ * | $p < 10^{-6}$ * | n.s |
| THAP11 (TFFM0608.1) | 2.3% | 2.8% | 3.1% | 0.000171 | $p < 10^{-6}$ * | n.s |
| VSX2 (TFFM0775.1) | 5.5% | 4.4% | 4.8% | $p < 10^{-6}$ * | 0.000384 | n.s |
| YY1 (TFFM0124.1) | 11.2% | 13.2% | 13.3% | $p < 10^{-6}$ * | $p < 10^{-6}$ * | n.s |
| YY1 (TFFM0714.1) | 6.2% | 7.6% | 7.9% | $p < 10^{-6}$ * | $p < 10^{-6}$ * | n.s |
| YY2 (TFFM0621.2) | 17.5% | 19.7% | 19.2% | $p < 10^{-6}$ * | $p < 10^{-6}$ * | n.s |
| ZBTB6 (TFFM0624.1) | 32.7% | 30.6% | 30.2% | $p < 10^{-6}$ * | $p < 10^{-6}$ * | n.s |
| ZFP42 (TFFM0695.1) | 9.3% | 12.0% | 10.7% | $p < 10^{-6}$ * | $p < 10^{-6}$ * | 0.000105 |
| ZIM3 (TFFM0908.1) | 3.2% | 2.2% | 2.2% | $p < 10^{-6}$ * | $p < 10^{-6}$ * | n.s |
| ZNF135 (TFFM0632.1) | 16.9% | 15.5% | 15.1% | $4.50 \times 10^{-5}$ * | $p < 10^{-6}$ * | n.s |
| ZNF341 (TFFM0700.1) | 14.4% | 12.5% | 12.5% | $p < 10^{-6}$ * | $p < 10^{-6}$ * | n.s |
| ZNF417 (TFFM0920.1) | 13.7% | 15.7% | 14.7% | $p < 10^{-6}$ * | 0.002549 | 0.007461 |
| ZNF708 (TFFM0923.1) | 5.2% | 6.5% | 5.9% | $p < 10^{-6}$ * | 0.000448 | 0.013277 |
| ZNF816 (TFFM0914.1) | 10.6% | 10.2% | 9.3% | n.s | $p < 10^{-6}$ * | 0.009962 |
| ZNF93 (TFFM0916.1) | 19.7% | 20.5% | 21.6% | n.s | $p < 10^{-6}$ * | 0.008743 |
| ATF1 (TFFM0002.1) | 13.1% | 14.6% | 14.2% | $p < 10^{-6}$ * | 0.000141 | n.s |
| CTCF (TFFM0014.1) | 29.7% | 31.6% | 30.6% | $p < 10^{-6}$ * | n.s | n.s |
| E2F1 (TFFM0016.1) | 13.7% | 14.7% | 15.2% | 0.000645 | $p < 10^{-6}$ * | n.s |
| ELF1 (TFFM0022.2) | 13.5% | 12.1% | 11.9% | $p < 10^{-6}$ * | $p < 10^{-6}$ * | n.s |
| ETV4 (TFFM0173.2) | 14.7% | 13.1% | 13.5% | $p < 10^{-6}$ * | 0.000197 | n.s |

Continued on next page

| TFBS | type 0 | UHP | DHP | 0 vs. UHP | 0 vs DHP | UHP vs. DHP |
|---|---|---|---|---|---|---|
| ETV4 (TFFM0784.1) | 11.7% | 10.5% | 10.4% | $3.80 \times 10^{-5}$ * | $p < 10^{-6}$ * | n.s |
| HSF1 (TFFM0048.1) | 4.4% | 5.3% | 5.0% | $p < 10^{-6}$ * | 0.000447 | n.s |
| JUND (TFFM0054.1) | 4.5% | 5.5% | 5.3% | $p < 10^{-6}$ * | $1.30 \times 10^{-5}$ * | n.s |
| KLF15 (TFFM0515.1) | 34.1% | 33.8% | 32.0% | n.s | $p < 10^{-6}$ * | 0.001070 |
| RFX1 (TFFM0733.1) | 6.6% | 7.8% | 7.5% | $p < 10^{-6}$ * | $9.80 \times 10^{-5}$ | n.s |
| SP4 (TFFM0591.1) | 26.5% | 25.8% | 24.6% | n.s | $p < 10^{-6}$ * | n.s |
| TFCP2L1 (TFFM0719.1) | 17.7% | 16.2% | 17.4% | $p < 10^{-6}$ * | n.s | 0.004856 |
| ZIC3 (TFFM0771.1) | 17.7% | 19.4% | 18.3% | $p < 10^{-6}$ * | n.s | 0.008155 |
| ZNF75D (TFFM0647.1) | 34.4% | 34.2% | 32.3% | n.s | $p < 10^{-6}$ * | 0.000751 |
| NFYA (TFFM0074.2) | 8.4% | 8.4% | 9.6% | n.s | $p < 10^{-6}$ * | 0.000330 |
| PPARG (TFFM0086.1) | 8.4% | 9.4% | 9.6% | $3.70 \times 10^{-5}$ * | $p < 10^{-6}$ * | n.s |
| SOX9 (TFFM0710.1) | 2.8% | 2.1% | 2.4% | $p < 10^{-6}$ * | n.s | n.s |
| SPI1 (TFFM0713.1) | 14.7% | 13.2% | 14.2% | $p < 10^{-6}$ * | n.s | 0.009807 |
| TBR1 (TFFM0792.1) | 1.7% | 1.4% | 1.2% | n.s | $p < 10^{-6}$ * | n.s |
| USF2 (TFFM0123.2) | 10.8% | 12.2% | 11.7% | $p < 10^{-6}$ * | 0.000614 | n.s |
| ESRRG (TFFM0753.1) | 2.3% | 1.7% | 1.9% | $p < 10^{-6}$ * | n.s | n.s |
| HOXC9 (TFFM0047.2) | 5.8% | 5.0% | 4.8% | 0.000209 | $p < 10^{-6}$ * | n.s |
| NR5A1 (TFFM0871.1) | 3.0% | 2.3% | 2.5% | $p < 10^{-6}$ * | n.s | n.s |
| FOXH1 (TFFM0037.1) | 1.9% | 1.4% | 1.5% | $p < 10^{-6}$ * | n.s | n.s |
| TFAP2C (TFFM0211.1) | 10.1% | 9.7% | 8.9% | n.s | $p < 10^{-6}$ * | 0.008757 |
| ZNF684 (TFFM0646.1) | 1.5% | 1.9% | 2.0% | n.s | $p < 10^{-6}$ * | n.s |
| IRF4 (TFFM0182.1) | 6.1% | 5.2% | 5.5% | $p < 10^{-6}$ * | 0.000962 | n.s |
| PRDM4 (TFFM0688.1) | 7.8% | 9.0% | 8.6% | $p < 10^{-6}$ * | 0.002101 | n.s |
| TCF12 (TFFM0899.1) | 11.8% | 11.4% | 10.5% | n.s | $p < 10^{-6}$ * | 0.008528 |
| FLI1 (TFFM0031.1) | 31.6% | 29.7% | 31.9% | $p < 10^{-6}$ * | n.s | $5.90 \times 10^{-5}$ * |
| NR2C2 (TFFM0079.1) | 3.8% | 3.4% | 3.0% | n.s | $p < 10^{-6}$ * | n.s |
| BHLHE40 (TFFM0007.1) | 13.1% | 14.5% | 14.1% | $p < 10^{-6}$ * | 0.001466 | n.s |
| RFX1 (TFFM0089.1) | 5.6% | 6.6% | 6.0% | $1.10 \times 10^{-5}$ * | n.s | 0.028718 |
| TRPS1 (TFFM0980.1) | 22.1% | 23.7% | 23.8% | $2.30 \times 10^{-5}$ * | $1.40 \times 10^{-5}$ * | n.s |
| CREB1 (TFFM0705.1) | 8.0% | 9.1% | 8.8% | $1.50 \times 10^{-5}$ * | 0.000364 | n.s |
| IRF9 (TFFM0757.1) | 6.3% | 5.4% | 5.4% | $2.10 \times 10^{-5}$ * | $1.50 \times 10^{-5}$ * | n.s |
| MSGN1 (TFFM0862.1) | 9.7% | 8.6% | 9.6% | $1.50 \times 10^{-5}$ * | n.s | 0.003044 |
| TFAP2A (TFFM0111.2) | 21.0% | 19.5% | 20.0% | $1.50 \times 10^{-5}$ * | n.s | n.s |
| ESRRA (TFFM0028.1) | 5.7% | 6.7% | 5.8% | $1.60 \times 10^{-5}$ * | n.s | 0.001858 |
| HOXB13 (TFFM0180.2) | 2.2% | 2.0% | 1.7% | n.s | $1.70 \times 10^{-5}$ * | n.s |
| GATA1 (TFFM0040.1) | 3.7% | 4.5% | 4.2% | $1.80 \times 10^{-5}$ * | n.s | n.s |
| TCF4 (TFFM0601.2) | 23.2% | 21.7% | 21.6% | $7.30 \times 10^{-5}$ * | $1.80 \times 10^{-5}$ * | n.s |
| MZF1 (TFFM0531.2) | 5.2% | 6.1% | 5.4% | $1.90 \times 10^{-5}$ * | n.s | 0.007048 |
| MAFK (TFFM0058.3) | 8.5% | 7.5% | 7.5% | $2.10 \times 10^{-5}$ * | $5.30 \times 10^{-5}$ * | n.s |
| THRB (TFFM0885.1) | 2.0% | 1.9% | 1.5% | n.s | $2.10 \times 10^{-5}$ * | n.s |
| KLF4 (TFFM0056.2) | 35.1% | 35.3% | 33.3% | n.s | $2.20 \times 10^{-5}$ * | 0.000394 |
| USF2 (TFFM0123.1) | 12.3% | 13.5% | 12.8% | $2.40 \times 10^{-5}$ * | n.s | 0.032162 |
| KLF6 (TFFM0518.1) | 19.4% | 20.9% | 20.5% | $2.50 \times 10^{-5}$ * | 0.001505 | n.s |
| ZNF317 (TFFM0639.1) | 3.4% | 3.8% | 4.1% | n.s | $2.60 \times 10^{-5}$ * | n.s |

Continued on next page

Table S3 – continued from previous page

| TFBS | type 0 | UHP | DHP | 0 vs. UHP | 0 vs DHP | UHP vs. DHP |
|---|---|---|---|---|---|---|
| OVOL2 (TFFM0548.1) | 1.4% | 1.9% | 1.3% | n.s | n.s | $2.70 \times 10^{-5}$ * |
| RFX1 (TFFM0089.2) | 6.5% | 7.5% | 7.1% | $2.70 \times 10^{-5}$ * | 0.008550 | n.s |
| AR (TFFM0001.1) | 20.1% | 18.7% | 19.8% | $2.80 \times 10^{-5}$ * | n.s | 0.011240 |
| JUNB (TFFM0829.1) | 1.8% | 2.3% | 2.2% | $2.90 \times 10^{-5}$ * | n.s | n.s |
| MITF (TFFM0141.1) | 10.8% | 12.0% | 11.3% | $3.50 \times 10^{-5}$ * | n.s | 0.043905 |
| TBX20 (TFFM0766.1) | 10.0% | 10.1% | 8.9% | n.s | $3.50 \times 10^{-5}$ * | 0.000815 |
| USF1 (TFFM0122.1) | 14.1% | 15.4% | 14.7% | $3.50 \times 10^{-5}$ * | n.s | n.s |
| NR3C1 (TFFM0080.1) | 6.7% | 5.8% | 5.9% | $3.70 \times 10^{-5}$ * | $5.90 \times 10^{-5}$ * | n.s |
| HSF2 (TFFM0786.1) | 6.9% | 7.8% | 7.8% | $9.60 \times 10^{-5}$ | $3.90 \times 10^{-5}$ * | n.s |
| PRDM1 (TFFM0087.2) | 4.9% | 4.9% | 4.2% | n.s | $4.10 \times 10^{-5}$ * | 0.002962 |
| HNF1A (TFFM0989.1) | 16.8% | 17.3% | 18.1% | n.s | $4.30 \times 10^{-5}$ * | n.s |
| OTX2 (TFFM0197.1) | 3.2% | 2.7% | 2.6% | n.s | $4.40 \times 10^{-5}$ * | n.s |
| ERF (TFFM0476.1) | 17.2% | 15.9% | 16.8% | $4.70 \times 10^{-5}$ * | n.s | 0.021609 |
| KLF1 (TFFM0055.1) | 19.5% | 20.4% | 18.6% | n.s | n.s | $4.90 \times 10^{-5}$ * |
| ZBTB7B (TFFM0770.1) | 5.3% | 6.1% | 5.8% | $5.40 \times 10^{-5}$ * | 0.007816 | n.s |
| STAT3 (TFFM0102.1) | 10.4% | 10.8% | 9.3% | n.s | 0.000102 | $6.30 \times 10^{-5}$ * |
| KLF4 (TFFM0056.3) | 36.7% | 38.0% | 35.8% | n.s | n.s | $6.80 \times 10^{-5}$ * |
| IRF2 (TFFM0181.1) | 3.6% | 3.4% | 3.0% | n.s | $6.90 \times 10^{-5}$ * | n.s |
| MAFG (TFFM0188.2) | 5.7% | 4.9% | 5.0% | $7.00 \times 10^{-5}$ * | 0.001299 | n.s |
| NR5A2 (TFFM0731.1) | 2.9% | 2.4% | 2.4% | 0.000255 | $7.90 \times 10^{-5}$ * | n.s |
| MAZ (TFFM0524.1) | 10.1% | 11.2% | 10.8% | $8.00 \times 10^{-5}$ * | 0.011105 | n.s |
| ESRRB (TFFM0029.1) | 3.8% | 3.2% | 3.3% | $8.40 \times 10^{-5}$ * | 0.000604 | n.s |
| MAFF (TFFM0057.2) | 4.8% | 4.8% | 5.6% | n.s | $9.10 \times 10^{-5}$ * | 0.000954 |
| ATF7 (TFFM0164.1) | 6.1% | 6.9% | 6.5% | $9.80 \times 10^{-5}$ | n.s | n.s |
| CREB1 (TFFM0012.1) | 16.7% | 17.7% | 18.0% | 0.002328 | $9.80 \times 10^{-5}$ | n.s |
| CEBPG (TFFM0893.1) | 7.7% | 8.3% | 8.7% | 0.010044 | 0.000103 | n.s |
| LHX2 (TFFM0185.2) | 3.6% | 3.0% | 3.3% | 0.000104 | n.s | n.s |
| CDX2 (TFFM0008.2) | 4.1% | 4.8% | 4.6% | 0.000110 | n.s | n.s |
| ZEB1 (TFFM0127.1) | 26.7% | 25.2% | 25.8% | 0.000112 | n.s | n.s |
| TFAP2C (TFFM0793.1) | 21.7% | 22.8% | 23.1% | n.s | 0.000114 | n.s |
| SCRT1 (TFFM0580.2) | 10.0% | 9.1% | 9.0% | 0.000356 | 0.000116 | n.s |
| RELB (TFFM0575.1) | 4.3% | 3.8% | 3.7% | 0.002573 | 0.000125 | n.s |
| FOXO3 (TFFM0488.1) | 13.3% | 12.2% | 12.2% | 0.000148 | 0.000258 | n.s |
| ZKSCAN1 (TFFM0630.1) | 2.5% | 2.9% | 3.0% | n.s | 0.000150 | n.s |
| SPI1 (TFFM0099.2) | 13.9% | 12.7% | 14.0% | 0.000157 | n.s | 0.001355 |
| KLF12 (TFFM0514.1) | 26.2% | 26.7% | 24.8% | n.s | 0.000159 | 0.000221 |
| HOXD13 (TFFM0808.1) | 8.2% | 8.9% | 7.7% | 0.004871 | 0.015241 | 0.000164 |
| NEUROD2 (TFFM0990.1) | 10.2% | 10.3% | 11.2% | n.s | 0.000170 | 0.007087 |
| ERG (TFFM0725.1) | 21.6% | 20.3% | 21.7% | 0.000175 | n.s | 0.002476 |
| THAP11 (TFFM0883.1) | 13.4% | 12.3% | 13.1% | 0.000185 | n.s | 0.020255 |
| MYF5 (TFFM0676.1) | 17.3% | 16.1% | 16.4% | 0.000191 | n.s | n.s |
| ZSCAN29 (TFFM0648.1) | 8.1% | 7.3% | 7.3% | 0.000307 | 0.000199 | n.s |
| JUN (TFFM0050.1) | 10.9% | 12.0% | 11.4% | 0.000214 | n.s | n.s |
| SRF (TFFM0100.1) | 4.7% | 5.4% | 4.5% | 0.000215 | n.s | 0.000642 |

Continued on next page

Table S3 – continued from previous page

| TFBS | type 0 | UHP | DHP | 0 vs. UHP | 0 vs DHP | UHP vs. DHP |
|---|---|---|---|---|---|---|
| EGR1 (TFFM0020.2) | 23.0% | 24.4% | 23.0% | 0.000219 | n.s | 0.002962 |
| GATA4 (TFFM0043.1) | 4.7% | 5.5% | 5.3% | 0.000221 | 0.001686 | n.s |
| PAX5 (TFFM0084.2) | 17.6% | 18.9% | 18.5% | 0.000228 | 0.005266 | n.s |
| ZNF692 (TFFM0984.1) | 14.5% | 14.2% | 13.4% | n.s | 0.000228 | 0.030029 |
| GATA6 (TFFM0137.2) | 5.6% | 6.4% | 6.2% | 0.000246 | 0.003164 | n.s |
| CTCFL (TFFM0133.1) | 22.6% | 23.9% | 23.9% | 0.000345 | 0.000255 | n.s |
| USF2 (TFFM0738.1) | 9.6% | 10.6% | 10.4% | 0.000258 | 0.003336 | n.s |
| ZIC2 (TFFM0628.1) | 3.0% | 2.4% | 2.5% | 0.000260 | n.s | n.s |
| NFIC (TFFM0863.1) | 8.7% | 9.3% | 8.1% | 0.007537 | 0.018181 | 0.000281 |
| NFE2L2 (TFFM0071.1) | 6.9% | 7.7% | 7.6% | 0.000285 | 0.000627 | n.s |
| RUNX3 (TFFM0093.1) | 5.5% | 5.9% | 5.0% | n.s | 0.006045 | 0.000297 |
| SP1 (TFFM0097.1) | 23.5% | 24.3% | 22.5% | n.s | n.s | 0.000302 |
| ZBTB7A (TFFM0126.2) | 22.1% | 21.4% | 20.9% | n.s | 0.000310 | n.s |
| MAFK (TFFM0058.2) | 5.7% | 5.1% | 6.0% | 0.001346 | n.s | 0.000328 |
| CTCF (TFFM0461.1) | 20.5% | 21.8% | 20.2% | 0.000337 | n.s | 0.000580 |
| BHLHA15 (TFFM0856.1) | 7.7% | 6.9% | 7.1% | 0.000339 | 0.004142 | n.s |
| KLF5 (TFFM0183.1) | 26.3% | 26.4% | 24.9% | n.s | 0.000345 | 0.003945 |
| NFATC1 (TFFM0751.1) | 6.8% | 7.6% | 7.1% | 0.000352 | n.s | 0.036032 |
| NR6A1 (TFFM0544.1) | 7.9% | 7.0% | 7.1% | 0.000352 | 0.000794 | n.s |
| TFAP2C (TFFM0117.1) | 13.8% | 12.8% | 13.8% | 0.000355 | n.s | 0.005043 |
| STAT6 (TFFM0105.1) | 9.3% | 10.3% | 9.9% | 0.000395 | 0.018147 | n.s |
| ZNF257 (TFFM0909.1) | 19.8% | 21.0% | 20.6% | 0.000395 | n.s | n.s |
| SOX17 (TFFM0587.1) | 16.0% | 15.6% | 14.9% | n.s | 0.000399 | n.s |
| IRF1 (TFFM0708.1) | 3.8% | 3.5% | 3.2% | n.s | 0.000404 | n.s |
| MAFB (TFFM0187.1) | 4.8% | 5.1% | 5.5% | n.s | 0.000430 | n.s |
| MYOD1 (TFFM0068.2) | 14.0% | 12.9% | 13.4% | 0.000431 | n.s | n.s |
| TBX19 (TFFM0597.1) | 15.8% | 14.7% | 14.7% | 0.001034 | 0.000437 | n.s |
| MAFG (TFFM0188.1) | 6.0% | 5.9% | 6.8% | n.s | 0.000453 | 0.001107 |
| MXI1 (TFFM0142.1) | 6.6% | 7.4% | 7.2% | 0.000481 | 0.007622 | n.s |
| NR2C1 (TFFM0542.1) | 4.2% | 4.3% | 3.6% | n.s | 0.000486 | 0.003941 |
| ZNF549 (TFFM0921.1) | 20.2% | 21.3% | 21.4% | 0.002466 | 0.000489 | n.s |
| SPDEF (TFFM0592.1) | 8.2% | 8.1% | 7.4% | n.s | 0.000528 | 0.019320 |
| TCF7 (TFFM0209.2) | 6.4% | 6.8% | 7.1% | n.s | 0.000531 | n.s |
| RFX3 (TFFM0577.1) | 7.2% | 8.0% | 7.7% | 0.000532 | 0.010058 | n.s |
| TBP (TFFM0106.1) | 5.3% | 5.9% | 6.0% | 0.007525 | 0.000569 | n.s |
| PTF1A (TFFM0567.1) | 17.6% | 16.7% | 16.5% | n.s | 0.000584 | n.s |
| ZNF189 (TFFM0918.1) | 19.7% | 20.1% | 18.6% | n.s | 0.000700 | 0.000628 |
| MAFF (TFFM0057.1) | 12.6% | 11.6% | 12.3% | 0.000651 | n.s | 0.042332 |
| HOXD13 (TFFM0506.1) | 8.8% | 9.7% | 8.9% | 0.000717 | n.s | 0.016009 |
| SCRT1 (TFFM0580.1) | 9.9% | 9.1% | 9.0% | 0.001167 | 0.000721 | n.s |
| SREBF1 (TFFM0797.1) | 5.3% | 5.1% | 5.9% | n.s | 0.000730 | 0.002175 |
| FOXJ2 (TFFM0175.1) | 14.1% | 13.6% | 13.1% | n.s | 0.000747 | n.s |
| SNAI2 (TFFM0203.2) | 6.0% | 5.3% | 5.8% | 0.000762 | n.s | n.s |
| RBPJ (TFFM0149.1) | 6.0% | 5.3% | 5.7% | 0.000772 | n.s | n.s |

Continued on next page

Table S3 – continued from previous page

| TFBS | type 0 | UHP | DHP | 0 vs. UHP | 0 vs DHP | UHP vs. DHP |
|---|---|---|---|---|---|---|
| USF2 (TFFM0123.3) | 10.3% | 11.1% | 11.2% | 0.002157 | 0.000773 | n.s |
| ZBTB26 (TFFM0623.1) | 18.7% | 18.5% | 19.8% | n.s | 0.000785 | 0.002750 |
| GATA2 (TFFM0041.2) | 3.8% | 4.4% | 4.4% | 0.000835 | 0.001841 | n.s |
| NFATC2 (TFFM0720.1) | 19.0% | 19.1% | 20.1% | n.s | 0.000911 | 0.017779 |
| PKNOX1 (TFFM0560.2) | 4.2% | 3.7% | 4.5% | n.s | n.s | 0.000981 |
| MAF (TFFM0523.1) | 24.1% | 22.9% | 23.8% | 0.000984 | n.s | n.s |
| ZNF281 (TFFM0637.1) | 18.3% | 19.4% | 19.3% | 0.001003 | 0.001663 | n.s |
| TBX5 (TFFM0599.1) | 4.5% | 5.1% | 4.6% | 0.001016 | n.s | 0.017567 |
| MEIS1 (TFFM0894.1) | 5.1% | 4.5% | 4.7% | 0.001044 | n.s | n.s |
| ZNF652 (TFFM0702.1) | 5.3% | 5.9% | 5.7% | 0.001052 | n.s | n.s |
| TFAP2C (TFFM0118.1) | 16.4% | 17.5% | 17.0% | 0.001085 | n.s | n.s |
| SOX10 (TFFM0152.1) | 13.0% | 12.1% | 12.1% | 0.001363 | 0.001096 | n.s |
| TP63 (TFFM0120.1) | 8.0% | 8.8% | 8.2% | 0.001129 | n.s | 0.030882 |
| MEF2B (TFFM0189.1) | 12.1% | 13.0% | 12.2% | 0.001134 | n.s | 0.025587 |
| NR2F1 (TFFM0194.1) | 3.5% | 3.6% | 3.0% | n.s | 0.001255 | 0.008668 |
| ELF1 (TFFM0022.1) | 18.7% | 17.6% | 17.8% | 0.001298 | n.s | n.s |
| NFYA (TFFM0074.1) | 3.2% | 2.9% | 3.5% | n.s | n.s | 0.001303 |
| ZNF449 (TFFM0701.1) | 19.2% | 19.7% | 18.2% | n.s | n.s | 0.001344 |
| EBF1 (TFFM0019.2) | 4.9% | 4.5% | 5.2% | n.s | n.s | 0.001498 |
| MLX (TFFM0527.1) | 4.4% | 4.5% | 5.0% | n.s | 0.001559 | n.s |
| ZBTB18 (TFFM0622.1) | 14.2% | 13.2% | 13.6% | 0.001605 | n.s | n.s |
| ISL1 (TFFM0512.1) | 13.8% | 12.9% | 13.8% | 0.001675 | n.s | 0.012928 |
| HNF4G (TFFM0046.1) | 4.0% | 3.4% | 3.8% | 0.001676 | n.s | n.s |
| ELK4 (TFFM0024.1) | 13.2% | 12.9% | 14.1% | n.s | 0.003207 | 0.001687 |
| NEUROD2 (TFFM0760.1) | 14.7% | 13.7% | 14.6% | 0.001714 | n.s | 0.028355 |
| GATA1 (TFFM0040.2) | 3.8% | 4.4% | 4.2% | 0.001775 | n.s | n.s |
| PRDM1 (TFFM0087.3) | 9.6% | 10.1% | 9.0% | 0.028021 | 0.026310 | 0.001798 |
| ELF5 (TFFM0718.1) | 17.0% | 16.0% | 16.2% | 0.001855 | n.s | n.s |
| SMAD4 (TFFM0151.1) | 3.7% | 4.0% | 4.2% | n.s | 0.001878 | n.s |
| SP9 (TFFM0878.1) | 3.9% | 3.4% | 3.8% | 0.001881 | n.s | n.s |
| ZNF343 (TFFM0910.1) | 12.6% | 13.5% | 13.3% | 0.001959 | 0.005886 | n.s |
| NEUROD2 (TFFM0533.1) | 13.3% | 12.4% | 13.1% | 0.002018 | n.s | 0.047582 |
| POU2F2 (TFFM0085.1) | 12.4% | 11.9% | 11.6% | n.s | 0.002080 | n.s |
| TCF12 (TFFM0107.1) | 15.5% | 14.6% | 14.7% | 0.002230 | 0.006327 | n.s |
| FOXJ3 (TFFM0801.1) | 9.9% | 9.3% | 9.2% | 0.011124 | 0.002298 | n.s |
| EHF (TFFM0471.2) | 17.9% | 17.6% | 18.9% | n.s | 0.002397 | 0.003895 |
| OLIG2 (TFFM0763.1) | 6.0% | 5.5% | 5.4% | 0.008476 | 0.002543 | n.s |
| YY2 (TFFM0621.1) | 11.5% | 12.3% | 11.8% | 0.002661 | n.s | n.s |
| RFX5 (TFFM0090.1) | 6.2% | 6.8% | 6.8% | 0.002692 | 0.004805 | n.s |
| CREB5 (TFFM0800.1) | 4.3% | 4.8% | 4.7% | 0.002748 | n.s | n.s |
| TFAP2B (TFFM0115.1) | 15.4% | 15.1% | 16.3% | n.s | 0.004641 | 0.002827 |
| MAFG (TFFM0759.1) | 10.8% | 10.0% | 10.5% | 0.002829 | n.s | n.s |
| MEIS1 (TFFM0062.1) | 10.5% | 11.3% | 10.9% | 0.002831 | n.s | n.s |
| TCF3 (TFFM0737.1) | 11.9% | 11.4% | 11.1% | n.s | 0.002893 | n.s |

Continued on next page

| TFBS | type 0 | UHP | DHP | 0 vs. UHP | 0 vs DHP | UHP vs. DHP |
|---|---|---|---|---|---|---|
| SP2 (TFFM0098.2) | 15.6% | 14.6% | 14.8% | 0.003062 | 0.009185 | n.s |
| ELF5 (TFFM0473.1) | 24.0% | 23.0% | 24.4% | n.s | n.s | 0.003274 |
| TBX6 (TFFM0880.1) | 12.1% | 12.5% | 13.0% | n.s | 0.003319 | n.s |
| PTF1A (TFFM0888.1) | 9.6% | 9.0% | 8.9% | 0.011872 | 0.003373 | n.s |
| ELF3 (TFFM0170.1) | 12.8% | 12.0% | 12.0% | 0.003439 | 0.004782 | n.s |
| TFCP2 (TFFM0604.1) | 8.8% | 9.2% | 9.6% | n.s | 0.003589 | n.s |
| REL (TFFM0715.1) | 13.5% | 13.8% | 14.4% | n.s | 0.003612 | n.s |
| WT1 (TFFM0620.1) | 4.3% | 4.6% | 3.9% | n.s | n.s | 0.004064 |
| ERG (TFFM0025.1) | 24.0% | 22.8% | 24.2% | n.s | n.s | 0.004167 |
| POU2F1 (TFFM0788.1) | 13.6% | 12.7% | 13.2% | 0.004268 | n.s | n.s |
| SOX17 (TFFM0711.1) | 4.9% | 4.7% | 5.4% | n.s | n.s | 0.004549 |
| FOXA1 (TFFM0035.1) | 11.7% | 11.2% | 10.9% | n.s | 0.004596 | n.s |
| DLX5 (TFFM0857.1) | 7.2% | 7.0% | 6.6% | n.s | 0.004615 | n.s |
| ETV2 (TFFM0479.1) | 25.0% | 23.8% | 25.2% | n.s | n.s | 0.004648 |
| CEBPD (TFFM0011.1) | 13.3% | 12.9% | 12.5% | n.s | 0.005008 | n.s |
| POU5F1 (TFFM0148.1) | 10.6% | 11.3% | 11.2% | 0.005319 | 0.012313 | n.s |
| TEAD1 (TFFM0210.1) | 6.0% | 5.6% | 6.3% | n.s | n.s | 0.006337 |
| MYCN (TFFM0067.2) | 12.8% | 13.6% | 13.5% | 0.006348 | 0.014029 | n.s |
| ZBTB7A (TFFM0126.1) | 5.3% | 5.9% | 5.7% | 0.006676 | n.s | n.s |
| PKNOX1 (TFFM0560.1) | 5.3% | 4.8% | 5.4% | 0.006826 | n.s | 0.008376 |
| OLIG2 (TFFM0991.1) | 3.8% | 3.6% | 4.2% | n.s | n.s | 0.006938 |
| BCL6 (TFFM0006.2) | 7.0% | 7.6% | 7.0% | 0.007097 | n.s | 0.029799 |
| GFI1B (TFFM0044.1) | 7.9% | 7.3% | 7.3% | 0.008345 | 0.007182 | n.s |
| MYCN (TFFM0067.1) | 8.5% | 7.9% | 8.0% | 0.007806 | 0.017610 | n.s |
| RFX3 (TFFM0577.2) | 8.4% | 9.0% | 8.4% | 0.007861 | n.s | 0.028152 |
| CRX (TFFM0013.1) | 10.8% | 11.5% | 11.1% | 0.008212 | n.s | n.s |
| CRX (TFFM0723.1) | 13.6% | 14.4% | 13.8% | 0.008256 | n.s | n.s |
| NFIL3 (TFFM0539.1) | 10.3% | 11.0% | 10.7% | 0.008745 | n.s | n.s |
| BCL6B (TFFM0447.1) | 11.6% | 11.3% | 10.9% | n.s | 0.009060 | n.s |
| TFAP2B (TFFM0116.1) | 14.7% | 15.3% | 15.5% | n.s | 0.009219 | n.s |
| ZNF136 (TFFM0633.1) | 5.5% | 6.0% | 5.6% | 0.009226 | n.s | n.s |
| GATA2 (TFFM0041.1) | 5.7% | 6.3% | 6.2% | 0.009236 | n.s | n.s |
| ELK1 (TFFM0023.1) | 13.6% | 14.0% | 14.4% | n.s | 0.009694 | n.s |
| TCF21 (TFFM0600.1) | 15.5% | 14.7% | 15.6% | 0.010412 | n.s | 0.019447 |
| BHLHE41 (TFFM0450.1) | 11.7% | 12.4% | 12.2% | 0.010570 | n.s | n.s |
| MYOD1 (TFFM0068.1) | 11.6% | 10.9% | 11.2% | 0.010642 | n.s | n.s |
| ZNF331 (TFFM0919.1) | 11.1% | 10.5% | 11.4% | 0.021488 | n.s | 0.011104 |
| MXI1 (TFFM0142.2) | 5.9% | 6.4% | 6.2% | 0.011695 | n.s | n.s |
| ATOH1 (TFFM0854.1) | 10.5% | 9.9% | 9.9% | 0.012599 | 0.012021 | n.s |
| NFYC (TFFM0682.1) | 5.1% | 4.9% | 5.5% | n.s | n.s | 0.013208 |
| USF1 (TFFM0122.2) | 11.7% | 12.4% | 12.1% | 0.013422 | n.s | n.s |
| PBX2 (TFFM0146.1) | 5.7% | 5.3% | 5.9% | n.s | n.s | 0.013657 |
| RFX2 (TFFM0576.1) | 7.0% | 7.5% | 7.5% | 0.013825 | n.s | n.s |
| NFKB2 (TFFM0193.1) | 9.9% | 10.5% | 9.7% | 0.015750 | n.s | 0.013931 |

Continued on next page

Table S3 – continued from previous page

| TFBS | type 0 | UHP | DHP | 0 vs. UHP | 0 vs DHP | UHP vs. DHP |
|---|---|---|---|---|---|---|
| BACH2 (TFFM0855.1) | 7.4% | 7.1% | 6.9% | n.s | 0.014747 | n.s |
| THRB (TFFM0609.1) | 10.4% | 9.8% | 10.1% | 0.015256 | n.s | n.s |
| NFATC1 (TFFM0535.1) | 13.7% | 14.4% | 14.2% | 0.016400 | n.s | n.s |
| HOXA13 (TFFM0756.1) | 5.9% | 6.1% | 5.5% | n.s | n.s | 0.016432 |
| NFYB (TFFM0075.2) | 7.0% | 7.1% | 7.5% | n.s | 0.017033 | n.s |
| HEY2 (TFFM0755.1) | 8.9% | 9.1% | 9.4% | n.s | 0.017042 | n.s |
| NR2F6 (TFFM0776.1) | 15.8% | 16.3% | 15.4% | n.s | n.s | 0.019455 |
| GRHL2 (TFFM0138.1) | 9.1% | 9.7% | 9.3% | 0.020826 | n.s | n.s |
| TP73 (TFFM0121.1) | 11.0% | 11.2% | 10.5% | n.s | n.s | 0.030064 |
| SOX2 (TFFM0095.1) | 8.8% | 8.4% | 9.1% | n.s | n.s | 0.030714 |
| NFE2L1 (TFFM0192.1) | 15.4% | 15.2% | 16.1% | n.s | n.s | 0.032053 |
| ATOH1 (TFFM0004.1) | 11.9% | 11.6% | 12.3% | n.s | n.s | 0.032501 |
| NR2F2 (TFFM0144.1) | 15.5% | 15.6% | 14.8% | n.s | n.s | 0.035768 |
| EOMES (TFFM0171.1) | 12.0% | 11.5% | 12.2% | n.s | n.s | 0.036098 |
| IRF5 (TFFM0852.1) | 11.2% | 11.0% | 11.7% | n.s | n.s | 0.039727 |
| TFAP4 (TFFM0212.1) | 9.7% | 9.4% | 10.0% | n.s | n.s | 0.049575 |
| TWIST1 (TFFM0155.2) | 10.2% | 9.7% | 10.3% | n.s | n.s | 0.08 |
| ARNT (TFFM0161.1) | 6.6% | 6.7% | 6.5% | n.s | n.s | n.s |
| ATF3 (TFFM0003.1) | 11.3% | 11.8% | 11.6% | n.s | n.s | n.s |
| ATF4 (TFFM0163.1) | 6.0% | 6.4% | 6.1% | n.s | n.s | n.s |
| BACH1 (TFFM0654.1) | 4.7% | 5.0% | 4.7% | n.s | n.s | n.s |
| BACH1 (TFFM0891.1) | 3.9% | 4.1% | 4.0% | n.s | n.s | n.s |
| BACH2 (TFFM0132.1) | 6.8% | 6.7% | 6.6% | n.s | n.s | n.s |
| BACH2 (TFFM0132.2) | 11.8% | 11.4% | 11.6% | n.s | n.s | n.s |
| BATF3 (TFFM0446.1) | 0.5% | 0.3% | 0.5% | n.s | n.s | n.s |
| BATF3 (TFFM0656.1) | 5.9% | 5.7% | 5.6% | n.s | n.s | n.s |
| BATF (TFFM0655.1) | 6.7% | 6.6% | 6.8% | n.s | n.s | n.s |
| BHLHA15 (TFFM0449.1) | 13.1% | 12.5% | 13.0% | n.s | n.s | n.s |
| BHLHA15 (TFFM0745.1) | 6.5% | 6.1% | 6.2% | n.s | n.s | n.s |
| CDX2 (TFFM0008.1) | 2.6% | 2.3% | 2.4% | n.s | n.s | n.s |
| CEBPA (TFFM0009.1) | 8.9% | 9.0% | 8.9% | n.s | n.s | n.s |
| CEBPA (TFFM0009.2) | 9.9% | 10.1% | 10.4% | n.s | n.s | n.s |
| CEBPB (TFFM0010.1) | 8.8% | 9.1% | 8.8% | n.s | n.s | n.s |
| CEBPB (TFFM0722.1) | 6.9% | 7.0% | 6.9% | n.s | n.s | n.s |
| CEBPD (TFFM0011.2) | 10.0% | 10.1% | 10.3% | n.s | n.s | n.s |
| CEBPE (TFFM0455.1) | 13.1% | 12.9% | 12.7% | n.s | n.s | n.s |
| CEBPE (TFFM0799.1) | 10.0% | 10.2% | 9.9% | n.s | n.s | n.s |
| CEBPG (TFFM0166.1) | 7.3% | 7.5% | 7.2% | n.s | n.s | n.s |
| CLOCK (TFFM0459.1) | 12.5% | 12.3% | 12.0% | n.s | n.s | n.s |
| CLOCK (TFFM0795.1) | 1.3% | 1.2% | 1.2% | n.s | n.s | n.s |
| CREB3L1 (TFFM0167.1) | 8.3% | 7.9% | 8.3% | n.s | n.s | n.s |
| CREB3L2 (TFFM0746.1) | 4.3% | 4.4% | 4.3% | n.s | n.s | n.s |
| CREM (TFFM0168.1) | 13.7% | 14.3% | 13.9% | n.s | n.s | n.s |
| CREM (TFFM0747.1) | 7.8% | 7.8% | 7.8% | n.s | n.s | n.s |

Continued on next page

**Table S3 – continued from previous page**

| TFBS | type 0 | UHP | DHP | 0 vs. UHP | 0 vs DHP | UHP vs. DHP |
|---|---|---|---|---|---|---|
| CUX1 (TFFM0169.1) | 2.2% | 2.0% | 2.1% | n.s | n.s | n.s |
| CUX1 (TFFM0781.1) | 8.6% | 8.6% | 8.5% | n.s | n.s | n.s |
| CUX2 (TFFM0782.1) | 3.3% | 3.2% | 3.1% | n.s | n.s | n.s |
| DLX1 (TFFM0804.1) | 10.7% | 10.2% | 10.3% | n.s | n.s | n.s |
| DLX2 (TFFM0805.1) | 7.9% | 7.8% | 7.6% | n.s | n.s | n.s |
| DMRT1 (TFFM0464.1) | 7.7% | 8.0% | 7.7% | n.s | n.s | n.s |
| DUX4 (TFFM0015.1) | 2.2% | 2.0% | 2.1% | n.s | n.s | n.s |
| E2F4 (TFFM0017.1) | 38.2% | 40.1% | 39.9% | n.s | n.s | n.s |
| E2F4 (TFFM0017.2) | 36.1% | 37.4% | 36.8% | n.s | n.s | n.s |
| E2F6 (TFFM0018.1) | 10.3% | 10.5% | 10.6% | n.s | n.s | n.s |
| E2F6 (TFFM0724.1) | 11.0% | 11.0% | 10.5% | n.s | n.s | n.s |
| EBF1 (TFFM0019.1) | 13.9% | 13.5% | 13.3% | n.s | n.s | n.s |
| EBF3 (TFFM0662.1) | 24.8% | 25.0% | 24.7% | n.s | n.s | n.s |
| EGR1 (TFFM0020.1) | 23.4% | 24.1% | 23.0% | n.s | n.s | n.s |
| EGR1 (TFFM0020.3) | 16.8% | 16.7% | 16.3% | n.s | n.s | n.s |
| EGR2 (TFFM0021.1) | 15.8% | 16.4% | 16.3% | n.s | n.s | n.s |
| EHF (TFFM0471.1) | 17.4% | 17.4% | 17.2% | n.s | n.s | n.s |
| ELK3 (TFFM0474.1) | 13.6% | 13.3% | 13.4% | n.s | n.s | n.s |
| ESR1 (TFFM0026.1) | 29.9% | 30.0% | 30.4% | n.s | n.s | n.s |
| ESR2 (TFFM0027.1) | 23.7% | 23.5% | 23.0% | n.s | n.s | n.s |
| ESRRA (TFFM0028.2) | 1.9% | 1.8% | 1.6% | n.s | n.s | n.s |
| ETS1 (TFFM0030.1) | 22.2% | 21.1% | 22.1% | n.s | n.s | n.s |
| ETV1 (TFFM0172.1) | 20.6% | 19.9% | 20.3% | n.s | n.s | n.s |
| ETV1 (TFFM0172.2) | 35.5% | 37.2% | 36.0% | n.s | n.s | n.s |
| ETV5 (TFFM0480.2) | 9.5% | 10.0% | 9.8% | n.s | n.s | n.s |
| FOS (TFFM0032.1) | 8.0% | 8.1% | 8.1% | n.s | n.s | n.s |
| FOSL1 (TFFM0033.1) | 5.1% | 5.4% | 5.4% | n.s | n.s | n.s |
| FOSL1 (TFFM0033.2) | 5.3% | 5.6% | 5.5% | n.s | n.s | n.s |
| FOSL2 (TFFM0034.1) | 6.5% | 6.5% | 6.3% | n.s | n.s | n.s |
| FOXA1 (TFFM0035.2) | 3.0% | 2.8% | 2.7% | n.s | n.s | n.s |
| FOXA2 (TFFM0036.2) | 7.5% | 7.3% | 7.2% | n.s | n.s | n.s |
| FOXA3 (TFFM0667.1) | 10.8% | 10.3% | 10.6% | n.s | n.s | n.s |
| FOXF1 (TFFM0486.1) | 11.6% | 11.8% | 12.2% | n.s | n.s | n.s |
| FOXF2 (TFFM0706.1) | 10.8% | 11.0% | 11.3% | n.s | n.s | n.s |
| FOXG1 (TFFM0749.1) | 12.9% | 12.3% | 12.7% | n.s | n.s | n.s |
| FOXK1 (TFFM0134.1) | 21.4% | 21.4% | 20.9% | n.s | n.s | n.s |
| FOXK2 (TFFM0135.1) | 6.4% | 6.3% | 6.2% | n.s | n.s | n.s |
| FOXK2 (TFFM0135.2) | 5.0% | 5.0% | 4.8% | n.s | n.s | n.s |
| FOXO1 (TFFM0038.1) | 7.4% | 7.6% | 7.7% | n.s | n.s | n.s |
| FOXO3 (TFFM0721.1) | 8.7% | 9.0% | 8.8% | n.s | n.s | n.s |
| FOXP1 (TFFM0136.1) | 3.2% | 3.1% | 3.0% | n.s | n.s | n.s |
| FOXP1 (TFFM0726.1) | 4.4% | 4.9% | 4.8% | n.s | n.s | n.s |
| GABPA (TFFM0039.1) | 17.1% | 16.7% | 17.4% | n.s | n.s | n.s |
| GATA3 (TFFM0042.1) | 1.9% | 1.8% | 1.8% | n.s | n.s | n.s |

Continued on next page

Table S3 – continued from previous page

| TFBS | type 0 | UHP | DHP | 0 vs. UHP | 0 vs DHP | UHP vs. DHP |
|---|---|---|---|---|---|---|
| GATA3 (TFFM0042.2) | 3.4% | 3.4% | 3.3% | n.s | n.s | n.s |
| GATA3 (TFFM0707.1) | 3.0% | 3.0% | 3.0% | n.s | n.s | n.s |
| GATA4 (TFFM0043.2) | 3.0% | 3.3% | 3.3% | n.s | n.s | n.s |
| GATA6 (TFFM0137.1) | 4.4% | 4.7% | 4.5% | n.s | n.s | n.s |
| GFI1 (TFFM0491.1) | 3.7% | 3.3% | 3.3% | n.s | n.s | n.s |
| GLI2 (TFFM0778.1) | 39.1% | 39.7% | 40.2% | n.s | n.s | n.s |
| GLI3 (TFFM0859.1) | 0.5% | 0.5% | 0.4% | n.s | n.s | n.s |
| GLIS2 (TFFM0493.1) | 0.3% | 0.2% | 0.2% | n.s | n.s | n.s |
| GLIS3 (TFFM0779.1) | 2.3% | 1.8% | 1.9% | n.s | n.s | n.s |
| GMEB1 (TFFM0750.1) | 12.0% | 12.0% | 11.6% | n.s | n.s | n.s |
| GRHL1 (TFFM0754.1) | 8.1% | 8.0% | 8.5% | n.s | n.s | n.s |
| GRHL2 (TFFM0138.2) | 9.3% | 9.3% | 9.3% | n.s | n.s | n.s |
| HES1 (TFFM0826.1) | 0.5% | 0.7% | 0.8% | n.s | n.s | n.s |
| HEY1 (TFFM0796.1) | 1.7% | 1.6% | 1.6% | n.s | n.s | n.s |
| HIF1A (TFFM0139.1) | 32.1% | 32.6% | 32.1% | n.s | n.s | n.s |
| HLF (TFFM0500.1) | 5.6% | 5.4% | 5.2% | n.s | n.s | n.s |
| HLF (TFFM0500.2) | 5.1% | 4.9% | 4.9% | n.s | n.s | n.s |
| HMBOX1 (TFFM0177.1) | 0.9% | 1.2% | 0.9% | n.s | n.s | n.s |
| HNF1A (TFFM0503.1) | 4.7% | 4.7% | 4.5% | n.s | n.s | n.s |
| HNF1B (TFFM0178.1) | 4.5% | 4.5% | 4.5% | n.s | n.s | n.s |
| HNF4A (TFFM0045.1) | 3.1% | 2.7% | 2.7% | n.s | n.s | n.s |
| HNF4A (TFFM0045.2) | 1.7% | 1.6% | 1.6% | n.s | n.s | n.s |
| HNF4A (TFFM0860.1) | 1.3% | 1.3% | 1.3% | n.s | n.s | n.s |
| HNF4G (TFFM0727.1) | 2.2% | 2.1% | 2.1% | n.s | n.s | n.s |
| HOXA9 (TFFM0179.1) | 1.6% | 1.6% | 1.5% | n.s | n.s | n.s |
| HOXA9 (TFFM0179.2) | 0.5% | 0.5% | 0.5% | n.s | n.s | n.s |
| HOXB4 (TFFM0505.1) | 8.0% | 7.6% | 7.9% | n.s | n.s | n.s |
| HOXB5 (TFFM0806.1) | 15.1% | 15.7% | 15.7% | n.s | n.s | n.s |
| HOXB8 (TFFM0861.1) | 0.4% | 0.4% | 0.4% | n.s | n.s | n.s |
| HOXC10 (TFFM0807.1) | 8.4% | 8.5% | 8.1% | n.s | n.s | n.s |
| IRF1 (TFFM0049.1) | 6.0% | 6.1% | 6.5% | n.s | n.s | n.s |
| IRF3 (TFFM0851.1) | 4.0% | 3.8% | 3.7% | n.s | n.s | n.s |
| JUN (TFFM0051.1) | 6.2% | 6.3% | 6.2% | n.s | n.s | n.s |
| JUN (TFFM0728.1) | 3.2% | 3.4% | 3.2% | n.s | n.s | n.s |
| JUNB (TFFM0052.1) | 4.7% | 4.8% | 4.7% | n.s | n.s | n.s |
| JUNB (TFFM0052.2) | 4.9% | 5.0% | 5.0% | n.s | n.s | n.s |
| JUND (TFFM0053.1) | 5.2% | 5.3% | 5.2% | n.s | n.s | n.s |
| JUND (TFFM0053.2) | 3.3% | 3.4% | 3.2% | n.s | n.s | n.s |
| KLF12 (TFFM0780.1) | 33.6% | 33.1% | 32.9% | n.s | n.s | n.s |
| KLF15 (TFFM0942.1) | 2.9% | 2.5% | 2.5% | n.s | n.s | n.s |
| KLF16 (TFFM0516.1) | 20.9% | 21.8% | 21.1% | n.s | n.s | n.s |
| KLF1 (TFFM0729.1) | 26.0% | 25.8% | 25.9% | n.s | n.s | n.s |
| KLF3 (TFFM0517.1) | 39.6% | 41.1% | 41.0% | n.s | n.s | n.s |
| KLF4 (TFFM0056.1) | 28.4% | 28.3% | 28.8% | n.s | n.s | n.s |

Continued on next page

Table S3 – continued from previous page

| TFBS | type 0 | UHP | DHP | 0 vs. UHP | 0 vs DHP | UHP vs. DHP |
|---|---|---|---|---|---|---|
| KLF9 (TFFM0140.1) | 21.6% | 21.8% | 20.8% | n.s | n.s | n.s |
| LEF1 (TFFM0184.1) | 6.1% | 5.9% | 5.8% | n.s | n.s | n.s |
| LHX2 (TFFM0185.1) | 1.7% | 1.4% | 1.6% | n.s | n.s | n.s |
| LHX3 (TFFM0717.1) | 2.0% | 1.9% | 2.1% | n.s | n.s | n.s |
| LHX6 (TFFM0758.1) | 9.2% | 9.0% | 9.2% | n.s | n.s | n.s |
| LMX1B (TFFM0772.1) | 5.1% | 5.3% | 5.0% | n.s | n.s | n.s |
| MAFF (TFFM0730.1) | 7.5% | 7.5% | 7.4% | n.s | n.s | n.s |
| MAFK (TFFM0058.1) | 16.4% | 16.1% | 16.3% | n.s | n.s | n.s |
| MAX (TFFM0059.1) | 11.1% | 11.2% | 10.8% | n.s | n.s | n.s |
| MEF2A (TFFM0060.1) | 9.2% | 9.3% | 9.2% | n.s | n.s | n.s |
| MEF2A (TFFM0060.2) | 9.3% | 9.2% | 9.5% | n.s | n.s | n.s |
| MEF2C (TFFM0061.1) | 15.0% | 15.7% | 15.6% | n.s | n.s | n.s |
| MEF2D (TFFM0525.1) | 10.4% | 10.5% | 10.6% | n.s | n.s | n.s |
| MEIS2 (TFFM0190.1) | 2.5% | 3.0% | 2.8% | n.s | n.s | n.s |
| MEIS2 (TFFM0895.1) | 6.4% | 6.1% | 6.1% | n.s | n.s | n.s |
| MGA (TFFM0526.1) | 11.6% | 11.6% | 11.2% | n.s | n.s | n.s |
| MYB (TFFM0064.1) | 13.2% | 13.3% | 13.5% | n.s | n.s | n.s |
| MYB (TFFM0064.2) | 8.5% | 8.6% | 8.3% | n.s | n.s | n.s |
| MYBL2 (TFFM0065.1) | 3.9% | 4.0% | 4.0% | n.s | n.s | n.s |
| MYC (TFFM0066.1) | 17.4% | 17.7% | 17.3% | n.s | n.s | n.s |
| MYC (TFFM0066.2) | 6.0% | 6.1% | 5.6% | n.s | n.s | n.s |
| MYOG (TFFM0069.1) | 8.1% | 7.8% | 8.0% | n.s | n.s | n.s |
| MYOG (TFFM0069.2) | 14.0% | 13.6% | 14.1% | n.s | n.s | n.s |
| NEUROG2 (TFFM0534.1) | 14.0% | 13.7% | 14.2% | n.s | n.s | n.s |
| NEUROG2 (TFFM0896.1) | 14.7% | 14.3% | 15.0% | n.s | n.s | n.s |
| NFE2 (TFFM0070.1) | 6.0% | 5.8% | 5.8% | n.s | n.s | n.s |
| NFE2L1 (TFFM0536.1) | 5.8% | 5.7% | 6.1% | n.s | n.s | n.s |
| NFIB (TFFM0681.1) | 2.8% | 2.4% | 2.3% | n.s | n.s | n.s |
| NFIL3 (TFFM0539.2) | 10.4% | 10.6% | 10.5% | n.s | n.s | n.s |
| NFIX (TFFM0864.1) | 2.8% | 2.5% | 2.5% | n.s | n.s | n.s |
| NFKB1 (TFFM0073.1) | 3.7% | 3.8% | 3.4% | n.s | n.s | n.s |
| NKX2-2 (TFFM0683.1) | 0.9% | 0.7% | 0.5% | n.s | n.s | n.s |
| NKX2-5 (TFFM0076.2) | 2.0% | 1.9% | 1.6% | n.s | n.s | n.s |
| NKX2-5 (TFFM0077.1) | 0.6% | 0.4% | 0.5% | n.s | n.s | n.s |
| NKX3-1 (TFFM0078.1) | 5.8% | 5.7% | 5.4% | n.s | n.s | n.s |
| NKX3-2 (TFFM0716.1) | 5.3% | 5.8% | 5.3% | n.s | n.s | n.s |
| NKX6-1 (TFFM0762.1) | 2.2% | 1.9% | 1.9% | n.s | n.s | n.s |
| NR1D1 (TFFM0865.1) | 0.4% | 0.4% | 0.4% | n.s | n.s | n.s |
| NR1D2 (TFFM0866.1) | 0.7% | 0.7% | 0.6% | n.s | n.s | n.s |
| NR1H2 (TFFM0993.1) | 11.4% | 11.7% | 11.8% | n.s | n.s | n.s |
| NR1H4 (TFFM0828.1) | 1.6% | 1.6% | 1.5% | n.s | n.s | n.s |
| NR2C2 (TFFM0867.1) | 2.7% | 2.7% | 2.2% | n.s | n.s | n.s |
| NR2F1 (TFFM0868.1) | 6.2% | 6.7% | 6.6% | n.s | n.s | n.s |
| NR2F6 (TFFM0195.1) | 7.3% | 7.0% | 7.3% | n.s | n.s | n.s |

Continued on next page

Table S3 – continued from previous page

| TFBS | type 0 | UHP | DHP | 0 vs. UHP | 0 vs DHP | UHP vs. DHP |
|---|---|---|---|---|---|---|
| NR2F6 (TFFM0870.1) | 21.3% | 21.5% | 21.1% | n.s | n.s | n.s |
| NR4A1 (TFFM0145.1) | 2.5% | 2.6% | 2.4% | n.s | n.s | n.s |
| NR4A1 (TFFM0145.2) | 1.8% | 2.1% | 1.7% | n.s | n.s | n.s |
| NR5A2 (TFFM0081.1) | 3.2% | 3.1% | 2.9% | n.s | n.s | n.s |
| ONECUT1 (TFFM0196.1) | 0.7% | 0.8% | 0.6% | n.s | n.s | n.s |
| ONECUT1 (TFFM0196.2) | 1.1% | 1.1% | 1.0% | n.s | n.s | n.s |
| ONECUT2 (TFFM0546.1) | 3.9% | 3.5% | 3.8% | n.s | n.s | n.s |
| ONECUT2 (TFFM0783.1) | 0.6% | 0.6% | 0.4% | n.s | n.s | n.s |
| OSR1 (TFFM0872.1) | 0.2% | 0.2% | 0.2% | n.s | n.s | n.s |
| OSR2 (TFFM0897.1) | 25.4% | 25.7% | 25.2% | n.s | n.s | n.s |
| OTX2 (TFFM0197.2) | 3.9% | 4.2% | 4.2% | n.s | n.s | n.s |
| OVOL1 (TFFM0873.1) | 1.5% | 1.8% | 1.8% | n.s | n.s | n.s |
| PAX3 (TFFM0787.1) | 0.3% | 0.3% | 0.3% | n.s | n.s | n.s |
| PAX3 (TFFM0874.1) | 4.6% | 4.8% | 5.0% | n.s | n.s | n.s |
| PAX5 (TFFM0084.1) | 5.6% | 5.4% | 5.5% | n.s | n.s | n.s |
| PAX7 (TFFM0550.1) | 1.3% | 1.3% | 1.6% | n.s | n.s | n.s |
| PAX7 (TFFM0764.1) | 2.7% | 2.7% | 3.1% | n.s | n.s | n.s |
| PBX1 (TFFM0551.1) | 5.5% | 5.3% | 5.4% | n.s | n.s | n.s |
| PBX2 (TFFM0146.2) | 4.9% | 5.2% | 5.4% | n.s | n.s | n.s |
| PBX3 (TFFM0147.1) | 5.7% | 5.6% | 5.4% | n.s | n.s | n.s |
| PDX1 (TFFM0198.2) | 3.6% | 3.1% | 3.3% | n.s | n.s | n.s |
| PHOX2A (TFFM0773.1) | 2.8% | 3.1% | 3.2% | n.s | n.s | n.s |
| PHOX2B (TFFM0554.1) | 0.7% | 0.7% | 0.6% | n.s | n.s | n.s |
| PHOX2B (TFFM0554.2) | 0.6% | 0.7% | 0.6% | n.s | n.s | n.s |
| PITX1 (TFFM0559.1) | 1.2% | 1.1% | 0.9% | n.s | n.s | n.s |
| POU2F3 (TFFM0562.1) | 6.6% | 6.8% | 6.6% | n.s | n.s | n.s |
| POU2F3 (TFFM0562.2) | 7.0% | 6.9% | 6.9% | n.s | n.s | n.s |
| POU3F1 (TFFM0789.1) | 0.8% | 0.8% | 0.8% | n.s | n.s | n.s |
| POU3F2 (TFFM0563.1) | 3.2% | 3.2% | 3.3% | n.s | n.s | n.s |
| PPARD (TFFM0875.1) | 3.5% | 4.0% | 3.6% | n.s | n.s | n.s |
| PRDM1 (TFFM0087.1) | 5.1% | 5.3% | 5.6% | n.s | n.s | n.s |
| PRDM4 (TFFM0898.1) | 4.1% | 4.3% | 4.2% | n.s | n.s | n.s |
| PROP1 (TFFM0774.1) | 4.9% | 5.1% | 5.1% | n.s | n.s | n.s |
| PROX1 (TFFM0199.1) | 0.1% | 0.0% | 0.1% | n.s | n.s | n.s |
| PTF1A (TFFM0887.1) | 9.5% | 9.1% | 9.2% | n.s | n.s | n.s |
| RARA (TFFM0571.1) | 2.6% | 2.4% | 2.6% | n.s | n.s | n.s |
| RARA (TFFM0777.1) | 0.2% | 0.1% | 0.1% | n.s | n.s | n.s |
| RARB (TFFM0802.1) | 0.4% | 0.3% | 0.3% | n.s | n.s | n.s |
| RARB (TFFM0876.1) | 0.3% | 0.2% | 0.2% | n.s | n.s | n.s |
| RELA (TFFM0200.1) | 4.5% | 4.1% | 4.3% | n.s | n.s | n.s |
| REST (TFFM0088.1) | 10.3% | 10.1% | 10.1% | n.s | n.s | n.s |
| RORA (TFFM0709.1) | 0.2% | 0.1% | 0.1% | n.s | n.s | n.s |
| RORB (TFFM0830.1) | 14.1% | 14.4% | 14.8% | n.s | n.s | n.s |
| RORC (TFFM0578.1) | 9.8% | 9.8% | 9.7% | n.s | n.s | n.s |

Continued on next page

| TFBS | type 0 | UHP | DHP | 0 vs. UHP | 0 vs DHP | UHP vs. DHP |
|---|---|---|---|---|---|---|
| RUNX1 (TFFM0091.1) | 8.3% | 7.9% | 8.3% | n.s | n.s | n.s |
| RUNX2 (TFFM0092.1) | 25.8% | 25.7% | 26.1% | n.s | n.s | n.s |
| RUNX3 (TFFM0093.2) | 4.4% | 4.3% | 3.9% | n.s | n.s | n.s |
| RXRA (TFFM0094.1) | 3.1% | 2.8% | 2.7% | n.s | n.s | n.s |
| RXRB (TFFM0579.1) | 0.7% | 0.7% | 0.8% | n.s | n.s | n.s |
| SCRT2 (TFFM0581.1) | 7.3% | 7.1% | 6.9% | n.s | n.s | n.s |
| SCRT2 (TFFM0581.2) | 6.9% | 6.9% | 6.5% | n.s | n.s | n.s |
| SIX2 (TFFM0150.1) | 5.2% | 5.1% | 5.5% | n.s | n.s | n.s |
| SMAD2 (TFFM0201.1) | 5.4% | 5.8% | 5.6% | n.s | n.s | n.s |
| SMAD3 (TFFM0202.1) | 0.2% | 0.1% | 0.2% | n.s | n.s | n.s |
| SOX11 (TFFM0803.1) | 5.4% | 5.5% | 5.0% | n.s | n.s | n.s |
| SOX13 (TFFM0586.1) | 5.8% | 6.1% | 5.9% | n.s | n.s | n.s |
| SOX2 (TFFM0095.2) | 7.6% | 7.6% | 7.3% | n.s | n.s | n.s |
| SOX3 (TFFM0096.1) | 9.8% | 9.6% | 9.8% | n.s | n.s | n.s |
| SOX6 (TFFM0588.1) | 3.0% | 2.6% | 2.6% | n.s | n.s | n.s |
| SP1 (TFFM0712.1) | 30.0% | 31.1% | 30.5% | n.s | n.s | n.s |
| SP2 (TFFM0735.1) | 27.5% | 28.1% | 27.3% | n.s | n.s | n.s |
| SP3 (TFFM0590.1) | 18.6% | 19.0% | 18.5% | n.s | n.s | n.s |
| SP4 (TFFM0765.1) | 26.6% | 27.5% | 26.7% | n.s | n.s | n.s |
| SREBF1 (TFFM0205.1) | 7.0% | 6.8% | 6.9% | n.s | n.s | n.s |
| SREBF1 (TFFM0206.1) | 6.5% | 6.3% | 6.3% | n.s | n.s | n.s |
| STAT1 (TFFM0101.1) | 6.5% | 6.6% | 6.2% | n.s | n.s | n.s |
| STAT4 (TFFM0103.1) | 13.6% | 13.5% | 13.4% | n.s | n.s | n.s |
| STAT5A (TFFM0594.1) | 7.7% | 7.3% | 7.3% | n.s | n.s | n.s |
| STAT5B (TFFM0595.1) | 9.3% | 9.2% | 9.3% | n.s | n.s | n.s |
| TBX21 (TFFM0208.1) | 3.9% | 3.7% | 3.5% | n.s | n.s | n.s |
| TBX21 (TFFM0767.1) | 0.3% | 0.2% | 0.2% | n.s | n.s | n.s |
| TBX3 (TFFM0598.1) | 3.9% | 4.1% | 3.7% | n.s | n.s | n.s |
| TBX3 (TFFM0879.1) | 0.5% | 0.5% | 0.5% | n.s | n.s | n.s |
| TCF21 (TFFM0881.1) | 32.4% | 31.7% | 32.9% | n.s | n.s | n.s |
| TCF7 (TFFM0209.1) | 1.9% | 2.0% | 1.7% | n.s | n.s | n.s |
| TCF7L2 (TFFM0109.1) | 2.5% | 2.4% | 2.3% | n.s | n.s | n.s |
| TEAD1 (TFFM0210.2) | 2.8% | 2.7% | 2.8% | n.s | n.s | n.s |
| TEAD3 (TFFM0603.1) | 1.2% | 1.2% | 1.3% | n.s | n.s | n.s |
| TEAD4 (TFFM0110.1) | 16.5% | 15.9% | 16.1% | n.s | n.s | n.s |
| TEAD4 (TFFM0110.2) | 1.8% | 1.9% | 2.1% | n.s | n.s | n.s |
| TFAP2A (TFFM0111.1) | 18.1% | 17.9% | 17.5% | n.s | n.s | n.s |
| TFAP2A (TFFM0113.1) | 17.3% | 17.4% | 17.5% | n.s | n.s | n.s |
| TFAP4 (TFFM0882.1) | 9.3% | 8.9% | 9.0% | n.s | n.s | n.s |
| TFCP2 (TFFM0929.1) | 3.5% | 3.7% | 3.7% | n.s | n.s | n.s |
| TFDP1 (TFFM0154.1) | 6.6% | 6.6% | 6.8% | n.s | n.s | n.s |
| TFE3 (TFFM0605.1) | 7.6% | 7.6% | 7.9% | n.s | n.s | n.s |
| TGIF1 (TFFM0606.1) | 17.2% | 17.8% | 17.9% | n.s | n.s | n.s |
| TGIF2 (TFFM0790.1) | 0.5% | 0.6% | 0.5% | n.s | n.s | n.s |

Continued on next page

**Table S3 – continued from previous page**

| TFBS | type 0 | UHP | DHP | 0 vs. UHP | 0 vs DHP | UHP vs. DHP |
|---|---|---|---|---|---|---|
| THAP1 (TFFM0213.1) | 2.8% | 2.7% | 2.8% | n.s | n.s | n.s |
| THAP1 (TFFM0744.1) | 1.8% | 2.3% | 2.1% | n.s | n.s | n.s |
| THRB (TFFM0884.1) | 6.0% | 6.1% | 6.2% | n.s | n.s | n.s |
| TP53 (TFFM0119.1) | 2.9% | 2.9% | 3.0% | n.s | n.s | n.s |
| TWIST1 (TFFM0155.1) | 12.5% | 12.0% | 12.5% | n.s | n.s | n.s |
| VDR (TFFM0769.1) | 0.2% | 0.3% | 0.1% | n.s | n.s | n.s |
| VEZF1 (TFFM0616.1) | 19.6% | 19.9% | 19.4% | n.s | n.s | n.s |
| XBP1 (TFFM0214.1) | 6.0% | 5.7% | 6.2% | n.s | n.s | n.s |
| ZBED2 (TFFM0979.1) | 4.6% | 5.0% | 4.8% | n.s | n.s | n.s |
| ZBTB12 (TFFM0693.1) | 3.5% | 3.5% | 3.7% | n.s | n.s | n.s |
| ZBTB14 (TFFM0694.1) | 0.8% | 0.8% | 0.7% | n.s | n.s | n.s |
| ZBTB33 (TFFM0125.1) | 3.7% | 3.6% | 3.7% | n.s | n.s | n.s |
| ZEB1 (TFFM0127.2) | 11.9% | 12.1% | 12.4% | n.s | n.s | n.s |
| ZFP57 (TFFM0627.1) | 26.8% | 28.0% | 27.6% | n.s | n.s | n.s |
| ZFX (TFFM0128.1) | 11.2% | 11.6% | 11.6% | n.s | n.s | n.s |
| ZIC5 (TFFM0886.1) | 23.6% | 23.9% | 23.2% | n.s | n.s | n.s |
| ZKSCAN3 (TFFM0981.1) | 0.9% | 0.9% | 0.8% | n.s | n.s | n.s |
| ZKSCAN5 (TFFM0697.1) | 20.9% | 20.7% | 21.0% | n.s | n.s | n.s |
| ZNF140 (TFFM0634.1) | 1.7% | 1.8% | 2.0% | n.s | n.s | n.s |
| ZNF143 (TFFM0129.1) | 5.1% | 5.6% | 5.3% | n.s | n.s | n.s |
| ZNF148 (TFFM0698.1) | 25.1% | 25.0% | 24.7% | n.s | n.s | n.s |
| ZNF16 (TFFM0699.1) | 0.4% | 0.2% | 0.3% | n.s | n.s | n.s |
| ZNF24 (TFFM0156.1) | 0.8% | 0.7% | 0.7% | n.s | n.s | n.s |
| ZNF263 (TFFM0130.1) | 11.3% | 11.2% | 11.6% | n.s | n.s | n.s |
| ZNF263 (TFFM0130.2) | 7.7% | 7.6% | 7.5% | n.s | n.s | n.s |
| ZNF281 (TFFM0889.1) | 16.0% | 16.1% | 16.1% | n.s | n.s | n.s |
| ZNF282 (TFFM0638.1) | 16.3% | 16.4% | 16.1% | n.s | n.s | n.s |
| ZNF382 (TFFM0640.1) | 1.8% | 1.7% | 1.7% | n.s | n.s | n.s |
| ZNF384 (TFFM0157.1) | 0.8% | 0.6% | 0.8% | n.s | n.s | n.s |
| ZNF416 (TFFM0982.1) | 10.7% | 10.5% | 10.2% | n.s | n.s | n.s |
| ZNF460 (TFFM0642.1) | 3.8% | 3.6% | 3.5% | n.s | n.s | n.s |
| ZNF528 (TFFM0643.1) | 1.4% | 1.5% | 1.5% | n.s | n.s | n.s |
| ZNF582 (TFFM0983.1) | 47.4% | 48.8% | 48.3% | n.s | n.s | n.s |
| ZNF675 (TFFM0911.1) | 3.5% | 3.0% | 3.2% | n.s | n.s | n.s |
| ZNF680 (TFFM0922.1) | 4.5% | 4.7% | 4.9% | n.s | n.s | n.s |
| ZNF682 (TFFM0645.1) | 4.6% | 4.4% | 4.5% | n.s | n.s | n.s |
| ZNF707 (TFFM0912.1) | 10.4% | 10.2% | 10.4% | n.s | n.s | n.s |
| ZNF768 (TFFM0924.1) | 3.5% | 3.6% | 3.6% | n.s | n.s | n.s |
| ZNF85 (TFFM0915.1) | 2.4% | 2.4% | 2.4% | n.s | n.s | n.s |
| ZNF8 (TFFM0913.1) | 4.9% | 4.7% | 4.6% | n.s | n.s | n.s |
| ZSCAN4 (TFFM0831.1) | 1.0% | 0.8% | 0.9% | n.s | n.s | n.s |

| **CPE/feature** | **type 0** | **UHP** | **DHP** | **0 vs. UHP** | **0 vs DHP** | **UHP vs. DHP** |
|---|---|---|---|---|---|---|
| AnyTags | 99.3% | 99.9% | 99.9% | n.s | n.s | n.s |
| BREd | 16.4% | 19.3% | 18.1% | 0.002699* | 0.06 | 0.18 |
| BREu | 15.1% | 13.8% | 14.6% | 0.09 | n.s | 0.26 |
| Bridge | 2.2% | 2.3% | 2.0% | n.s | n.s | n.s |
| Broad | 90.8% | 91.8% | 92.1% | n.s | n.s | n.s |
| CpG | 61.3% | 65.4% | 66.4% | 0.000963* | $4.20 \times 10^{-5}$ * | n.s |
| DCE | 2.2% | 2.6% | 2.4% | n.s | n.s | n.s |
| DCE3 | 14.3% | 14.3% | 13.8% | n.s | n.s | n.s |
| DPE | 12.6% | 12.4% | 13.0% | n.s | n.s | n.s |
| Inr | 39.7% | 40.4% | 38.8% | n.s | n.s | n.s |
| MTE | 0.9% | 0.8% | 0.9% | n.s | n.s | n.s |
| Sharp | 8.6% | 8.1% | 7.8% | n.s | 0.16 | n.s |
| TATA | 10.0% | 7.4% | 6.4% | 0.000472* | $p < 10^{-6}$ * | 0.15 |
| TCT | 12.4% | 11.8% | 11.3% | n.s | 0.10 | n.s |
| XCPE1 | 3.1% | 2.6% | 2.4% | n.s | 0.08 | n.s |
| XCPE2 | 2.2% | 2.7% | 3.0% | n.s | 0.036007 | n.s |

**Table S4:** Frequency of occurrence of core promoter elements (CPEs) and other promoter-related features defined as in (56). See main text for definitions of `type 0`, `UHP`, and `DHP`. The Bonferroni corrected threshold of $\alpha$ = 0.05 is $3.57 \times 10^{-3}$. *) significant at this threshold.

**Table S5:** Frequency of occurrence of RNA Binding Protein (RBP) Binding Sites. See main text for definitions of `type 0`, `UHP`, and `DHP`. The Bonferroni corrected threshold of $\alpha$ = 0.05 is $7.35 \times 10^{-4}$. *) significant at this threshold.

| RBP | type 0 | UHP | DHP | 0 vs. UHP | 0 vs DHP | UHP vs. DHP |
|---|---|---|---|---|---|---|
| ELAVL1 (1170_19561594) | 48.6% | 55.2% | 54.4% | $p < 10^{-6}$ * | $p < 10^{-6}$ * | n.s |
| RBM4 (1172_19561594) | 25.0% | 20.8% | 19.6% | $p < 10^{-6}$ * | $p < 10^{-6}$ * | 0.018992 |
| KHDRBS3 (1174_19561594) | 30.6% | 36.7% | 35.5% | $p < 10^{-6}$ * | $p < 10^{-6}$ * | n.s |
| Vts1 (1176_19561594) | 19.7% | 17.5% | 17.8% | $p < 10^{-6}$ * | $p < 10^{-6}$ * | n.s |
| YBX1 (1177_19561594) | 36.7% | 32.8% | 32.8% | $p < 10^{-6}$ * | $p < 10^{-6}$ * | n.s |
| QKI (1215_16041388) | 17.9% | 20.9% | 20.2% | $p < 10^{-6}$ * | $p < 10^{-6}$ * | n.s |
| ZRANB2 (1285_19304800) | 36.3% | 40.7% | 41.5% | $p < 10^{-6}$ * | $p < 10^{-6}$ * | n.s |
| HNRNPA1 (23_7510636) | 31.0% | 33.5% | 32.7% | $p < 10^{-6}$ * | $9.40 \times 10^{-5}$ * | n.s |
| SFRS1 (243_7543047) | 4.7% | 4.0% | 3.5% | $9.00 \times 10^{-5}$ * | $p < 10^{-6}$ * | 0.035681 |
| PABPC1 (24_7908267) | 20.2% | 22.9% | 22.7% | $p < 10^{-6}$ * | $p < 10^{-6}$ * | n.s |
| EIF4B (351_8846295) | 29.1% | 27.7% | 26.4% | 0.000237* | $p < 10^{-6}$ * | n.s |
| a2bp1 (36_12574126) | 25.2% | 22.9% | 23.1% | $p < 10^{-6}$ * | $p < 10^{-6}$ * | n.s |
| NONO (488_9001221) | 41.1% | 42.9% | 43.6% | n.s | $p < 10^{-6}$ * | n.s |
| FUS (637_11098054) | 28.3% | 25.4% | 24.7% | $p < 10^{-6}$ * | $p < 10^{-6}$ * | n.s |
| MBNL1 (669_20071745) | 32.0% | 29.5% | 28.9% | $p < 10^{-6}$ * | $p < 10^{-6}$ * | n.s |
| ELAVL2 (782_8497264) | 5.9% | 6.9% | 7.2% | $1.00 \times 10^{-5}$ * | $p < 10^{-6}$ * | n.s |
| ELAVL2 (783_7972035) | 10.9% | 14.1% | 14.2% | $p < 10^{-6}$ * | $p < 10^{-6}$ * | n.s |
| ELAVL2 (784_7972035) | 8.6% | 11.2% | 10.8% | $p < 10^{-6}$ * | $p < 10^{-6}$ * | n.s |
| RBMX (922_19282290) | 49.6% | 45.2% | 46.9% | $p < 10^{-6}$ * | $p < 10^{-6}$ * | n.s |
| PABPC1 (950_7908267) | 25.4% | 28.2% | 28.6% | $p < 10^{-6}$ * | $p < 10^{-6}$ * | n.s |
| ZFP36 (951_12324455) | 7.0% | 9.7% | 9.3% | $p < 10^{-6}$ * | $p < 10^{-6}$ * | n.s |
| SFRS1 (242_7543047) | 20.7% | 20.9% | 19.0% | n.s | $p < 10^{-6}$ * | 0.000267* |
| KHSRP (1186_17893325) | 9.3% | 8.7% | 8.1% | 0.015073 | $p < 10^{-6}$ * | 0.06 |
| Pum2 (329_11780640) | 8.3% | 9.4% | 8.9% | $p < 10^{-6}$ * | 0.009904 | n.s |
| sap-49 (145_9163526) | 18.9% | 17.4% | 17.9% | $1.80 \times 10^{-5}$ * | 0.004345 | n.s |
| Psi (915_11565747) | 19.5% | 21.0% | 20.5% | $2.50 \times 10^{-5}$ * | 0.002388 | n.s |
| ybx2-a (114_7499328) | 14.2% | 15.5% | 15.0% | $3.50 \times 10^{-5}$ * | 0.008450 | n.s |
| KHDRBS3 (1216_19457263) | 3.4% | 4.1% | 4.0% | 0.000121* | 0.000766 | n.s |
| ybx2-a (115_7499328) | 11.9% | 13.0% | 12.7% | 0.000294* | 0.006111 | n.s |
| ZFP36 (221_12324455) | 3.3% | 3.7% | 3.9% | n.s | 0.000311* | n.s |
| SNRPA (662_1717938) | 4.1% | 4.7% | 4.3% | 0.000955 | n.s | n.s |
| SFRS1 (952_7543047) | 9.7% | 9.0% | 9.2% | 0.002765 | n.s | n.s |
| QKI (149_16041388) | 2.3% | 2.8% | 2.2% | n.s | n.s | 0.003777 |
| RBMY1A1 (1052_17318228) | 15.3% | 14.4% | 15.5% | 0.005060 | n.s | 0.009171 |
| SNRPA (946_10094314) | 2.9% | 3.1% | 2.5% | n.s | n.s | 0.008204 |
| PTBP1 (1171_19561594) | 6.4% | 6.9% | 6.3% | n.s | n.s | 0.039223 |
| NCL (1004_8676391) | 0.0% | 0.0% | 0.0% | n.s | n.s | n.s |
| NCL (1026_10858445) | 5.1% | 5.3% | 4.9% | n.s | n.s | n.s |
| RBMY1A1 (1053_17318228) | 35.4% | 35.1% | 36.5% | n.s | n.s | n.s |
| SFRS13A (1169_19561594) | 61.5% | 61.1% | 61.1% | n.s | n.s | n.s |
| SFRS1 (1173_19561594) | 36.6% | 36.1% | 34.8% | n.s | n.s | n.s |

Continued on next page

Table S5 – continued from previous page

| RBP | type 0 | UHP | DHP | 0 vs. UHP | 0 vs DHP | UHP vs. DHP |
|---|---|---|---|---|---|---|
| SNRPA (1175_19561594) | 6.6% | 6.5% | 6.4% | n.s | n.s | n.s |
| KHSRP (1185_17893325) | 1.7% | 1.4% | 1.6% | n.s | n.s | n.s |
| ACO1 (1213_8021254) | 52.2% | 50.9% | 50.8% | n.s | n.s | n.s |
| ybx2-a (130_11376140) | 0.3% | 0.2% | 0.2% | n.s | n.s | n.s |
| NCL (131_11376140) | 1.1% | 1.2% | 1.2% | n.s | n.s | n.s |
| KHDRBS3 (147_19457263) | 2.9% | 3.0% | 3.0% | n.s | n.s | n.s |
| SFRS2 (244_7543047) | 11.3% | 11.8% | 11.4% | n.s | n.s | n.s |
| sus (254_1714588) | 8.9% | 9.0% | 8.9% | n.s | n.s | n.s |
| pum (323_16537387) | 2.0% | 2.4% | 2.3% | n.s | n.s | n.s |
| Pum2 (330_11780640) | 1.2% | 1.0% | 1.2% | n.s | n.s | n.s |
| EIF4B (350_8846295) | 30.4% | 30.0% | 29.7% | n.s | n.s | n.s |
| EIF4B (352_8846295) | 48.6% | 47.9% | 47.3% | n.s | n.s | n.s |
| IGF2BP1 (359_12507992) | 0.6% | 0.7% | 0.6% | n.s | n.s | n.s |
| Rna15 (376_9199325) | 0.3% | 0.2% | 0.3% | n.s | n.s | n.s |
| Rna15 (377_9199325) | 0.2% | 0.1% | 0.1% | n.s | n.s | n.s |
| A2BP1 (37_16537540) | 12.4% | 12.1% | 11.9% | n.s | n.s | n.s |
| SNRPA (661_1717938) | 3.4% | 3.8% | 3.5% | n.s | n.s | n.s |
| SNRPA (663_1717938) | 2.6% | 2.8% | 2.7% | n.s | n.s | n.s |
| NOVA2 (680_9789075) | 1.0% | 1.1% | 0.9% | n.s | n.s | n.s |
| NOVA2 (682_10811881) | 1.4% | 1.3% | 1.1% | n.s | n.s | n.s |
| SFRS7 (790_10094314) | 2.5% | 2.5% | 2.6% | n.s | n.s | n.s |
| SFRS2 (791_10094314) | 3.9% | 4.3% | 4.4% | n.s | n.s | n.s |
| SFRS9 (797_17548433) | 80.1% | 80.3% | 78.8% | n.s | n.s | n.s |
| B52 (802_9111335) | 0.0% | 0.0% | 0.0% | n.s | n.s | n.s |
| SNRPA (947_10094314) | 7.1% | 6.7% | 6.7% | n.s | n.s | n.s |
| SNRPA (948_10094314) | 1.8% | 1.8% | 1.7% | n.s | n.s | n.s |
| SNRPA (949_10094314) | 32.8% | 33.1% | 31.6% | n.s | n.s | n.s |
| SFRS2 (953_7543047) | 19.2% | 19.6% | 18.7% | n.s | n.s | n.s |
| SFRS7 (954_10094314) | 10.2% | 10.7% | 10.7% | n.s | n.s | n.s |
| YTHDC1 (969_20167602) | 83.0% | 81.8% | 81.6% | n.s | n.s | n.s |

| | | |
|---|---|---|
| ENCFF012VQA | ENCFF039TJY | ENCFF043FHJ |
| ENCFF061DKX | ENCFF064ASM | ENCFF065MQK |
| ENCFF082DKP | ENCFF105LDY | ENCFF107SJD |
| ENCFF129APS | ENCFF132MQB | ENCFF151LHR |
| ENCFF159PYD | ENCFF160KVW | ENCFF166EUJ |
| ENCFF175PDD | ENCFF195PBD | ENCFF211PWC |
| ENCFF216PEY | ENCFF218GFN | ENCFF226BZE |
| ENCFF227ELA | ENCFF229KAY | ENCFF235UTX |
| ENCFF245CAL | ENCFF254MJA | ENCFF258NNN |
| ENCFF292PBX | ENCFF322DAE | ENCFF341YAZ |
| ENCFF354VWZ | ENCFF355MNE | ENCFF371CYY |
| ENCFF376CIT | ENCFF389ZBU | ENCFF394NVG |
| ENCFF396ZQL | ENCFF410CPY | ENCFF411PMA |
| ENCFF448ZOJ | ENCFF451XZC | ENCFF456FTV |
| ENCFF469KDX | ENCFF471ZTS | ENCFF473YGW |
| ENCFF482BDW | ENCFF493EPF | ENCFF502RVO |
| ENCFF519DDH | ENCFF535TAL | ENCFF536KQX |
| ENCFF538YRS | ENCFF540RGI | ENCFF555UBC |
| ENCFF558UJR | ENCFF567ELA | ENCFF569MJS |
| ENCFF574UKD | ENCFF607OBG | ENCFF608MSL |
| ENCFF615EAT | ENCFF616GPO | ENCFF618BDP |
| ENCFF627GBZ | ENCFF634JRD | ENCFF653SLQ |
| ENCFF657NRR | ENCFF658XFZ | ENCFF672HZG |
| ENCFF673HLW | ENCFF673VSN | ENCFF680DFX |
| ENCFF681FKP | ENCFF681MRC | ENCFF683YQZ |
| ENCFF698HVQ | ENCFF712JXT | ENCFF730QNU |
| ENCFF745RUH | ENCFF750UAQ | ENCFF770ENO |
| ENCFF779EDF | ENCFF834UYS | ENCFF836KEF |
| ENCFF838LKZ | ENCFF842JME | ENCFF843PJA |
| ENCFF848IHI | ENCFF869HZM | ENCFF870KJE |
| ENCFF872EBX | ENCFF880CLF | ENCFF885ZWB |
| ENCFF886PSD | ENCFF890XQW | ENCFF898ZLY |
| ENCFF900JDD | ENCFF909CTD | ENCFF918UHB |
| ENCFF921FKB | ENCFF923UMY | ENCFF946CSJ |
| ENCFF952WTH | ENCFF967TFR | ENCFF993GPP |

**Table S6:** ChIP-Seq files used in computing the relative binding of RNA Polymerase subunit 2 to exons of types I,II and 0. Each of the entries refers to a BED file downloaded from ENCODE (43).
