## Supplemental file 2 for "Alternative splicing is coupled to gene expression in a subset of variably expressed genes"

| term | p.value | Coverage | adj.p | Term Name |
|---|---|---|---|---|
| GO:0004705 | 2.04E-54 | 0.61864407 | 2.89E-51 | JUN kinase activity |
| GO:0004707 | 2.22E-54 | 0.48648649 | 3.13E-51 | MAP kinase activity |
| GO:0004708 | 7.68E-50 | 0.59482759 | 1.08E-46 | MAP kinase kinase activity |
| GO:0009416 | 7.90E-44 | 0.6 | 1.11E-40 | response to light stimulus |
| GO:0003779 | 2.78E-35 | 0.19362187 | 3.91E-32 | actin binding |
| GO:0038095 | 2.66E-33 | 0.50980392 | 3.74E-30 | Fc-epsilon receptor signaling pathway |
| GO:0042765 | 6.37E-31 | 0.65 | 8.95E-28 | GPI-anchor transamidase complex |
| GO:0007010 | 7.16E-31 | 0.31557377 | 1.01E-27 | cytoskeleton organization |
| GO:0008092 | 1.30E-30 | 0.30952381 | 1.82E-27 | cytoskeletal protein binding |
| GO:0090398 | 1.02E-29 | 0.43801653 | 1.43E-26 | cellular senescence |
| GO:0016255 | 8.23E-29 | 0.65454545 | 1.15E-25 | attachment of GPI anchor to protein |
| GO:1990709 | 1.10E-28 | 1 | 1.54E-25 | presynaptic active zone organization |
| GO:0007254 | 4.24E-28 | 0.42857143 | 5.94E-25 | JNK cascade |
| GO:0010859 | 1.61E-27 | 1 | 2.26E-24 | calcium-dependent cysteine-type endopeptidase inhibitor activity |
| GO:0005739 | 2.41E-27 | 0.15780731 | 3.38E-24 | mitochondrion |
| GO:0051090 | 6.02E-26 | 0.48275862 | 8.41E-23 | regulation of DNA-binding transcription factor activity |
| GO:0006941 | 1.16E-25 | 0.79411765 | 1.62E-22 | striated muscle contraction |
| GO:0006099 | 7.22E-25 | 0.44791667 | 1.01E-21 | tricarboxylic acid cycle |
| GO:0006096 | 1.51E-24 | 0.42056075 | 2.11E-21 | glycolytic process |
| GO:0005200 | 1.09E-22 | 0.28703704 | 1.51E-19 | structural constituent of cytoskeleton |
| GO:0045296 | 6.22E-22 | 0.1626575 | 8.65E-19 | cadherin binding |
| GO:2000675 | 6.73E-22 | 0.75 | 9.36E-19 | negative regulation of type B pancreatic cell apoptotic process |
| GO:0048511 | 2.01E-21 | 0.34814815 | 2.79E-18 | rhythmic process |
| GO:0097340 | 4.38E-20 | 0.64102564 | 6.08E-17 | inhibition of cysteine-type endopeptidase activity |
| GO:0005759 | 2.67E-19 | 0.19396552 | 3.71E-16 | mitochondrial matrix |
| GO:0032559 | 8.69E-19 | 0.72413793 | 1.21E-15 | adenyl ribonucleotide binding |
| GO:0031032 | 2.55E-18 | 0.38297872 | 3.54E-15 | actomyosin structure organization |
| GO:0046514 | 2.82E-18 | 0.74074074 | 3.91E-15 | ceramide catabolic process |
| GO:0009317 | 3.43E-18 | 1 | 4.74E-15 | acetyl-CoA carboxylase complex |
| GO:0019901 | 1.14E-17 | 0.15589354 | 1.58E-14 | protein kinase binding |
| GO:0050810 | 1.45E-17 | 0.62857143 | 2.01E-14 | regulation of steroid biosynthetic process |
| GO:0030913 | 3.07E-17 | 0.73076923 | 4.24E-14 | paranodal junction assembly |
| GO:0005198 | 4.29E-17 | 0.21100917 | 5.93E-14 | structural molecule activity |
| GO:0062098 | 9.68E-17 | 0.7037037 | 1.34E-13 | regulation of programmed necrotic cell death |
| GO:0051287 | 9.91E-17 | 0.35714286 | 1.37E-13 | NAD binding |
| GO:0023051 | 1.86E-16 | 0.84210526 | 2.56E-13 | regulation of signaling |
| GO:0006937 | 3.26E-16 | 0.45 | 4.48E-13 | regulation of muscle contraction |
| GO:0042752 | 5.79E-16 | 0.29496403 | 7.96E-13 | regulation of circadian rhythm |
| GO:0004108 | 7.04E-16 | 0.93333333 | 9.68E-13 | citrate (Si)-synthase activity |
| GO:0004819 | 3.42E-15 | 0.76190476 | 4.69E-12 | glutamine-tRNA ligase activity |
| GO:0007274 | 6.04E-15 | 0.57142857 | 8.28E-12 | neuromuscular synaptic transmission |
| GO:0017064 | 7.35E-15 | 0.64285714 | 1.01E-11 | fatty acid amide hydrolase activity |
| GO:0006425 | 1.02E-14 | 0.78947368 | 1.40E-11 | glutaminyl-tRNA aminoacylation |
| GO:0003723 | 1.55E-14 | 0.10666181 | 2.12E-11 | RNA binding |
| GO:0005743 | 1.76E-14 | 0.17118998 | 2.41E-11 | mitochondrial inner membrane |
| GO:0045261 | 3.61E-14 | 0.69565217 | 4.93E-11 | proton-transporting ATP synthase complex, catalytic core F(1) |
| GO:0051402 | 6.77E-14 | 0.30357143 | 9.25E-11 | neuron apoptotic process |
| GO:0007339 | 9.80E-14 | 0.51282051 | 1.34E-10 | binding of sperm to zona pellucida |
| GO:0030955 | 1.18E-13 | 0.77777778 | 1.61E-10 | potassium ion binding |
| GO:0010807 | 1.18E-13 | 0.77777778 | 1.61E-10 | regulation of synaptic vesicle priming |
| GO:0004743 | 1.31E-13 | 0.92307692 | 1.78E-10 | pyruvate kinase activity |
| GO:0061049 | 3.19E-13 | 0.44 | 4.34E-10 | cell growth involved in cardiac muscle cell development |
| GO:0030182 | 4.33E-13 | 0.30188679 | 5.89E-10 | neuron differentiation |
| GO:0030866 | 6.24E-13 | 0.31578947 | 8.47E-10 | cortical actin cytoskeleton organization |
| GO:0044065 | 8.57E-13 | 0.85714286 | 1.16E-09 | regulation of respiratory system process |
| GO:0070296 | 8.57E-13 | 0.85714286 | 1.16E-09 | sarcoplasmic reticulum calcium ion transport |
| GO:0070972 | 8.57E-13 | 0.85714286 | 1.16E-09 | protein localization to endoplasmic reticulum |
| GO:0004449 | 9.90E-13 | 0.65217391 | 1.34E-09 | isocitrate dehydrogenase (NAD+) activity |
| GO:0005524 | 1.34E-12 | 0.10528407 | 1.81E-09 | ATP binding |
| GO:0046912 | 1.34E-12 | 0.76470588 | 1.82E-09 | acyltransferase activity, acyl groups converted into alkyl on transfer |
| GO:0035516 | 1.76E-12 | 0.91666667 | 2.38E-09 | oxidative DNA demethylase activity |
| GO:0071344 | 1.76E-12 | 0.91666667 | 2.38E-09 | diphosphate metabolic process |
| GO:0086066 | 1.76E-12 | 0.91666667 | 2.38E-09 | atrial cardiac muscle cell to AV node cell communication |
| GO:0030315 | 1.79E-12 | 0.45454545 | 2.42E-09 | T-tubule |
| GO:0106310 | 2.53E-12 | 0.16444444 | 3.40E-09 | protein serine kinase activity |
| GO:0032781 | 4.67E-12 | 0.37704918 | 6.29E-09 | positive regulation of ATP-dependent activity |
| GO:0000226 | 8.32E-12 | 0.20425532 | 1.12E-08 | microtubule cytoskeleton organization |
| GO:0099525 | 1.07E-11 | 0.84615385 | 1.45E-08 | presynaptic dense core vesicle exocytosis |
| GO:0044331 | 1.20E-11 | 0.5 | 1.62E-08 | cell-cell adhesion mediated by cadherin |
| GO:1904813 | 1.27E-11 | 0.22404372 | 1.70E-08 | ficolin-1-rich granule lumen |
| GO:0043202 | 1.49E-11 | 0.2556391 | 1.99E-08 | lysosomal lumen |
| GO:0043531 | 1.49E-11 | 0.359375 | 2.00E-08 | ADP binding |
| GO:0004672 | 1.60E-11 | 0.12465116 | 2.14E-08 | protein kinase activity |
| GO:0016082 | 1.68E-11 | 0.46153846 | 2.25E-08 | synaptic vesicle priming |
| GO:0033539 | 1.68E-11 | 0.46153846 | 2.25E-08 | fatty acid beta-oxidation using acyl-CoA dehydrogenase |
| GO:0015629 | 1.85E-11 | 0.16839378 | 2.47E-08 | actin cytoskeleton |
| GO:0071205 | 2.27E-11 | 0.60869565 | 3.03E-08 | protein localization to juxtaparanode region of axon |
| GO:0002175 | 2.36E-11 | 0.90909091 | 3.16E-08 | protein localization to paranode region of axon |
| GO:0014902 | 2.60E-11 | 0.51612903 | 3.47E-08 | myotube differentiation |
| GO:0099536 | 2.60E-11 | 0.51612903 | 3.47E-08 | synaptic signaling |
| GO:0030018 | 2.74E-11 | 0.24 | 3.65E-08 | Z disc |
| GO:0031013 | 3.35E-11 | 1 | 4.46E-08 | troponin I binding |
| GO:0005915 | 4.87E-11 | 0.5 | 6.48E-08 | zonula adherens |
| GO:0006754 | 5.09E-11 | 0.58333333 | 6.77E-08 | ATP biosynthetic process |
| GO:0046933 | 5.43E-11 | 0.53571429 | 7.22E-08 | proton-transporting ATP synthase activity, rotational mechanism |
| GO:0007043 | 6.24E-11 | 0.33823529 | 8.28E-08 | cell-cell junction assembly |
| GO:0006101 | 8.84E-11 | 0.61904762 | 1.17E-07 | citrate metabolic process |
| GO:0046513 | 1.40E-10 | 0.3559322 | 1.86E-07 | ceramide biosynthetic process |
| GO:0060048 | 1.40E-10 | 0.3559322 | 1.86E-07 | cardiac muscle contraction |
| GO:0005856 | 1.75E-10 | 0.14054927 | 2.32E-07 | cytoskeleton |
| GO:0006102 | 2.02E-10 | 0.59090909 | 2.68E-07 | isocitrate metabolic process |
| GO:0005912 | 2.72E-10 | 0.19815668 | 3.59E-07 | adherens junction |
| GO:0106022 | 3.14E-10 | 0.9 | 4.15E-07 | positive regulation of vesicle docking |
| GO:0071900 | 3.43E-10 | 0.63157895 | 4.52E-07 | regulation of protein serine/threonine kinase activity |
| GO:0098696 | 3.43E-10 | 0.63157895 | 4.52E-07 | regulation of neurotransmitter receptor localization to postsynaptic specialization membrane |
| GO:2001259 | 3.43E-10 | 0.63157895 | 4.52E-07 | positive regulation of cation channel activity |
| GO:0007015 | 3.49E-10 | 0.18215613 | 4.60E-07 | actin filament organization |
| GO:0030507 | 4.08E-10 | 0.3 | 5.37E-07 | spectrin binding |
| GO:0008307 | 4.12E-10 | 0.25 | 5.42E-07 | structural constituent of muscle |
| GO:0002639 | 4.28E-10 | 0.51851852 | 5.62E-07 | positive regulation of immunoglobulin production |
| GO:0051015 | 4.39E-10 | 0.14677104 | 5.75E-07 | actin filament binding |
| GO:0004347 | 4.89E-10 | 1 | 6.41E-07 | glucose-6-phosphate isomerase activity |
| GO:0032972 | 4.89E-10 | 1 | 6.41E-07 | regulation of muscle filament sliding speed |
| GO:0090646 | 4.89E-10 | 1 | 6.41E-07 | mitochondrial tRNA processing |
| GO:0032229 | 5.41E-10 | 0.76923077 | 7.07E-07 | negative regulation of synaptic transmission, GABAergic |
| GO:0106006 | 8.91E-10 | 0.54166667 | 1.16E-06 | cytoskeletal protein-membrane anchor activity |
| GO:0030049 | 8.91E-10 | 0.54166667 | 1.16E-06 | muscle filament sliding |
| GO:0042645 | 1.04E-09 | 0.33898305 | 1.36E-06 | mitochondrial nucleoid |
| GO:0006468 | 1.04E-09 | 0.11606392 | 1.36E-06 | protein phosphorylation |
| GO:0070061 | 1.08E-09 | 0.45454545 | 1.41E-06 | fructose binding |
| GO:0006446 | 1.50E-09 | 0.26530612 | 1.95E-06 | regulation of translational initiation |
| GO:0006120 | 1.51E-09 | 0.31818182 | 1.96E-06 | mitochondrial electron transport, NADH to ubiquinone |
| GO:0016747 | 1.60E-09 | 0.36734694 | 2.08E-06 | acyltransferase activity, transferring groups other than amino-acyl groups |
| GO:0043217 | 1.62E-09 | 0.81818182 | 2.11E-06 | myelin maintenance |
| GO:0031430 | 1.74E-09 | 0.52 | 2.26E-06 | M band |
| GO:0042776 | 1.74E-09 | 0.52 | 2.26E-06 | proton motive force-driven mitochondrial ATP synthesis |
| GO:0030176 | 2.50E-09 | 0.1959799 | 3.24E-06 | integral component of endoplasmic reticulum membrane |
| GO:0016342 | 2.55E-09 | 0.46666667 | 3.30E-06 | catenin complex |
| GO:0001725 | 2.67E-09 | 0.22222222 | 3.46E-06 | stress fiber |
| GO:0008093 | 2.78E-09 | 0.30882353 | 3.60E-06 | cytoskeletal anchor activity |
| GO:0061157 | 3.00E-09 | 0.4 | 3.88E-06 | mRNA destabilization |
| GO:0034604 | 3.17E-09 | 0.61111111 | 4.10E-06 | pyruvate dehydrogenase (NAD+) activity |
| GO:0016651 | 3.26E-09 | 0.5 | 4.20E-06 | oxidoreductase activity, acting on NAD(P)H |
| GO:0004332 | 3.62E-09 | 0.54545455 | 4.66E-06 | fructose-bisphosphate aldolase activity |
| GO:0005764 | 3.68E-09 | 0.15789474 | 4.74E-06 | lysosome |
| GO:0008360 | 3.73E-09 | 0.19047619 | 4.79E-06 | regulation of cell shape |
| GO:0045104 | 3.85E-09 | 0.31746032 | 4.96E-06 | intermediate filament cytoskeleton organization |
| GO:0045259 | 4.13E-09 | 0.88888889 | 5.31E-06 | proton-transporting ATP synthase complex |
| GO:0036371 | 4.13E-09 | 0.88888889 | 5.31E-06 | protein localization to T-tubule |
| GO:0090258 | 4.13E-09 | 0.88888889 | 5.31E-06 | negative regulation of mitochondrial fission |
| GO:0090335 | 4.13E-09 | 0.88888889 | 5.31E-06 | regulation of brown fat cell differentiation |
| GO:0010881 | 4.77E-09 | 0.41666667 | 6.11E-06 | regulation of cardiac muscle contraction by regulation of the release of sequestered calcium ion |
| GO:0048312 | 4.99E-09 | 0.66666667 | 6.39E-06 | intracellular distribution of mitochondria |
| GO:0015631 | 5.15E-09 | 0.22142857 | 6.58E-06 | tubulin binding |
| GO:0003988 | 5.89E-09 | 0.48148148 | 7.52E-06 | acetyl-CoA C-acyltransferase activity |
| GO:1904753 | 5.89E-09 | 0.48148148 | 7.52E-06 | negative regulation of vascular associated smooth muscle cell migration |
| GO:0005515 | 6.64E-09 | 0.09007982 | 8.47E-06 | protein binding |
| GO:0006086 | 7.07E-09 | 0.57894737 | 9.01E-06 | acetyl-CoA biosynthetic process from pyruvate |
| GO:0003989 | 7.09E-09 | 0.52173913 | 9.03E-06 | acetyl-CoA carboxylase activity |
| GO:0012501 | 7.09E-09 | 0.52173913 | 9.03E-06 | programmed cell death |
| GO:0030172 | 7.14E-09 | 1 | 9.08E-06 | troponin C binding |
| GO:0031071 | 7.14E-09 | 1 | 9.08E-06 | cysteine desulfurase activity |
| GO:0047860 | 7.14E-09 | 1 | 9.08E-06 | diiodophenylpyruvate reductase activity |
| GO:0042245 | 7.14E-09 | 1 | 9.08E-06 | RNA repair |
| GO:0098907 | 7.14E-09 | 1 | 9.08E-06 | regulation of SA node cell action potential |
| GO:1990938 | 7.14E-09 | 1 | 9.08E-06 | peptidyl-aspartic acid autophosphorylation |
| GO:0015031 | 9.30E-09 | 0.14255765 | 1.18E-05 | protein transport |
| GO:0055114 | 9.78E-09 | 0.13355593 | 1.24E-05 | oxidation-reduction process |
| GO:0030388 | 1.16E-08 | 0.39473684 | 1.47E-05 | fructose 1,6-bisphosphate metabolic process |
| GO:0003680 | 1.18E-08 | 0.42424242 | 1.49E-05 | minor groove of adenine-thymine-rich DNA binding |
| GO:0035552 | 1.25E-08 | 0.625 | 1.58E-05 | oxidative single-stranded DNA demethylation |
| GO:0008137 | 1.34E-08 | 0.32727273 | 1.69E-05 | NADH dehydrogenase (ubiquinone) activity |
| GO:0098685 | 1.44E-08 | 0.25531915 | 1.81E-05 | Schaffer collateral - CA1 synapse |
| GO:0003211 | 1.86E-08 | 0.69230769 | 2.34E-05 | cardiac ventricle formation |
| GO:0010882 | 1.86E-08 | 0.69230769 | 2.34E-05 | regulation of cardiac muscle contraction by calcium ion signaling |
| GO:0086070 | 1.94E-08 | 0.8 | 2.44E-05 | SA node cell to atrial cardiac muscle cell communication |

| | | | | |
|---|---|---|---|---|
| GO:0050750 | 2.52E-08 | 0.31578947 | 3.17E-05 | low-density lipoprotein particle receptor binding |
| GO:0043039 | 2.65E-08 | 0.375 | 3.32E-05 | tRNA aminoacylation |
| GO:0061732 | 2.84E-08 | 0.58823529 | 3.57E-05 | mitochondrial acetyl-CoA biosynthetic process from pyruvate |
| GO:0007268 | 2.88E-08 | 0.20666667 | 3.61E-05 | chemical synaptic transmission |
| GO:1900015 | 2.90E-08 | 0.52380952 | 3.63E-05 | regulation of cytokine production involved in inflammatory response |
| GO:0005829 | 4.69E-08 | 0.09137056 | 5.87E-05 | cytosol |
| GO:0046034 | 4.73E-08 | 0.32075472 | 5.91E-05 | ATP metabolic process |
| GO:0004658 | 4.88E-08 | 0.64285714 | 6.09E-05 | propionyl-CoA carboxylase activity |
| GO:0008308 | 4.88E-08 | 0.64285714 | 6.09E-05 | voltage-gated anion channel activity |
| GO:0003006 | 4.88E-08 | 0.64285714 | 6.09E-05 | developmental process involved in reproduction |
| GO:0019626 | 4.88E-08 | 0.64285714 | 6.09E-05 | short-chain fatty acid catabolic process |
| GO:0005134 | 5.37E-08 | 0.875 | 6.68E-05 | interleukin-2 receptor binding |
| GO:0016615 | 5.37E-08 | 0.875 | 6.68E-05 | malate dehydrogenase activity |
| GO:0045145 | 5.37E-08 | 0.875 | 6.68E-05 | single-stranded DNA 5'-3' exodeoxyribonuclease activity |
| GO:1901019 | 5.37E-08 | 0.875 | 6.68E-05 | regulation of calcium ion transmembrane transporter activity |
| GO:1904428 | 5.37E-08 | 0.875 | 6.68E-05 | negative regulation of tubulin deacetylation |
| GO:1904692 | 5.37E-08 | 0.875 | 6.68E-05 | positive regulation of type B pancreatic cell proliferation |
| GO:0000159 | 6.31E-08 | 0.2739726 | 7.81E-05 | protein phosphatase type 2A complex |
| GO:0003065 | 6.68E-08 | 0.72727273 | 8.27E-05 | positive regulation of heart rate by epinephrine |
| GO:0016078 | 6.68E-08 | 0.72727273 | 8.27E-05 | tRNA catabolic process |
| GO:1902115 | 6.68E-08 | 0.72727273 | 8.27E-05 | regulation of organelle assembly |
| GO:0006936 | 7.22E-08 | 0.19512195 | 8.92E-05 | muscle contraction |
| GO:0045773 | 8.09E-08 | 0.27027027 | 9.98E-05 | positive regulation of axon extension |
| GO:0006635 | 8.81E-08 | 0.22727273 | 0.000108675 | fatty acid beta-oxidation |
| GO:0034551 | 9.77E-08 | 0.47826087 | 0.000120328 | mitochondrial respiratory chain complex III assembly |
| GO:1902807 | 9.77E-08 | 0.47826087 | 0.000120328 | negative regulation of cell cycle G1/S phase transition |
| GO:2001135 | 9.77E-08 | 0.47826087 | 0.000120328 | regulation of endocytic recycling |
| GO:0005865 | 1.04E-07 | 1 | 0.000128027 | striated muscle thin filament |
| GO:1990584 | 1.04E-07 | 1 | 0.000128027 | cardiac Troponin complex |
| GO:0005460 | 1.04E-07 | 1 | 0.000128027 | UDP-glucose transmembrane transporter activity |
| GO:0032549 | 1.04E-07 | 1 | 0.000128027 | ribonucleoside binding |
| GO:0018283 | 1.04E-07 | 1 | 0.000128027 | iron incorporation into metallo-sulfur cluster |
| GO:0032272 | 1.04E-07 | 1 | 0.000128027 | negative regulation of protein polymerization |
| GO:0034111 | 1.04E-07 | 1 | 0.000128027 | negative regulation of homotypic cell-cell adhesion |
| GO:0072684 | 1.04E-07 | 1 | 0.000128027 | mitochondrial tRNA 3'-trailer cleavage, endonucleolytic |
| GO:0050633 | 1.14E-07 | 0.6 | 0.000139781 | acetyl-CoA C-myristoyltransferase activity |
| GO:0070476 | 1.14E-07 | 0.6 | 0.000139781 | rRNA (guanine-N7)-methylation |
| GO:2000553 | 1.14E-07 | 0.6 | 0.000139781 | positive regulation of T-helper 2 cell cytokine production |
| GO:0086005 | 1.15E-07 | 0.42857143 | 0.000140475 | ventricular cardiac muscle cell action potential |
| GO:0031333 | 1.18E-07 | 0.30357143 | 0.000143246 | negative regulation of protein-containing complex assembly |
| GO:0043560 | 1.19E-07 | 0.52631579 | 0.000144592 | insulin receptor substrate binding |
| GO:0060292 | 1.19E-07 | 0.52631579 | 0.000144592 | long-term synaptic depression |
| GO:0086004 | 1.19E-07 | 0.52631579 | 0.000144592 | regulation of cardiac muscle cell contraction |
| GO:0030426 | 1.36E-07 | 0.19375 | 0.000164386 | growth cone |
| GO:0005861 | 1.47E-07 | 0.35897436 | 0.000177562 | troponin complex |
| GO:0072659 | 1.49E-07 | 0.1745283 | 0.000180284 | protein localization to plasma membrane |
| GO:0030674 | 1.49E-07 | 0.18934911 | 0.000180649 | protein-macromolecule adaptor activity |
| GO:0004176 | 1.69E-07 | 0.45833333 | 0.000204405 | ATP-dependent peptidase activity |
| GO:0045214 | 1.71E-07 | 0.22641509 | 0.000206385 | sarcomere organization |
| GO:0097512 | 1.88E-07 | 0.66666667 | 0.000227287 | cardiac myofibril |
| GO:0009134 | 1.88E-07 | 0.66666667 | 0.000227287 | nucleoside diphosphate catabolic process |
| GO:0055012 | 1.88E-07 | 0.66666667 | 0.000227287 | ventricular cardiac muscle cell differentiation |
| GO:0031902 | 2.09E-07 | 0.19018405 | 0.000252038 | late endosome membrane |
| GO:0055010 | 2.11E-07 | 0.35 | 0.000253983 | ventricular cardiac muscle tissue morphogenesis |
| GO:0045176 | 2.23E-07 | 0.5 | 0.000268161 | apical protein localization |
| GO:0060973 | 2.23E-07 | 0.5 | 0.000268161 | cell migration involved in heart development |
| GO:0030060 | 2.27E-07 | 0.77777778 | 0.000272509 | L-malate dehydrogenase activity |
| GO:0032034 | 2.27E-07 | 0.77777778 | 0.000272509 | myosin II head/neck binding |
| GO:0007412 | 2.27E-07 | 0.77777778 | 0.000272509 | axon target recognition |
| GO:0005967 | 2.46E-07 | 0.5625 | 0.000293999 | mitochondrial pyruvate dehydrogenase complex |
| GO:1990829 | 2.46E-07 | 0.5625 | 0.000293999 | C-rich single-stranded DNA binding |
| GO:0032873 | 2.46E-07 | 0.5625 | 0.000293999 | negative regulation of stress-activated MAPK cascade |
| GO:0044805 | 2.46E-07 | 0.5625 | 0.000293999 | late nucleophagy |
| GO:0045761 | 2.46E-07 | 0.5625 | 0.000293999 | regulation of adenylate cyclase activity |
| GO:0051170 | 2.46E-07 | 0.5625 | 0.000293999 | import into nucleus |
| GO:0004674 | 2.51E-07 | 0.13319672 | 0.000299456 | protein serine/threonine kinase activity |
| GO:0008206 | 2.83E-07 | 0.44 | 0.000336881 | bile acid metabolic process |
| GO:0005761 | 3.01E-07 | 0.30188679 | 0.000357559 | mitochondrial ribosome |
| GO:0030216 | 3.77E-07 | 0.36111111 | 0.000448206 | keratinocyte differentiation |
| GO:0016874 | 4.00E-07 | 0.2962963 | 0.00047495 | ligase activity |
| GO:0015986 | 4.22E-07 | 0.33333333 | 0.000500469 | proton motive force-driven ATP synthesis |
| GO:0043274 | 4.26E-07 | 0.3125 | 0.000504582 | phospholipase binding |
| GO:0044224 | 4.41E-07 | 0.38709677 | 0.000521628 | juxtaparanode region of axon |
| GO:0033270 | 4.60E-07 | 0.61538462 | 0.000544055 | paranode region of axon |
| GO:0051560 | 4.60E-07 | 0.42307692 | 0.000544055 | mitochondrial calcium ion homeostasis |
| GO:1902966 | 4.60E-07 | 0.61538462 | 0.000544055 | positive regulation of protein localization to early endosome |
| GO:2000463 | 4.60E-07 | 0.42307692 | 0.000544055 | positive regulation of excitatory postsynaptic potential |
| GO:1900063 | 4.90E-07 | 0.52941176 | 0.000577619 | regulation of peroxisome organization |
| GO:1990504 | 6.86E-07 | 0.85714286 | 0.000808551 | dense core granule exocytosis |
| GO:0035196 | 6.87E-07 | 0.45454545 | 0.000808551 | miRNA processing |
| GO:0042698 | 6.86E-07 | 0.85714286 | 0.000808551 | ovulation cycle |
| GO:0070350 | 6.86E-07 | 0.85714286 | 0.000808551 | regulation of white fat cell proliferation |
| GO:0070407 | 6.86E-07 | 0.85714286 | 0.000808551 | oxidation-dependent protein catabolic process |
| GO:1901021 | 6.86E-07 | 0.85714286 | 0.000808551 | positive regulation of calcium ion transmembrane transporter activity |
| GO:1903296 | 6.86E-07 | 0.85714286 | 0.000808551 | positive regulation of glutamate secretion, neurotransmission |
| GO:0005962 | 7.12E-07 | 0.7 | 0.000833391 | mitochondrial isocitrate dehydrogenase complex (NAD+) |
| GO:0016715 | 7.12E-07 | 0.7 | 0.000833391 | oxidoreductase activity, acting on paired donors, with incorporation or reduction of molecular oxygen, reduced ascorbate as one donor, and incorporation of one atom of oxygen |
| GO:0055069 | 7.12E-07 | 0.7 | 0.000833391 | zinc ion homeostasis |
| GO:0060327 | 7.12E-07 | 0.7 | 0.000833391 | cytoplasmic actin-based contraction involved in cell motility |
| GO:0034452 | 7.28E-07 | 0.40740741 | 0.000849723 | dynactin binding |
| GO:0060307 | 7.28E-07 | 0.40740741 | 0.000849723 | regulation of ventricular cardiac muscle cell membrane repolarization |
| GO:0016020 | 8.55E-07 | 0.09134615 | 0.000995948 | membrane |
| GO:0051893 | 9.77E-07 | 0.265625 | 0.001137385 | regulation of focal adhesion assembly |
| GO:0008802 | 1.01E-06 | 0.57142857 | 0.001172347 | betaine-aldehyde dehydrogenase activity |
| GO:0032754 | 1.01E-06 | 0.57142857 | 0.001172347 | positive regulation of interleukin-5 production |
| GO:0035542 | 1.01E-06 | 0.57142857 | 0.001172347 | regulation of SNARE complex assembly |
| GO:0080111 | 1.01E-06 | 0.57142857 | 0.001172347 | DNA demethylation |
| GO:0086014 | 1.01E-06 | 0.57142857 | 0.001172347 | atrial cardiac muscle cell action potential |
| GO:0006744 | 1.09E-06 | 0.33333333 | 0.001263595 | ubiquinone biosynthetic process |
| GO:0072542 | 1.12E-06 | 0.39285714 | 0.001301208 | protein phosphatase activator activity |
| GO:0046512 | 1.12E-06 | 0.39285714 | 0.001301208 | sphingosine biosynthetic process |
| GO:0060038 | 1.12E-06 | 0.39285714 | 0.001301208 | cardiac muscle cell proliferation |
| GO:0030017 | 1.47E-06 | 0.30434783 | 0.001702042 | sarcomere |
| GO:0005459 | 1.52E-06 | 1 | 0.001752411 | UDP-galactose transmembrane transporter activity |
| GO:0035605 | 1.52E-06 | 1 | 0.001752411 | peptidyl-cysteine S-nitrosylase activity |
| GO:0097109 | 1.52E-06 | 1 | 0.001752411 | neuroligin family protein binding |
| GO:0031536 | 1.52E-06 | 1 | 0.001752411 | positive regulation of exit from mitosis |
| GO:0072386 | 1.52E-06 | 1 | 0.001752411 | plus-end-directed organelle transport along microtubule |
| GO:1900148 | 1.52E-06 | 1 | 0.001752411 | negative regulation of Schwann cell migration |
| GO:1902047 | 1.52E-06 | 1 | 0.001752411 | polyamine transmembrane transport |
| GO:1903772 | 1.52E-06 | 1 | 0.001752411 | regulation of viral budding via host ESCRT complex |
| GO:1904719 | 1.52E-06 | 1 | 0.001752411 | positive regulation of AMPA glutamate receptor clustering |
| GO:0034614 | 1.55E-06 | 0.24657534 | 0.001773413 | cellular response to reactive oxygen species |
| GO:0016616 | 1.60E-06 | 0.20168067 | 0.001831646 | oxidoreductase activity, acting on the CH-OH group of donors, NAD or NADP as acceptor |
| GO:0052548 | 1.64E-06 | 0.47368421 | 0.001873573 | regulation of endopeptidase activity |
| GO:2000727 | 1.64E-06 | 0.47368421 | 0.001873573 | positive regulation of cardiac muscle cell differentiation |
| GO:0005744 | 1.70E-06 | 0.37931034 | 0.00193712 | TIM23 mitochondrial import inner membrane translocase complex |
| GO:0099072 | 1.70E-06 | 0.37931034 | 0.00193712 | regulation of postsynaptic membrane neurotransmitter receptor levels |
| GO:0005791 | 1.75E-06 | 0.28301887 | 0.001996506 | rough endoplasmic reticulum |
| GO:0032456 | 1.81E-06 | 0.2345679 | 0.002059421 | endocytic recycling |
| GO:0009083 | 1.83E-06 | 0.41666667 | 0.002084352 | branched-chain amino acid catabolic process |
| GO:0010940 | 1.83E-06 | 0.41666667 | 0.002084352 | positive regulation of necrotic cell death |
| GO:0003185 | 1.84E-06 | 0.63636364 | 0.002086794 | sinoatrial valve morphogenesis |
| GO:0071374 | 1.84E-06 | 0.63636364 | 0.002086794 | cellular response to parathyroid hormone stimulus |
| GO:0034599 | 1.89E-06 | 0.20535714 | 0.002140414 | cellular response to oxidative stress |
| GO:0098609 | 1.99E-06 | 0.14569536 | 0.002245502 | cell-cell adhesion |
| GO:0031672 | 2.03E-06 | 0.53333333 | 0.002293045 | A band |
| GO:0016435 | 2.03E-06 | 0.53333333 | 0.002293045 | rRNA (guanine) methyltransferase activity |
| GO:0030518 | 2.03E-06 | 0.53333333 | 0.002293045 | intracellular steroid hormone receptor signaling pathway |
| GO:0070989 | 2.03E-06 | 0.53333333 | 0.002293045 | oxidative demethylation |
| GO:0008016 | 2.43E-06 | 0.26229508 | 0.002734143 | regulation of heart contraction |
| GO:0004598 | 2.58E-06 | 0.75 | 0.002907526 | peptidylamidoglycolate lyase activity |
| GO:0005483 | 2.58E-06 | 0.75 | 0.002907526 | soluble NSF attachment protein activity |
| GO:0008592 | 2.58E-06 | 0.75 | 0.002907526 | regulation of Toll signaling pathway |
| GO:0044571 | 2.58E-06 | 0.75 | 0.002907526 | [2Fe-2S] cluster assembly |
| GO:0009098 | 2.58E-06 | 0.75 | 0.002907526 | leucine biosynthetic process |
| GO:0061762 | 2.58E-06 | 0.75 | 0.002907526 | CAMKK-AMPK signaling cascade |
| GO:1902988 | 2.58E-06 | 0.75 | 0.002907526 | neurofibrillary tangle assembly |
| GO:0016491 | 2.78E-06 | 0.13013699 | 0.003103414 | oxidoreductase activity |
| GO:0051258 | 2.80E-06 | 0.45 | 0.003128035 | protein polymerization |
| GO:0047496 | 2.82E-06 | 0.30952381 | 0.003148881 | vesicle transport along microtubule |
| GO:0035050 | 2.83E-06 | 0.33333333 | 0.003154052 | embryonic heart tube development |
| GO:0045202 | 2.99E-06 | 0.14634146 | 0.003329435 | synapse |
| GO:0007165 | 3.13E-06 | 0.10049192 | 0.003488311 | signal transduction |
| GO:0017101 | 3.66E-06 | 0.35483871 | 0.004067494 | aminoacyl-tRNA synthetase multienzyme complex |
| GO:0007269 | 3.66E-06 | 0.35483871 | 0.004067494 | neurotransmitter secretion |
| GO:0001836 | 3.75E-06 | 0.26785714 | 0.004161455 | release of cytochrome c from mitochondria |
| GO:2000643 | 3.81E-06 | 0.5 | 0.004228476 | positive regulation of early endosome to late endosome transport |
| GO:0000814 | 4.15E-06 | 0.58333333 | 0.004601374 | ESCRT II complex |
| GO:2000234 | 4.15E-06 | 0.58333333 | 0.004601374 | positive regulation of rRNA processing |
| GO:0098978 | 4.27E-06 | 0.1443299 | 0.004718612 | glutamatergic synapse |
| GO:0008179 | 4.38E-06 | 0.38461538 | 0.004834408 | adenylate cyclase binding |
| GO:0055117 | 5.23E-06 | 0.34375 | 0.005771391 | regulation of cardiac muscle contraction |
| GO:0048854 | 5.74E-06 | 0.2745098 | 0.006329322 | brain morphogenesis |
| GO:0050714 | 5.97E-06 | 0.24615385 | 0.006576854 | positive regulation of protein secretion |
| GO:0006939 | 6.65E-06 | 0.28888889 | 0.007316597 | smooth muscle contraction |
| GO:0099092 | 6.77E-06 | 0.47058824 | 0.007443267 | postsynaptic density, intracellular component |

| | | | | |
|---|---|---|---|---|
| GO:0031630 | 6.77E-06 | 0.47058824 | 0.007443267 | regulation of synaptic vesicle fusion to presynaptic active zone membrane |
| GO:0051983 | 6.77E-06 | 0.47058824 | 0.007443267 | regulation of chromosome segregation |
| GO:0071625 | 6.77E-06 | 0.47058824 | 0.007443267 | vocalization behavior |
| GO:0030141 | 6.92E-06 | 0.23287671 | 0.00758786 | secretory granule |
| GO:0004739 | 7.30E-06 | 0.66666667 | 0.007992648 | pyruvate dehydrogenase (acetyl-transferring) activity |
| GO:0035515 | 7.30E-06 | 0.66666667 | 0.007992648 | oxidative RNA demethylase activity |
| GO:0043532 | 7.30E-06 | 0.66666667 | 0.007992648 | angiostatin binding |
| GO:0002576 | 7.30E-06 | 0.66666667 | 0.007992648 | platelet degranulation |
| GO:0006333 | 7.30E-06 | 0.66666667 | 0.007992648 | chromatin assembly or disassembly |
| GO:0032231 | 7.30E-06 | 0.66666667 | 0.007992648 | regulation of actin filament bundle assembly |
| GO:1903748 | 7.30E-06 | 0.66666667 | 0.007992648 | negative regulation of establishment of protein localization to mitochondrion |
| GO:1904714 | 7.31E-06 | 0.40909091 | 0.007992648 | regulation of chaperone-mediated autophagy |
| GO:0075522 | 7.35E-06 | 0.33333333 | 0.007994779 | IRES-dependent viral translational initiation |
| GO:0006633 | 7.59E-06 | 0.22222222 | 0.008246428 | fatty acid biosynthetic process |
| GO:0007219 | 7.98E-06 | 0.21348315 | 0.008661203 | Notch signaling pathway |
| GO:0016072 | 8.46E-06 | 0.53846154 | 0.00917208 | rRNA metabolic process |
| GO:0035984 | 8.46E-06 | 0.53846154 | 0.00917208 | cellular response to trichostatin A |
| GO:0036309 | 8.46E-06 | 0.53846154 | 0.00917208 | protein localization to M-band |
| GO:0042426 | 8.46E-06 | 0.53846154 | 0.00917208 | choline catabolic process |
| GO:0060081 | 8.46E-06 | 0.53846154 | 0.00917208 | membrane hyperpolarization |
| GO:0086015 | 8.46E-06 | 0.53846154 | 0.00917208 | SA node cell action potential |
| GO:0043564 | 8.60E-06 | 0.83333333 | 0.00926904 | Ku70:Ku80 complex |
| GO:0004748 | 8.60E-06 | 0.83333333 | 0.00926904 | ribonucleoside-diphosphate reductase activity, thioredoxin disulfide as acceptor |
| GO:0031812 | 8.60E-06 | 0.83333333 | 0.00926904 | P2Y1 nucleotide receptor binding |
| GO:0046625 | 8.60E-06 | 0.83333333 | 0.00926904 | sphingolipid binding |
| GO:0004852 | 8.60E-06 | 0.83333333 | 0.00926904 | uroporphyrinogen-III synthase activity |
| GO:0006780 | 8.60E-06 | 0.83333333 | 0.00926904 | uroporphyrinogen III biosynthetic process |
| GO:0009386 | 8.60E-06 | 0.83333333 | 0.00926904 | translational attenuation |
| GO:0010124 | 8.60E-06 | 0.83333333 | 0.00926904 | phenylacetate catabolic process |
| GO:0033108 | 8.60E-06 | 0.83333333 | 0.00926904 | mitochondrial respiratory chain complex assembly |
| GO:0060762 | 8.60E-06 | 0.83333333 | 0.00926904 | regulation of branching involved in mammary gland duct morphogenesis |
| GO:0062112 | 8.60E-06 | 0.83333333 | 0.00926904 | fatty acid primary amide biosynthetic process |
| GO:1904292 | 8.60E-06 | 0.83333333 | 0.00926904 | regulation of ERAD pathway |
| GO:1905166 | 8.60E-06 | 0.83333333 | 0.00926904 | negative regulation of lysosomal protein catabolic process |
| GO:0030673 | 9.52E-06 | 0.35714286 | 0.010136498 | axolemma |
| GO:0016079 | 9.52E-06 | 0.35714286 | 0.010136498 | synaptic vesicle exocytosis |
| GO:0016208 | 1.12E-05 | 0.27659574 | 0.011958212 | AMP binding |
| GO:0070531 | 1.13E-05 | 0.39130435 | 0.011975999 | BRCA1-A complex |
| GO:0016624 | 1.13E-05 | 0.39130435 | 0.011975999 | oxidoreductase activity, acting on the aldehyde or oxo group of donors, disulfide as acceptor |
| GO:0032026 | 1.13E-05 | 0.39130435 | 0.011975999 | response to magnesium ion |
| GO:0005758 | 1.30E-05 | 0.21428571 | 0.013740893 | mitochondrial intermembrane space |
| GO:1902494 | 1.36E-05 | 0.34482759 | 0.014423605 | catalytic complex |
| GO:0019887 | 1.36E-05 | 0.34482759 | 0.014423605 | protein kinase regulator activity |
| GO:0098686 | 1.40E-05 | 0.31428571 | 0.014731418 | hippocampal mossy fiber to CA3 synapse |
| GO:0001947 | 1.45E-05 | 0.24193548 | 0.01530219 | heart looping |
| GO:0042060 | 1.56E-05 | 0.17777778 | 0.016423605 | wound healing |
| GO:0050821 | 1.57E-05 | 0.14767932 | 0.016513938 | protein stabilization |
| GO:0005251 | 1.59E-05 | 0.5 | 0.016742181 | delayed rectifier potassium channel activity |
| GO:0030054 | 1.62E-05 | 0.15577889 | 0.016979538 | cell junction |
| GO:0070062 | 1.65E-05 | 0.09348739 | 0.017306362 | extracellular exosome |
| GO:0031091 | 1.69E-05 | 0.375 | 0.017772207 | platelet alpha granule |
| GO:0001659 | 1.69E-05 | 0.375 | 0.017772207 | temperature homeostasis |
| GO:0032059 | 1.72E-05 | 0.6 | 0.017988002 | bleb |
| GO:0044304 | 1.72E-05 | 0.6 | 0.017988002 | main axon |
| GO:1903135 | 1.72E-05 | 0.6 | 0.017988002 | cupric ion binding |
| GO:0042780 | 1.72E-05 | 0.6 | 0.017988002 | tRNA 3'-end processing |
| GO:0004860 | 1.78E-05 | 0.23809524 | 0.018601713 | protein kinase inhibitor activity |
| GO:0097001 | 1.86E-05 | 0.42105263 | 0.019353926 | ceramide binding |
| GO:0005747 | 1.99E-05 | 0.22535211 | 0.020718534 | mitochondrial respiratory chain complex I |
| GO:0031146 | 2.18E-05 | 0.234375 | 0.022696804 | SCF-dependent proteasomal ubiquitin-dependent protein catabolic process |
| GO:0003824 | 2.20E-05 | 0.11952191 | 0.022826206 | catalytic activity |
| GO:0005886 | 2.21E-05 | 0.09145775 | 0.022945226 | plasma membrane |
| GO:0021987 | 2.49E-05 | 0.19791667 | 0.025846612 | cerebral cortex development |
| GO:0006000 | 2.52E-05 | 0.2972973 | 0.026061008 | fructose metabolic process |
| GO:0048787 | 2.81E-05 | 0.46666667 | 0.029046578 | presynaptic active zone membrane |
| GO:0051222 | 2.81E-05 | 0.46666667 | 0.029046578 | positive regulation of protein transport |
| GO:0014704 | 2.85E-05 | 0.27272727 | 0.029314831 | intercalated disc |
| GO:0004565 | 2.84E-05 | 0.71428571 | 0.029314831 | beta-galactosidase activity |
| GO:0010181 | 2.85E-05 | 0.27272727 | 0.029314831 | FMN binding |
| GO:0022904 | 2.85E-05 | 0.27272727 | 0.029314831 | respiratory electron transport chain |
| GO:0000038 | 2.85E-05 | 0.27272727 | 0.029314831 | very long-chain fatty acid metabolic process |
| GO:0001519 | 2.84E-05 | 0.71428571 | 0.029314831 | peptide amidation |
| GO:0015786 | 2.84E-05 | 0.71428571 | 0.029314831 | UDP-glucose transmembrane transport |
| GO:0040040 | 2.84E-05 | 0.71428571 | 0.029314831 | thermosensory behavior |
| GO:0060770 | 2.84E-05 | 0.71428571 | 0.029314831 | negative regulation of epithelial cell proliferation involved in prostate gland development |
| GO:0098910 | 2.84E-05 | 0.71428571 | 0.029314831 | regulation of atrial cardiac muscle cell action potential |
| GO:1903829 | 2.84E-05 | 0.71428571 | 0.029314831 | positive regulation of protein localization |
| GO:0004866 | 2.94E-05 | 0.25490196 | 0.030050775 | endopeptidase inhibitor activity |
| GO:0046726 | 3.56E-05 | 0.54545455 | 0.036336257 | positive regulation by virus of viral protein levels in host cell |
| GO:0005523 | 3.63E-05 | 0.3125 | 0.037023327 | tropomyosin binding |
| GO:0006515 | 3.63E-05 | 0.3125 | 0.037023327 | protein quality control for misfolded or incompletely synthesized proteins |
| GO:2001016 | 3.63E-05 | 0.3125 | 0.037023327 | positive regulation of skeletal muscle cell differentiation |
| GO:0019905 | 3.68E-05 | 0.18691589 | 0.037436086 | syntaxin binding |
| GO:0043005 | 4.14E-05 | 0.13953488 | 0.042014302 | neuron projection |
| GO:0005975 | 4.14E-05 | 0.14112903 | 0.042014302 | carbohydrate metabolic process |
| GO:0007006 | 4.42E-05 | 0.38095238 | 0.044777489 | mitochondrial membrane organization |
| GO:0048143 | 4.42E-05 | 0.38095238 | 0.044777489 | astrocyte activation |
| GO:0071549 | 4.57E-05 | 0.24528302 | 0.046200166 | cellular response to dexamethasone stimulus |
| GO:0006094 | 4.62E-05 | 0.26086957 | 0.04670055 | gluconeogenesis |
| GO:0086091 | 4.62E-05 | 0.26086957 | 0.04670055 | regulation of heart rate by cardiac conduction |
| GO:0001401 | 4.69E-05 | 0.4375 | 0.047348907 | SAM complex |
| GO:0031674 | 4.69E-05 | 0.4375 | 0.047348907 | I band |
| GO:0045792 | 4.69E-05 | 0.4375 | 0.047348907 | negative regulation of cell size |
| GO:0086046 | 4.69E-05 | 0.4375 | 0.047348907 | membrane depolarization during SA node cell action potential |
| GO:2000987 | 4.69E-05 | 0.4375 | 0.047348907 | positive regulation of behavioral fear response |
| GO:0006891 | 4.86E-05 | 0.21052632 | 0.048746278 | intra-Golgi vesicle-mediated transport |
| GO:1903672 | 4.89E-05 | 0.3030303 | 0.049027708 | positive regulation of sprouting angiogenesis |
